# Islands of retroelements are the major components of *Drosophila* centromeres

**DOI:** 10.1101/537357

**Authors:** Ching-Ho Chang, Ankita Chavan, Jason Palladino, Xiaolu Wei, Nuno M. C. Martins, Bryce Santinello, Chin-Chi Chen, Jelena Erceg, Brian J. Beliveau, Chao-Ting Wu, Amanda M. Larracuente, Barbara G. Mellone

**Affiliations:** Department of Biology, University of Rochester; Rochester, NY 14627; Department of Molecular and Cell Biology, University of Connecticut; Storrs, CT 06269; Department of Biomedical Genetics, University of Rochester Medical Center, Rochester, NY 14642; Department of Genetics, Harvard Medical School; Boston, MA 02115; Department of Pathology, Johns Hopkins Medical Institutions, Baltimore, MD, 21287; Wyss Institute for Biologically Inspired Engineering; Harvard Medical School, Boston, MA 02115; Department of Systems Biology, Harvard Medical School; Boston, MA 02115; Department of Genome Sciences, University of Washington; Seattle, WA 98195; Institute for Systems Genomics, University of Connecticut; Storrs, CT 06269

## Abstract

Centromeres are essential chromosomal regions that mediate kinetochore assembly and spindle attachments during cell division. Despite their functional conservation, centromeres are amongst the most rapidly evolving genomic regions and can shape karyotype evolution and speciation across taxa. Although significant progress has been made in identifying centromere-associated proteins, the highly repetitive centromeres of metazoans have been refractory to DNA sequencing and assembly, leaving large gaps in our understanding of their functional organization and evolution. Here, we identify the sequence composition and organization of the centromeres of *Drosophila melanogaster* by combining long-read sequencing, chromatin immunoprecipitation for the centromeric histone CENP-A, and high-resolution chromatin fiber imaging. Contrary to previous models that heralded satellite repeats as the major functional components, we demonstrate that functional centromeres form on islands of complex DNA sequences enriched in retroelements that are flanked by large arrays of satellite repeats. Each centromere displays distinct size and arrangement of its DNA elements but is similar in composition overall. We discover that a specific retroelement, *G2/Jockey-3*, is the most highly enriched sequence in CENP-A chromatin and is the only element shared among all centromeres. *G2/Jockey-3* is also associated with CENP-A in the sister species *Drosophila simulans*, revealing an unexpected conservation despite the reported turnover of centromeric satellite DNA. Our work reveals the DNA sequence identity of the active centromeres of a premier model organism and implicates retroelements as conserved features of centromeric DNA.

## Introduction

Centromeres are marked by the histone H3 variant, CENP-A (also called Cid in *Drosophila*), which is necessary and sufficient for kinetochore activity [1, 2]. Although epigenetic mechanisms play a major role in centromere identity and propagation [3], centromeric DNA sequences can initiate centromere assembly in fission yeast [4] and humans [5], and centromeric transcripts play a role in centromere propagation in human cells [6], suggesting that centromeric DNA-encoded properties may contribute to centromere specification [7]. However, our current understanding of most centromeres remains at the cytological level, as metazoan centromeres are embedded in highly repetitive, satellite-rich pericentric heterochromatin and thus are largely missing from even the most complete genome assemblies. Only recently, long-read single molecule sequencing technologies made it possible to obtain linear assemblies of highly repetitive parts of multicellular genomes such as the human Y chromosome centromere [8] and maize centromere 10 [9].

*Drosophila melanogaster* provides an ideal model to investigate centromere genomic organization as it has a relatively small genome (~180 Mb), organized in just three autosomes (Chr2, Chr3, and Chr4) and two sex chromosomes (X and Y) [10]. The estimated centromere sizes in Drosophila cultured cells range between ~200–500 kb [11] and map to regions within large blocks of tandem repeats [12–15]. While CENP-A associates with simple satellites in ChIP-seq data [16], it may bind to additional undiscovered sequences. The linear organization at the sequence level of any of the centromeres is unknown in this species. Early efforts to determine the structural organization of centromeres in *D. melanogaster* combined deletion analyses and sequencing of an X-derived minichromosome, *Dp1187*. These studies mapped the minimal DNA sequences sufficient for centromere function to a 420-kb region containing the AAGAG and AATAT satellites interspersed with “islands” of complex sequences [14, 15]. However, it is unclear which parts of this minimal region comprise the active centromere, whether or not it corresponds the native X chromosome centromere, and if other centromeres have a similar organization. By and large, satellites have been regarded as the major structural elements of *Drosophila*, humans, and mouse centromeres [2, 3, 17].

In this study, we reveal the detailed organization of all functional centromeres in *D. melanogaster*. By mapping CENP-A on single chromatin fibers at high-resolution, we discover that CENP-A primarily occupies islands of complex DNA enriched in retroelements, which are flanked by large blocks of simple satellites. Our genomic analyses show that all centromeres have a unique sequence organization, even though many of the centromeric elements are shared among them. In particular, all centromeres are enriched for a non-LTR retroelement in the *Jockey* family, *G2/Jockey-3*. While none of these elements are specific to centromeres, they are significantly enriched within these regions. We also find *G2/Jockey-3* enriched at the centromeres of *D. simulans*, which has centromeric satellite arrays highly divergent from those of *D. melanogaster* [16]. Collectively, these data are consistent with the model that retroelements may have a conserved role in centromere specification and function, as proposed for other species (for review see [18]).

## Results

### Identification of candidate centromeres by long-read sequencing and ChIP-seq

To identify the centromeric DNA sequences of *D. melanogaster*, we combined a long-read genome assembly approach [19] with four replicate CENP-A chromatin immunoprecipitations (ChIP) on chromatin from *Drosophila* embryos, followed by paired-end Illumina sequencing (ChIP-seq). We also performed ChIP-seq in *Drosophila melanogaster* Schneider (S2) cells, a widely used model for cell division studies. We took four complementary approaches to discover regions of the genome enriched for CENP-A: *1*) identifying simple repeats enriched for CENP-A based on kmers; *2*) mapping reads to a comprehensive repeat library to summarize enriched transposable elements (TEs) and complex repeats; *3*) using *de novo* assembly methods to assemble contigs from the ChIP reads and calculating enrichment relative to input *post hoc;* and *4*) mapping reads to a heterochromatin-enriched assembly and calling ChIP peaks (Fig. 1A).

**Figure 1.**
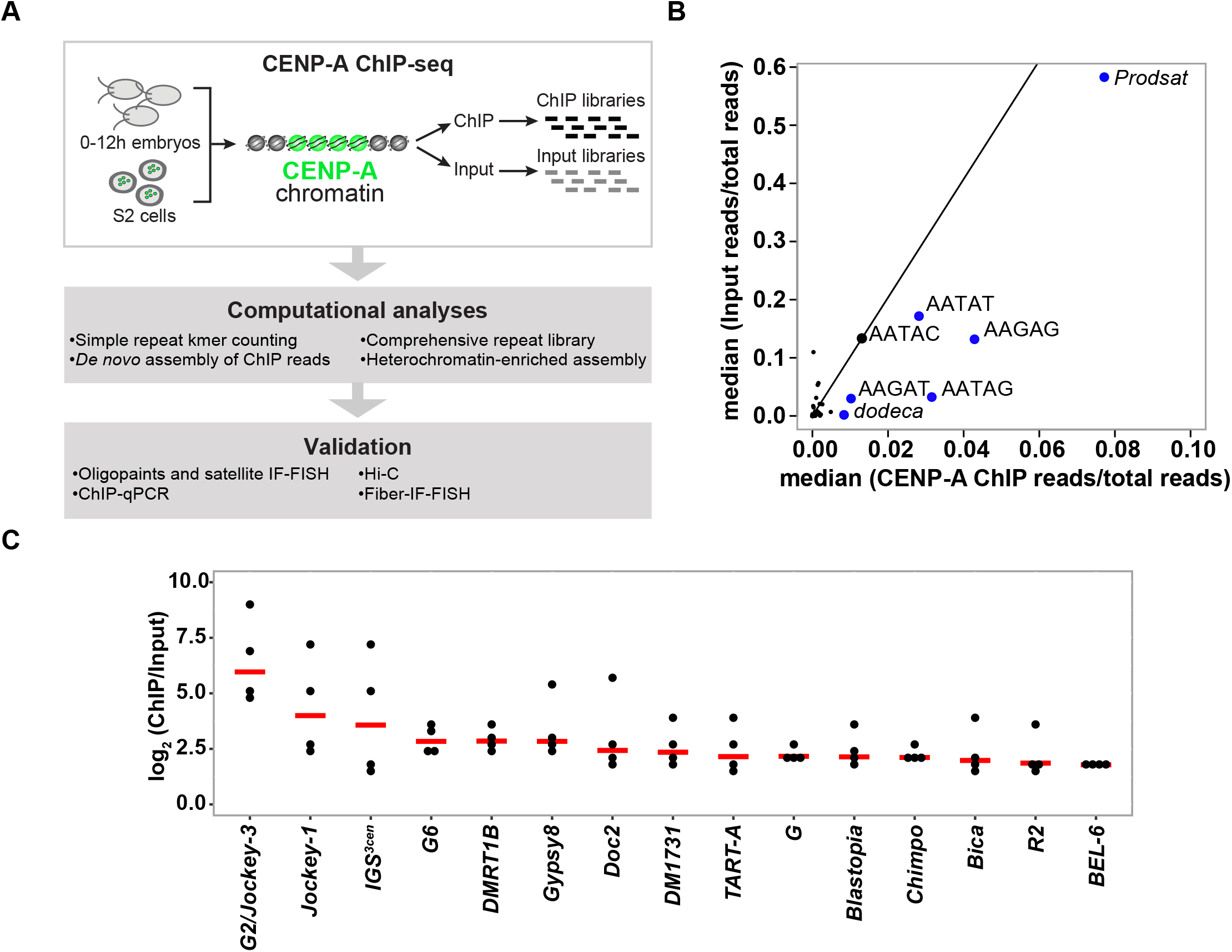
CENP-A binding association with satellites and transposable elements. **A)** Schematic of the strategy used to identify the DNA sequence of *D. melanogaster* centromeres. The Illumina reads are 2×150bp. **B)** Kseek plot showing the relative abundance of simple repeat sequences in CENP-A ChIP compared to the input. Plotted on the x-axis is the median of CENP-A ChIP reads normalized over total mapped CENP-A ChIP reads across four ChIP replicates. Plotted on the y-axis is the median of input reads normalized over total mapped input reads across four replicates. The top 7 kmers in the ChIP read abundance are labeled. The line represents the enrichment of CENP-A ChIP/Input for AATAC, a non-centromeric simple repeat. Repeats to the right of the line are putatively enriched in CENP-A. **C)** Plot of the normalized CENP-A/Input reads on a log scale for each replicate, sorted by median (red lines) for complex repeat families. Shown are only the complex repeats in the top 20% across all four CENP-A ChIP replicates.

In our ChIP experiments, CENP-A pulls down simple satellites, consistent with a previous study [16]. Among the kmers most enriched in CENP-A ChIP relative to input are the *dodeca* satellite and its variants, and complex kmers that include tandem (AATAG)_n_ and (AATAT)_n_ repeats (Fig. 1B; Fig. S1; Table S1). *Prodsat* (also known as the 10bp satellite) is enriched in the CENP-A ChIP, but not relative to input (Fig. 1B). In addition to satellites, we found that CENP-A is also strongly associated with retroelements, particularly non-LTR LINE-like elements in the *Jockey* family and with the Ribosomal Intergenic Spacer (IGS). Among the *Jockey* elements the most highly enriched in CENP-A ChIPs are annotated as *G2* and *Jockey-3* (Fig. 1C; Table S2). However, our phylogenetic analysis suggests that these repeats correspond to the same type of element, as genomic copies of *G2* and *Jockey-3* are interleaved across the tree and not monophyletic (Fig. S2). Thus, we hereafter collectively refer to these elements as *G2/Jockey-3*.

To detect CENP-A-enriched sequences independently of known repeats in repeat libraries or of genome assemblies, we *de novo* assembled CENP-A ChIP reads into contigs (*i.e*. ChIPtigs [20]) and calculated their CENP-A enrichments. The resulting CENP-A enriched ChIPtigs primarily contained fragments of TEs and other complex repeats, and some simple satellite repeats (Table S3).

To determine the genomic location of CENP-A enriched sequences, we mapped ChIP reads to a new reference genome assembly that we generated using a heterochromatin-enriched assembly method resulting in greater representation of heterochromatin-associated regions [19] (Table S4, and supplemental results). Five contigs were consistently the most CENP-A enriched in the assembly, with highly reproducible ChIP peaks across technical and biological replicates (IDR < 0.05; Table S5; Fig. S3). These CENP-A-enriched contigs have a similar organization: they contain islands of complex DNA (*e.g*. TEs) flanked by simple tandem satellite repeats with known centromeric locations (Fig. 2; Fig. S4; Table 1). The candidate centromeric contig for the X chromosome (Contig79) is 70-kb and contains a 44-kb island of complex DNA (called *Maupiti* [15]), flanked by a short stretch of AAGAT satellite on one side and embedded in AAGAG satellite (Fig. 2A). This region has an organization that is nearly identical to that of the *Dp1187* minichromosome putative centromere [14, 15], suggesting that this contig may contain at least part of the endogenous X centromere. The candidate centromeric contig for chromosome 4 (Contig119) contains a 42.8-kb island (we named *Lampedusa*) flanked by the AAGAT satellite (Fig. 2B). A recent study mapped the AAGAT satellite cytologically to centromeres of chromosome 4 and a B chromosome derived from this chromosome [21], consistent with *Lampedusa* being a candidate for centromere 4. The candidate centromeric contig for chromosome Y (Y_Contig26) consists of a 138-kb island (we named *Lipari;* Fig. 2C). The candidate centromeric contig for chromosome 3 (Contig 3R_5) contains a 68.5-kb island (we named *Giglio*) flanked by *Prodsat* and the *dodeca* satellite, which map to this centromere cytologically [12, 22, 23] (Fig. 2D). Finally, the candidate contig for chromosome 2 (tig00057289) contains a small 1.8-kb complex island (we named *Capri*), flanked by the AATAG and AAGAG satellites (Fig. 2E). The majority of the top enriched *de novo* ChIPtigs (88/100 for R1; 19/30 for R2; 26/30 for R3; and 82/100 for R4) map uniquely to these five contigs (Table S3), providing independent support for the assembly and further substantiating our hypothesis that these contigs correspond to the centromeres.

**Figure 2.**
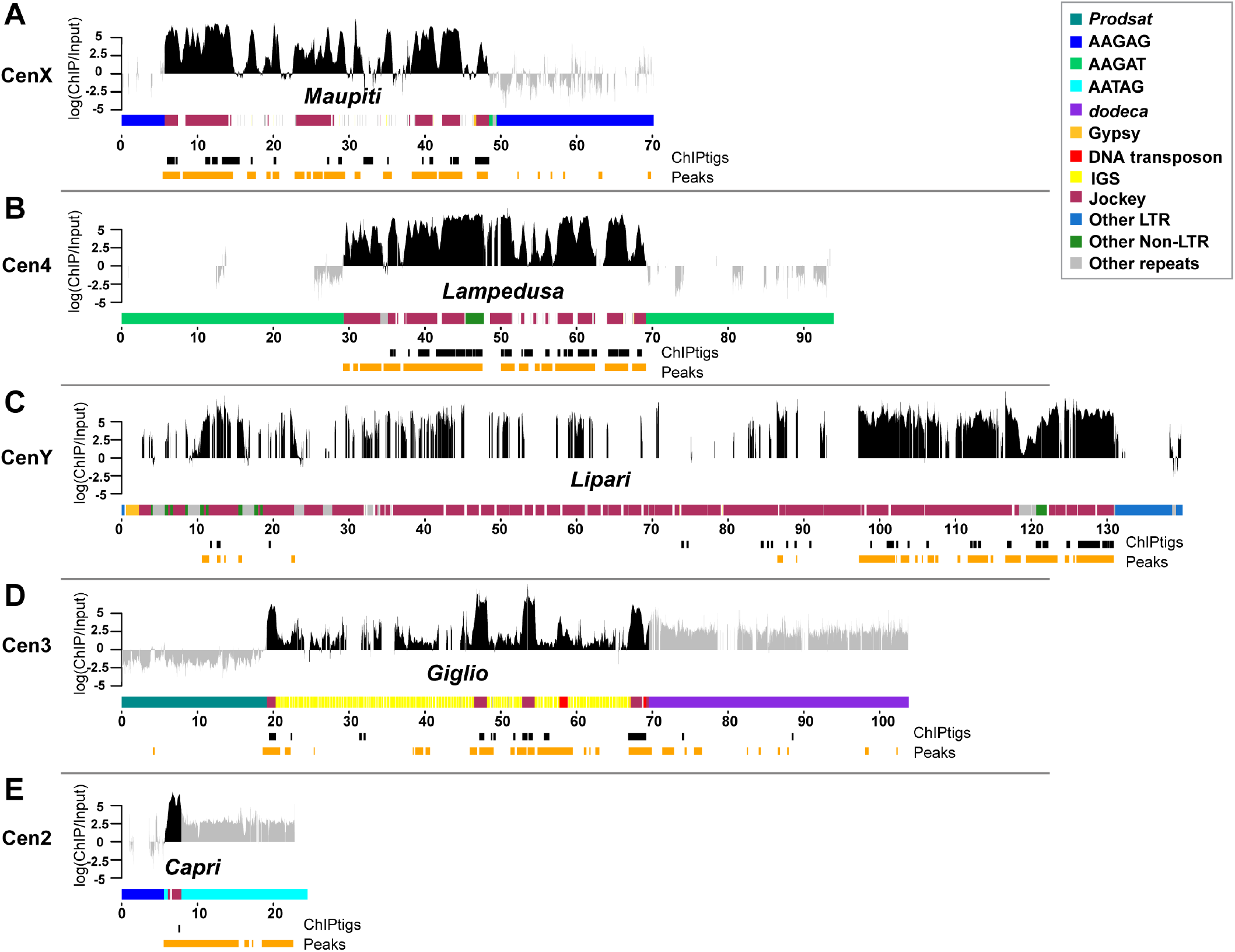
CENP-A occupies DNA sequences within putative centromere contigs. Organization of each CENP-A enriched island corresponding to centromere candidates: **A)** X centromere, **B)** centromere 4; **C)** Y centromere; **D)** centromere 3; **E)** centromere 2. Different repeat families are color coded (see legend; note that *Jockey* elements are shown in one color even though they are distinct elements). Shown are the normalized CENP-A enrichment over input (plotted on a log scale) from one replicate (Replicate 2, other replicates are in Fig. S4) colored in gray for simple repeats and black for complex island sequences. While the mapping quality scores are high in simple repeat regions, we do not use these data to make inferences about CENP-A distribution (see text for details). The coordinates of the significantly CENP-A-enriched ChIPtigs mapped to these contigs (black) and the predicted ChIP peaks (orange) are shown below each plot.

**Table 1:** Location of centromeric and centromere-proximal satellites in *Drosophila melanogaster*. Locations of satellite on chromosomes X, Y, 2, 3, and 4 according to previous reports as well our observations in this report by IF-FISH in the *D. melanogaster* sequenced strain iso-1. Each satellite location is characterized as being centromeric (Cen, overlaps with CENP-C), pericentric (Peri, juxtaposed to CENP-C) or heterochromatic (Het, more distal than pericentric). Note that the *dodeca* satellite includes its variant and *Prodsat* is also known as the 10bp satellite. *Indicates a small block not easily detected by FISH. ^a^ Lohe et al., 1993 [66]; Jagannathan et al. 2017 [37] ^b^ Talbert et al., 2018 [16] ^c^ Sun et al., 2003 [14] ^d^ Tolchov et al., 2000 [67] ^e^ Hanlon et al. 2018 [21] ^f^ Abad et al., 1992 [63]; Garavís et al., 2015 [12]; Jagannathan et al., 2017 [37] ^g^ Torok et al., 1997,2000 [68, 69]; Blower and Karpen, 2001 [70]; Garavis et al., 2015 [12]

|  |  | Previous Reports |  |  | This study |  |  |
| --- | --- | --- | --- | --- | --- | --- | --- |
| Satellite | Sequence | Cen | Peri | Het | Cen | Peri | Het |
| AATAT<sup>a,b,c,d</sup> | (AATAT)<sub>n</sub> | X | - | 3,4,Y | X | Y | 3,4,Y |
| AAGAG<sup>a,b,c</sup> | (AAGAG)<sub>n</sub> | X | - | 2,3,4,Y | 2,X | 4 | 3,Y |
| AATAG<sup>a,b</sup> | (AATAG)<sub>n</sub> | - | - | 2,Y | 2* | 3 | 2,Y |
| AAGAT<sup>e</sup> | (AAGAT)<sub>n</sub> | 4 | - | - | 4 | X | 2 |
| *dodeca*<sup>f</sup> | (CGGTCCCGTACT/<br>GGTCCCGTACT)<sub>n</sub> | 3 | - | - | 3 | - | - |
| *Prodsat*<sup>a,g</sup> | (AATAACATAG)<sub>n</sub> | - | 2,3 | - | 2 | 2,3 | - |

### Genomic distribution of CENP-A in embryos and S2 cells

Our ChIP-seq experiments and their analyses provide evidence that CENP-A is specifically associated with the island DNA sequences for Contig79 (X^*Maupiti*^), Contig119 (4^*Lampedusa*^), Y_Contig26 (Y^*Lipari*^), 3R_5 (3^*Giglio*^), and with a single interspersed *G2/Jockey-3* fragment within tig00057289 (2^*Capri*^; Fig. 2; Fig. S4). A previous study that used a *D. melanogaster* native ChIP-seq dataset (using anti-GFP antibodies and CENP-A-GFP expressing embryos) focused exclusively on the quantification of simple repeats and did not identify any complex DNA associated with CENP-A [16]. However, our re-analysis of this dataset showed association of CENP-A-GFP with the centromere islands (Fig. S3B; Tables S4-S6). We validated individual elements for which we could design contig-specific qPCR primers in additional independent CENP-A ChIP experiments and confirmed that the CENP-A peaks in these regions are not a result of library amplification bias [24] (Fig. S5; Table S7).

Having shown that CENP-A is associated with the complex islands, we next analyzed if the centromere extends to the surrounding satellite DNA. Simple sequences flanking the islands appear among the kmers enriched in the CENP-A ChIP (Fig. 1B; Table S1; Fig. S1). However, it is difficult to quantify the enrichment of CENP-A on simple satellite repeats for several reasons: *1*) simple satellite sequences may be over or underrepresented as an artifact of library preparation [24], particularly for ChIPseq experiments that rely on PCR amplification to construct libraries; *2*) satellites are abundant genomic sequences that are largely missing from whole genome assemblies [10], making it difficult to precisely quantitate how much of these sequences exist in genomes (and therefore how much to expect in the input); *3*) highly abundant repeats are expected to have a low signal-to-noise ratio if a relatively small fraction of a simple repeat is enriched in CENP-A relative to the overall abundance of this satellite in the genome; and *4*) simple satellite repeats present a challenge for even long-read based genome assembly methods [25]. While we may be confident in large-scale structural features of assemblies involving highly repetitive sequences—we see even PacBio read depth in islands, but uneven depth on simple satellites (Fig. S6). Due to these limitations, we caution against using strictly assembly-based approaches in regions with simple repeats. Nonetheless, we report the ChIP peaks on simple satellites (shaded in gray in Fig. 2). To confirm satellite localization near each centromere, we employed IF with anti-CENP-C (an inner kinetochore protein that co-localizes with CENP-A), followed by FISH with probes for the satellites *dodeca*, AAGAG, AATAT, AAGAT, AATAG, and *Prodsat* on metaphase chromosome spreads from 3^rd^ instar larval brains (Fig. S7); a summary of the co-localization data is shown in Table 1.

Although CENP-A localizes exclusively to the centromeres at the cytological level, it is possible that low-levels of CENP-A occupy non-centromeric DNA. We found a low, but consistent CENP-A enrichment at genomic regions outside of the centromere islands, including some telomere-associated elements (e.g. *TART-A*), rDNA genes from the rDNA clusters, and the LINE-like retroelements DMRT1B and R2 (Fig. 1C; Table S8 and supplemental results). Many of these associations likely represent non-specific peaks [26], as they were not highly enriched by CENP-A ChIP-qPCR (Fig. S5). However, previous studies found evidence for an association of some centromeric proteins with the nucleolus [27], perhaps relating to the possible association that we detect for rDNA and rDNA-associated retroelements (*e.g*. R2). We also noted that non-centromeric copies of *G2/Jockey-3* were not consistently enriched in CENP-A (Table S8).

CENP-A ChIP-seq reads from S2 cells showed a similar enrichment profile of sequences represented in the embryo ChIP-seq data (*e.g*. IGS and *G2/Jockey-3*) but were much more enriched for additional retroelements that were not represented within our centromere contigs (*e.g*. LTR elements *Dm1731, HMSBeagle*, and *Max-I;* Table S2). We also observed a similar pattern of CENP-A enrichment on simple satellite repeats in S2 cells (AATAT, AATAG, AAGAG, *Prodsat*, and *dodeca;* Table S1) and we confirmed that these satellites are also near centromeres cytologically using IF/FISH in S2 cells (Fig. S8). However, complex satellites that are pericentric in embryos, including complex satellites in the 1.688 family and *Rsp* (Table S2), are CENP-A-enriched in S2 cells. This suggests that the centromeres of S2 cells may have expanded into regions that are pericentromeric in flies; the additional retroelements enriched in CENP-A may be pericentric or they may represent new retroelements insertions occurred in this cell line. Our findings are consistent with the extensive structural rearrangements and polyploidy reported for these cells [28].

### Centromeres are unique, but are composed of similar non-LTR retrotransposons

Although each island has a distinct arrangement of AT-rich sequences, repeats, and TEs, their compositions are overall similar. In particular, non-LTR retroelements in the *Jockey* family such as *G2/Jockey-3, Doc*, and *Doc-2* are especially abundant within CenX, Cen4, and CenY (Fig. 2; Fig. 3A). Strikingly, *G2/Jockey-3* is the only element present in all five of our centromere contigs, suggesting a potential role in centromere function or specification. In our phylogenetic analysis of genomic *G2/Jockey-3* repeats in *D. melanogaster*, we cannot distinguish *G2/Jockey-* 3 elements at centromeres from those across the genome, suggesting that centromeric TEs do not have a single origin (Fig. 3B and supplemental results). Although *G2/Jockey-3* is not unique to centromeres, and thus cannot be sufficient for centromere identity, it is significantly enriched at centromeres: ~63% of all genomic copies of *G2/Jockey-3* are found within our candidate centromere contigs (Fig. 4 and Table S9). *G2/Jockey-3* elements show signs of recent or ongoing activity based on their insertion polymorphism [29], pattern of 5’ truncation (see supplemental results), and expression (Fig. S9A). At least some of this expression comes from the centromeres: we analyzed total embryo RNA extracts by reverse-transcription qPCR (RT-qPCR) using primers targeting centromere-associated copies and found evidence for low-levels of *G2/Jockey-3* transcription from copies in the X, 4 and 3 centromeres, with no or negligible expression from the Y and 2 (Fig. S9B).

**Figure 3.**
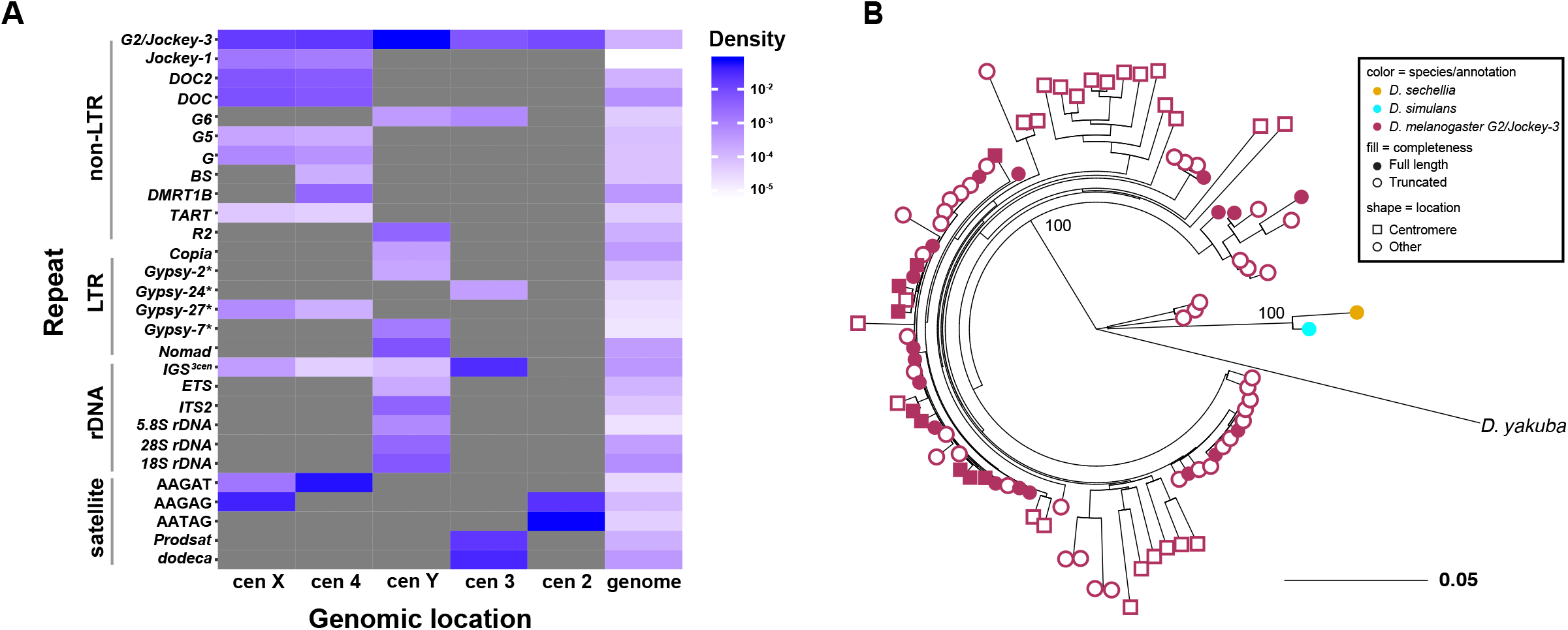
Centromeres are enriched in non-LTR retroelements in the *Jockey* family. **A)** Density of all repetitive elements on each candidate centromere contig and the entire genome (minus the centromeres) grouped by type: non-LTR retroelements, LTR retroelements, rDNA-related sequences, and simple satellites. *G2/Jockey-3* is present on all centromeres. **B)** Maximum likelihood phylogenetic tree based on the entire sequence of all *G2/Jockey-3* copies in *D. melanogaster* inside (squares) and outside (circles) of centromeric contigs, and on the consensus repeat in its sister species *D. sechellia* and *D. simulans*, and a more distantly related species (*D. yakuba*). The tree shows that centromeric *G2/Jockey-3* elements do not have a single origin.

**Figure 4.**
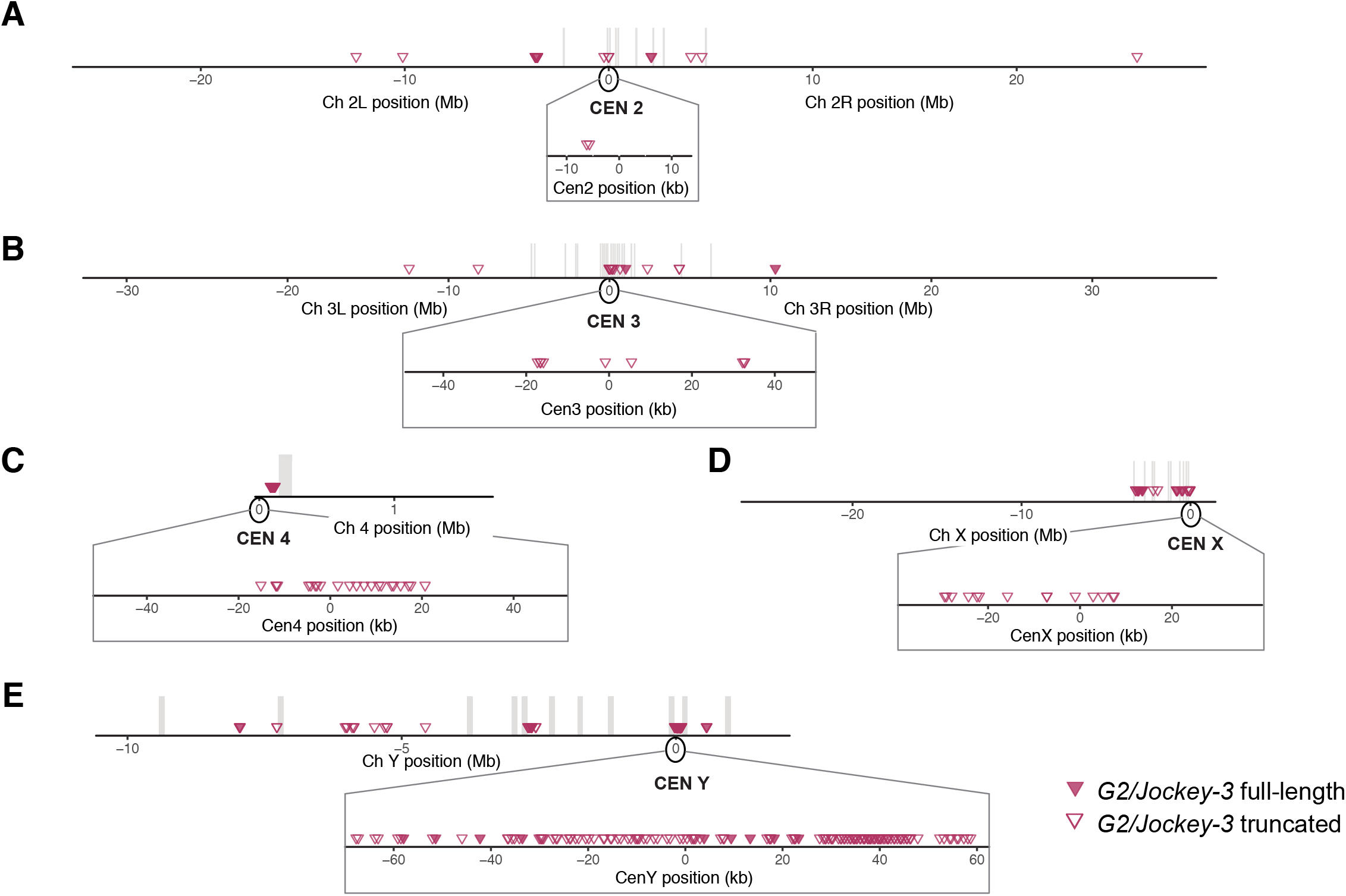
Genomic distribution of *G2/Jockey-3* elements in the *D. melanogaster* genome. Location of *G2/Jockey-3* elements across chromosome 2 (A), 3 (B), 4 (C), X (D), and Y (E). Contigs from each chromosome were concatenated in order with an arbitrary insertion of 100 kb of ‘N’. Distances along the x-axis are approximate. The order and orientation of the Y chromosome contigs is based on gene order (see [19]). Each triangle corresponds to one TE, where filled shapes indicate full length TEs and open shapes indicate truncated TEs. The vertical gray bars represent the arbitrary 100 kb window inserted between contigs, indicating where there are gaps in our assembly. The centromere positions are set to 0 for each chromosome. The insets zoom in to show the distribution of *G2/Jockey-3* elements on the centromere contigs. Chromosomes are not drawn to scale (chromosome 4 and Y are enlarged).

In addition to *G2/Jockey-3*, the 3^*Giglio*^ island has 240 copies of a centromere-enriched variant of the ribosomal IGS (supplemental results; Fig. S10). Among the islands, 2^*Capri*^ differs the most, being the smallest and harboring only a single fragment of *G2/Jockey-3* (Fig. 2E). Importantly, as was previously reported for the X-derived *Dp1187* centromere [14, 15], none of the sequences contained within these islands are exclusive to centromeres. However, several of these elements are enriched in these regions compared to the genome in addition to *G2/Jockey-3*. For example, *DOC2, G*, and *Jockey-1* elements are non-LTR retroelements enriched in CENP-A with a genomic distribution biased toward centromeres (Fig. 3A, columns labeled “genome”; Fig. S11; Table S9).

### Validation of centromeric contigs

To verify the association of our contigs with the centromeres, we performed IF with anti-CENP-C antibodies, followed by FISH with satellite probes and custom-designed Oligopaints libraries [30](see methods) for X^*Maupiti*^, 4^*Lampedusa*^, Y^*Lipari*^, and 3^*Giglio*^ (Fig. 5 and Fig. S12). The X^*Maupiti*^ Oligopaints hybridized to the X as well as the Y centromeres on 3^rd^ instar male larval brain metaphase spreads (Fig. 5A and Fig. S12A). Similarly, the Oligopaints for 4^*Lampedusa*^ hybridized to the 4^th^ as well as to the Y centromere (Fig. 5B and S12B), suggesting that Oligopaints for X^*Maupiti*^ and 4^*Lampedusa*^ have homology to sequences at or near the Y centromere. In contrast, the Oligopaints for Y^*Lipari*^ (Fig. 5C and Fig. S12C) and 3^*Giglio*^ were specific for their respective centromeres (Fig. 5D and Fig. S12D). We could not use Oligopaints to validate 2^*Capri*^ because of its small size, but its organization, with AATAG and AAGAG satellites flanking a small CENP-A enriched island (Fig. 2E), is consistent with our FISH analyses (Fig. 5E). In line with the CENP-A ChIP-seq data, we observed significant differences between S2 and embryo centromeres by Oligopaint FISH. With the exception of 3^*Giglio*^, centromeric island organization in S2 cells is dramatically different from larval brain metaphase spreads (Fig. S13 and supplemental results), in contrast to the conservation of the centromeric distribution of simple satellites (Fig. S8; Table S1).

**Figure 5.**
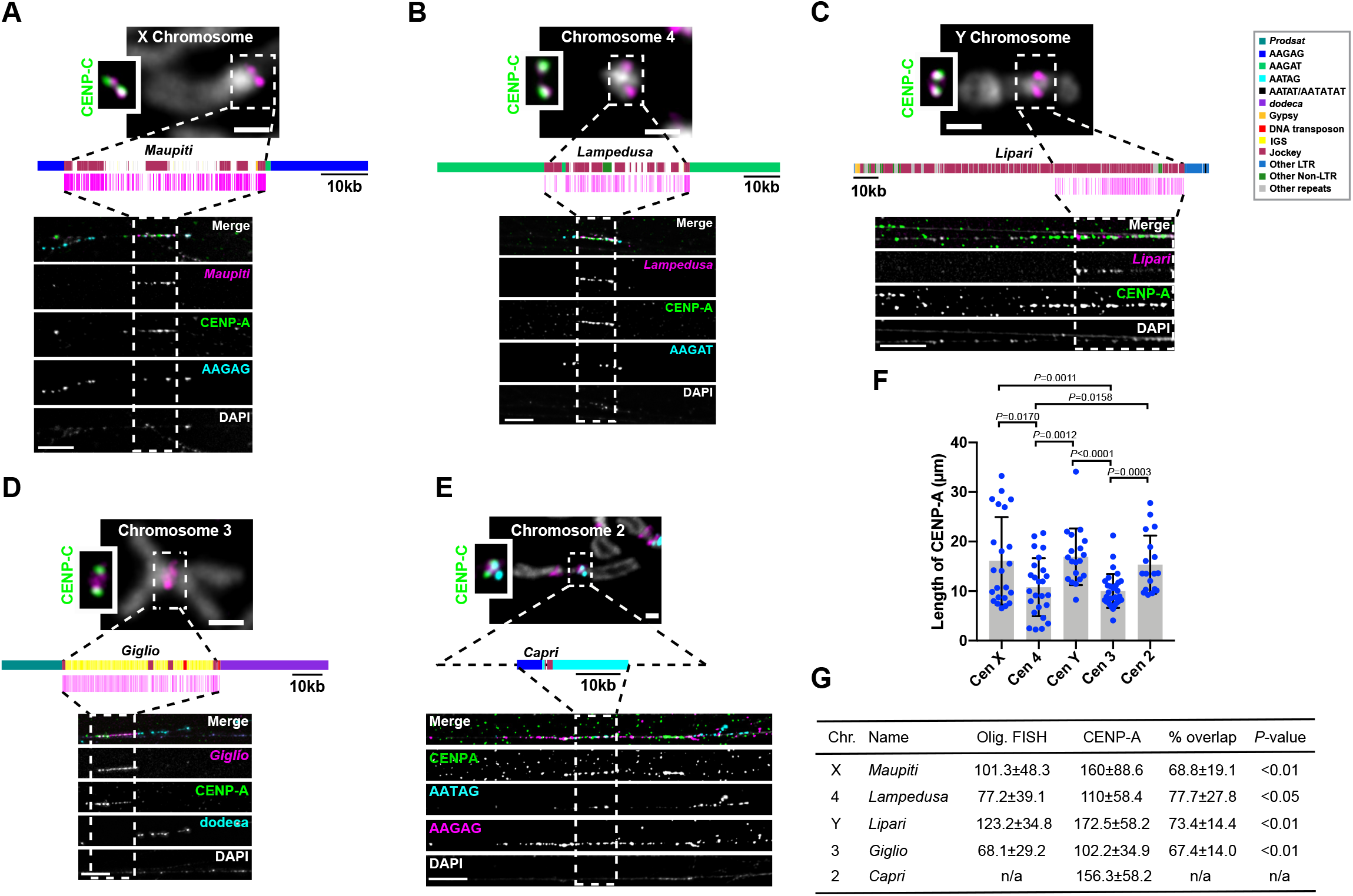
Islands of complex DNA are major components of centromeres. **A-D)** Top, mitotic chromosomes from male larval brains showing IF with anti-CENP-C antibodies (green, inset) and FISH with chromosome-specific Oligopaints (magenta). Bar 1 μm. Middle, schematic of centromere contigs (see key) and location of Oligopaint targets (magenta). Bottom, IF-FISH on extended chromatin fibers from female larval brains. Anti-CENP-A antibodies (green), Oligopaints FISH (in panels A, B, and D; magenta) and a centromere-specific satellite (cyan, and in E also in magenta). Dashed rectangles show the span of the Oligopaint probes, except for E, where it is placed arbitrarily within the CENP-A domain where the cen 2 contig could be located. Bar 5μm. **A)** X centromere; **B)** Centromere 4; **C)** Y centromere; **D)** Centromere 3 (see also Fig. S20); **E)** Centromere 2 using FISH probes AAGAG (magenta) and AATAG (cyan). The scale shown for the centromere 2 diagram is approximate. **F)** Scatter plot of CENP-A IF signal length for each centromere. Error bars=SD. n=18-30 fibers for each centromere. Significant P-values are shown (unpaired T-test). **G)** Table showing the lengths of Oligopaint FISH and CENP-A IF signals on fibers (kb±SD estimated based on 10μm=100kb, Fig. S15). % overlap corresponds to CENP-A domain length/Oligopaint FISH length. The difference between the sizes of the CENP-A domain and the corresponding islands is significant (unpaired t-test). n/a = not applicable. Additional fibers are shown in Figs. S16-21.

*Drosophila* centromeres tend to cluster in the nucleus cytologically [31, 32]. We found independent support for the complex islands being centromeric by analyzing previously published Hi-C data from *D. melanogaster* embryos. Island-island interactions were among the most frequent inter-chromosome interactions, followed by interactions between islands and their own proximal pericentric heterochromatin, and lastly by interactions between islands and distal pericentric heterochromatin or euchromatin (Fig. S14; supplemental results). This analysis also shows that indeed native centromeres interact with one another physically in the 3D nucleus.

### Analysis of extended chromatin fibers reveals that CENP-A primarily occupies the islands

Based on the enrichment of CENP-A with island-associated repeats, we hypothesized that the TE-enriched islands are the major centromere components in *D. melanogaster*. To test this, we investigated CENP-A occupancy, a direct reflection of centromere activity, and estimated the size of each centromere by visualizing extended chromatin fibers [11, 33]. This method has two major advantages: it does not rely on mapping low complexity ChIP reads, thus providing more information that can be inferred by ChIP, and it affords single-chromosome, rather than population, information on CENP-A localization. We carried out IF with CENP-A antibodies and FISH with Oligopaint and satellite probes on cells from 3^rd^ instar larval brains, selecting females to ensure specificity for our X^*Maupiti*^ and 4^*Lampedusa*^ Oligopaints (Fig. 5). First, we calibrated our fiber stretching using three FISH probes spanning 100kb: two heterochromatic (one for the *Rsp* locus, Table S11 [34], and one Oligopaint targeting the pericentromere of chromosome 3L, see methods for coordinates) and one euchromatic (an Oligopaint targeting a region ~600kb from the telomere of chromosome 3L, see methods). The estimated stretching for these fibers is ~10 kb/μm for all three locations, with no significant difference among them (*P*=0.085; Fig. S15). We next determined the sizes of the CENP-A domain and corresponding island of each centromere (Fig. 5 and Figs. S16–21). The size of the CENP-A domain varies between centromeres, ranging in mean size between 101–172-kb (~11μm-17μm), smaller than previous estimates that relied on the measuring of a mixture of centromeres in *Drosophila* Kc and S2 cells [11]. This is consistent with our ChIP-seq analysis suggesting that S2 cells may have expanded centromeres. X, Y and 2 are the largest centromeres, while 3 and 4 are the smallest (Fig. 5F–G). Importantly, CENP-A primarily occupies the centromeric islands X^*Maupiti*^, 4^*Lampedusa*^, Y^*Lipari*^ and 3^*Giglio*^ (~70% of the CENP-A domain overlaps with the Oligopaint FISH signal; Fig. 5G and Figs. S16-21). In some fibers, the X^*Maupiti*^ Oligopaint FISH signal showed interspersion with FISH signal for the AAGAG satellite (Fig. S16), this could be due to non-specific binding of the AAGAG probe during FISH, which is optimized for Oligopaint specificity, or because AAGAG repeats are collapsed in our assembly, including within *Maupiti*. We also noticed that the estimated length of the Oligopaint-stained region was larger than the size of *Maupiti* in our CenX contig (101.3 ±48.3-kb versus 44-kb; Fig. 2A and Fig. 5G), a discrepancy that we attribute to variability in *Maupiti* Oligopaint probe hybridization. Alternatively, there could be additional sequences with similarity to *Maupiti* interspersed in the flanking satellites nearby the contig (and not in our assembly).

Analysis of centromere 4 shows that the CENP-A domain overlaps primarily with 4^*Lampedusa*^, and partially with the flanking AAGAT satellite (Fig. 5B, F and Fig. S17). The Oligopaints for Y^*Lipari*^ target only the part of the island with the highest enrichment of CENP-A (Fig. 5C). Fibers for this centromere show a continuous CENP-A domain that extends past the FISH signal, likely representing the remainder of the Y^*Lipari*^ island (Fig. 5C and Fig. S18).

Fibers for 3^*Giglio*^ show co-localization between CENP-A and the island as well as a short, variable region of co-localization with flanking *dodeca* satellite (Figure 5D, Fig. S19–20). We did not observe CENP-A signal on the opposite side of *Giglio*, where *Prodsat* is located according to our assembly (Fig. 5D). The centromere 3 satellite *dodeca* co-localizes with CENP-A on fibers in S2 cells [12] and is highly enriched in our CENP-A ChIP-seq (Fig. 1B; Fig. S1). When we tracked longer fibers from 3^*Giglio*^ along *dodeca*, we observed a second CENP-A domain where *dodeca* is interrupted by short fragments of Oligopaint FISH signal (Fig. S20), suggesting the existence of DNA sequences with homology to *Giglio* interspersed within *dodeca* that are not included in our assembly. A previous study identified sequences with homology to IGS within the *dodeca* satellite in one BAC [12]. It is possible that the *dodeca*-associated Oligopaint FISH signal in our extended fibers corresponds to these additional IGS sequences. These data indicate that centromere 3 has two CENP-A domains, a major one on 3^*Giglio*^ and one minor one on *dodeca*, although these appear as a single domain in standard metaphase spread IF. Unlike centromere 3, all other centromeres display a single CENP-A domain by fiber analysis (see Fig. S21 for centromere 2 and data not shown). Our conclusions differ from the Talbert et al. study [16], which concluded that *dodeca* was not associated with CENP-A. As recognized by the authors, it is possible that different chromatin preparations, such as the MNase digestion, may introduce biases, leading to an underrepresentation of sequences like *dodeca* in ChIPs [16]. Lastly, we analyzed the organization of 2^*Capri*^ using FISH with a satellite combination unique to this chromosome: AATAG, AAGAG, and *Prodsat* and found that the CENP-A domain overlapped with all three satellites (Fig. 5E and Fig. S21). Thus, we speculate that the *Prodsat* sequences pulled down by CENP-A as seen in our kmer analysis (Fig. 1B) and reported previously [16] are coming from the centromere 2, not 3. We therefore conclude that *D. melanogaster* CENP-A is primarily associated with the centromeric islands of chromosomes X, 4, Y, and 3, and less predominantly with the flanking satellites (Fig. 5G).

### *G2/Jockey-3* is centromere-associated in *Drosophila simulans*

The *G2/Jockey-3* retroelement is a recently active transposon [29] shared amongst all *D. melanogaster* centromeres (Fig. 3A; Table S2). To determine if *G2/Jockey-3* is enriched at the centromeres outside of *D. melanogaster*, we investigated its centromeric distribution in the sister species, *D. simulans*, which diverged from *D. melanogaster* only ~2 million years ago [35] and yet displays major differences in satellite composition and distribution [36, 37]. These differences are especially apparent in centromeric regions, where *D. melanogaster* displays simple satellite repeats while *D. simulans* contains complex satellite repeats with larger repeat units [16]. We reanalyzed published *D. simulans* cell line CENP-A ChIP-seq data [16] (see supplemental results) and found that *G2/Jockey-3* elements are also highly enriched in CENP-A in this species, similar to *D. melanogaster*. The pileup of CENP-A ChIP reads on *G2/Jockey-3* show that CENP-A is associated with the entire length of the retroelement in both *D. simulans* and *D. melanogaster*, with no apparent affinity for any particular sequence (Fig. 6A–B).

**Figure 6.**
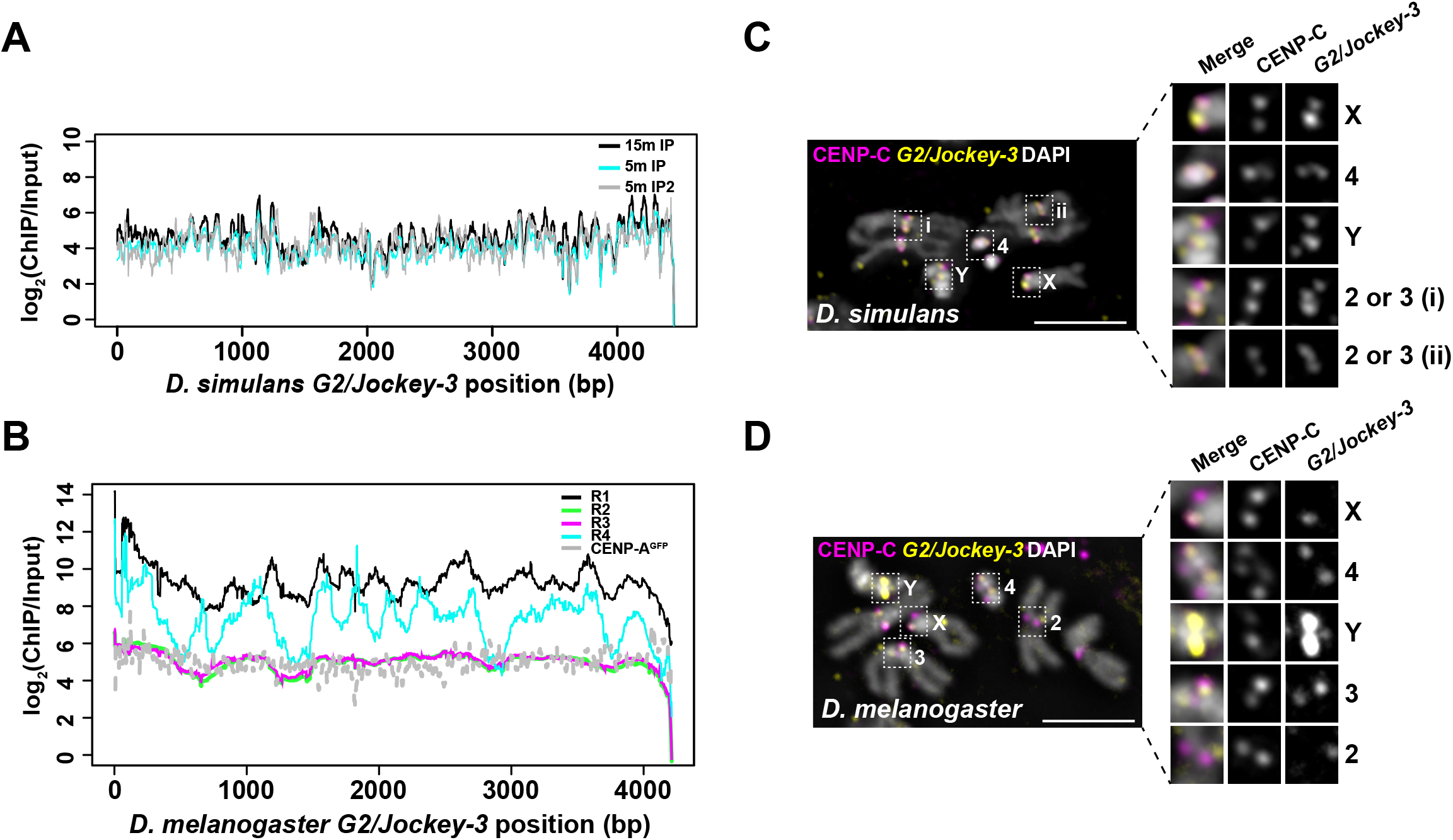
The association between *G2/Jockey-3* and centromeres is conserved in *D. simulans*. **A)** Plot of the normalized CENP-A enrichment over input across the *D. simulans G2/Jockey-3* consensus sequence using CENP-A ChIP-seq data from *D. simulans* ML82-19a cells (Talbert *et al*. 2018) showing that *G2/Jockey-3* is enriched in CENP-A in *D. simulans*. 15m and 5m indicate duration of MNase digestion and IP and IP2 are technical replicates. Note that the first 487bp of *D. simulans G2/Jockey-3* consensus sequence, which are homologous to the 500bp satellite, are not included in this figure; the 500bp satellite was previously reported as enriched in CENP-A in *D. simulans* [16]. **B)** Plot of the normalized CeNp-A enrichment over input across the *D. melanogaster G2/Jockey-3* consensus sequence using our CENP-A ChIP-seq replicates (R1-R4) and ChIP-seq from CENP-A-GFP transgenic flies from Talbert *et al*. 2018. IF-FISH on **C)** *D. simulans* (w501) and **D)** *D. melanogaster* (iso-1) mitotic chromosomes from male larval brains using an antibody for CENP-C (magenta) and FISH with a *G2/Jockey-3* Digoxigenin labeled FISH probe (yellow). DAPI is shown in gray. Bar 5μm.

To validate the association of *G2/Jockey-3* with *D. simulans* centromeres, we designed a FISH probe that targets ~1.6 kb at the 3’ of the *D. melanogaster G2/Jockey-3* consensus sequence (see methods; ~94% identical to *D. simulans G2/Jockey-3* consensus sequence) and performed IF/FISH on male larval brain metaphase spreads with anti-CENP-C antibodies, which recognize CENP-A in both species [38]. We observed co-localization between CENP-C and *G2/Jockey-3* at all *D. simulans* centromeres (Fig. 6C; note that chromosome 2 and 3 of *D. simulans* cannot be distinguished morphologically [37]). The same probe showed co-localization of CENP-C and *G2/Jockey-3* at all *D. melanogaster* centromeres, except at centromere 2, which is consistent with our model for this centromere showing only one copy of *G2/Jockey-3* (Fig. 6D and Fig. 2E). Based on these observations, we propose that *G2/Jockey-3* is a conserved centromere-associated retroelement in these species.

## Discussion

Our study shows that combining long-read sequencing with ChIP-seq and chromatin fiber FISH is a powerful approach to discover centromeric DNA sequences and their organization. We reveal that, for all but one chromosome (chromosome 2, which has a single *G2/Jockey-3* element), ~70% of the functional centromeric DNA of *D. melanogaster* is composed of complex DNA islands. The islands are rich in non-LTR retroelements and are buried within large blocks of tandem repeats (Fig. 7A). They likely went undetected in previous studies of centromere organization (*e.g*. [12]) because three of the five islands are either missing or are incomplete in the published reference *D. melanogaster* genome [10]. A recent study reported that satellite DNA repeats comprise the majority of centromeric DNA in *D. melanogaster* embryos and S2 cells, by counting the relative number of motifs matching simple repeats in CENP-A ChIP relative to input [16]. Our re-analysis of those data showed CENP-A enrichment on the islands, suggesting that having an improved reference genome assembly [19] is crucial for identifying centromeric DNA. To our knowledge, this is the first detailed report on the linear sequence of all centromeres in a multicellular organism. Our overall strategy therefore provides a blueprint for determining the composition and organization of centromeric DNA in other species.

**Figure 7.**
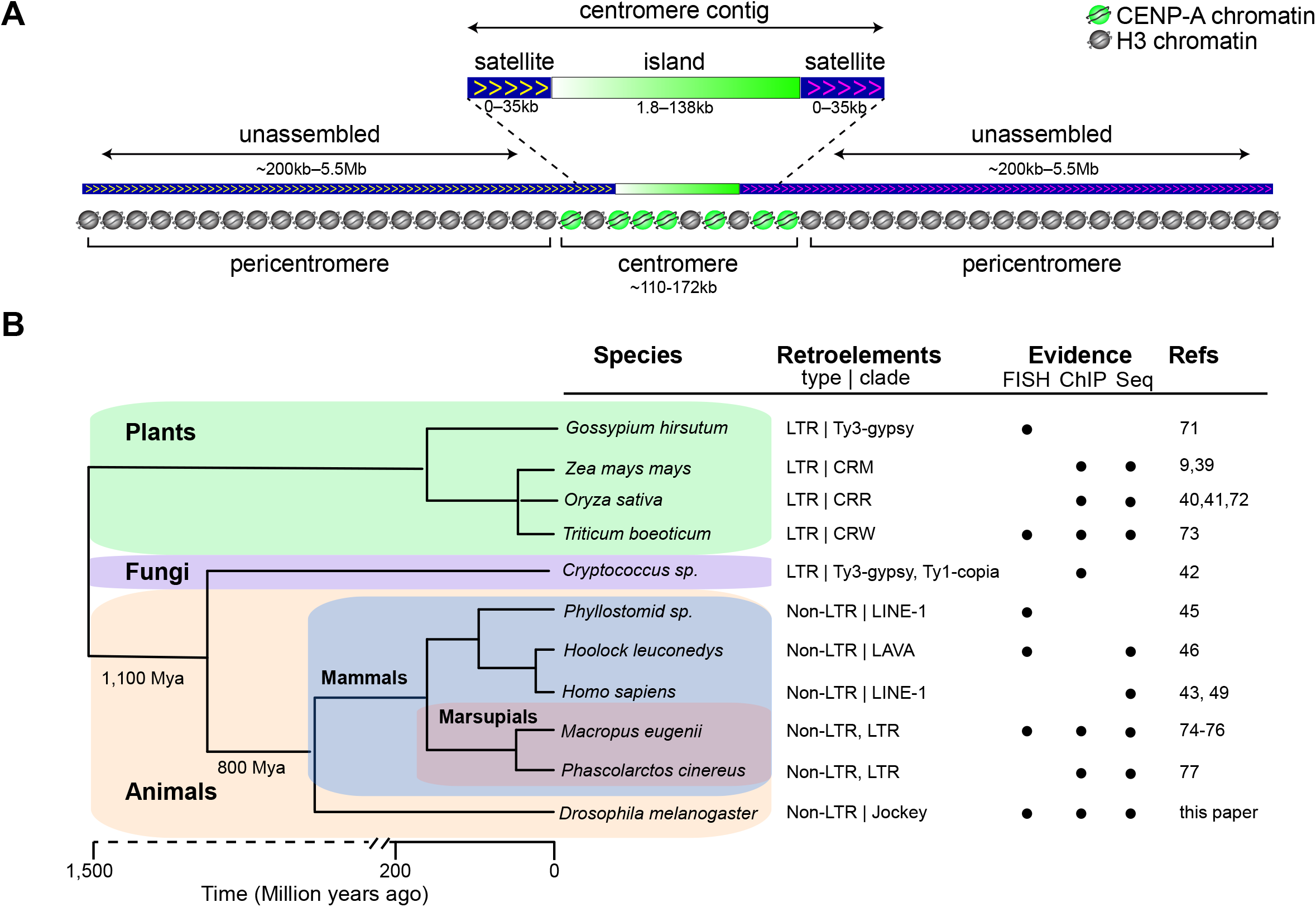
Drosophila centromere organization and widespread presence of retroelements at centromeres. **A)** Schematic showing centromere organization of *Drosophila melanogaster*. For at least centromeres X, 4 and 3 (since the sequences flanking the Y centromere are not in our assembly), the bulk of CENP-A chromatin is associated with the centromere islands, while the remaining CENP-A is on the flanking satellites. Although the complexity of island DNA allowed us to identify centromere contigs by long-read sequencing, the flanking satellites remain largely missing from our genome assembly, due to their highly repetitive nature. The approximate satellite size estimates are based on Jagannathan et al. [37]. **B)** Phylogenetic tree showing that centromere-associated retroelements are common across highly diverged lineages: *Gossypium hirsutum* (cotton) [71], *Zea mays mays* (maize) [9, 39], *Oryza sativa* (rice) [40, 41, 72], *Triticum boeoticum* (wild wheat) [73], *Cryptococcus* [42], *Phyllostomid* (bat) [45], *Hoolock leuconedys* (gibbon) [46], *Homo sapiens* (human) [43] and a human neocentromere [49], *Macropus eugenii* (tammar wallaby) [74–76], *Phascolarctos cinereus* (koala) [77], *Drosophila melanogaster* (this paper). The Phylogeny was constructed using TimeTree [78]. Indicated are the retroelement type and the clade that the element belongs to with element types as follows: LTR, Long Terminal Repeat retroelement; Non-LTR, Non-Long Terminal Repeat retroelement. The circles indicate the experimental evidence for centromere-association of retroelements: FISH, CENP-A ChIP-seq (ChIP), and genome or BAC sequencing (Seq).

To date, satellite DNAs have been regarded as the main sequence components of the centromeres of primary animal model systems—humans, mice, and *Drosophila* [2, 3, 17]. However, retroelements are also abundant and widespread at the centromeres of plants such as maize [39] and rice [40, 41]. Retroelements are also found at the centromeres of fungi [42], humans [43], marsupials [44], bats [45], and gibbons [46], suggesting that they are pervasive centromeric features (Fig. 7B). Our study shows that retroelements, particularly *G2/Jockey-3*, are not merely present near centromeres but are components of the active centromere cores through their association with CENP-A. Our BLAST search for *G2/Jockey-3* retroelements suggests that they are restricted to the *melanogaster* subgroup, therefore we hypothesize that different non-LTR retroelements may be present at the centromeres of other *Drosophila* species. Why retroelements are such ubiquitous components of centromeres and if they play an active role in centromere function remain open questions. In maize, centromeric retroelements invade neocentromeres following their inception [47], suggesting a preference for DNA sequences associated with CENP-A chromatin for retroelement insertion [18]. On the other hand, a LINE-element was found to be an integral component of a human neocentromere [48, 49], raising the possibility that it is CENP-A that may bind preferentially to retroelement-associated genomic regions [18]. Other models have proposed that retroelements could produce non-coding RNAs that affect centromere specification [18, 48], and that retroelement activity could help maintain centromere size through retrotransposition or by giving rise to tandem repeats via recombination-mediated mechanisms (*e.g*. [50, 51]; reviewed in [52]). Centromeric transcription contributes to centromere homeostasis in several organisms, including fission yeast [53, 54], wallaby [55], human [6, 56], and *Drosophila* cells [57, 58]. Our reanalysis of publicly available total RNA-seq reads was inconclusive about steady state transcription levels coming from centromeric contigs (data not shown) due to extremely low expression levels and insufficient read length, however our preliminary analysis with quantitative RT-PCR with centromere-specific *G2/Jockey-3* primer sets shows some evidence for a low level of centromere expression.

In addition to retroelements, the centromeres of *D. melanogaster* display a diverse assortment of repeats, none of which are exclusive to centromeres, with the exception of IGS, for which we identified a centromere-enriched variant. The identification of the IGS tandem repeat within 3^*Giglio*^ is intriguing, as IGS sequences are dynamic in the potato [59], where they are located near the centromere, as well as in the tobacco [60], the tomato [61], and the common bean [62], where they show a dispersed pattern over several chromosomes. The origin of novel tandem repeats is still elusive, but one way it has been proposed to occur for the IGS repeat in plants is through the initial insertion of a retroelement within rDNA, followed by IGS duplication, amplification, and transposition to a new locus [61].

Defining the span of the CENP-A domain is important to understand precisely which sequences are associated with centromere activity and which are part of pericentric heterochromatin. Although we are able to confidently map our ChIP-seq reads to the islands to determine CENP-A occupancy, the same cannot be done for simple satellites due to the limitations of mapping to highly repetitive DNA. We therefore infer the organization of the centromere from analyzing extended chromatin fibers by IF/FISH. Blocks of simple satellite sequences flank the islands on each of our contigs, with the exception of the Y centromere contig, however, these regions represent only a fraction of the estimated abundance of those repeats in the genome. For example, *dodeca* satellite occupies ~1Mb of the genome [63] and we have ~570 kb total *dodeca* sequence in the assembly, however only ~35 kb of *dodeca* is on the centromere 3 contig. Therefore, for many satellite sequences, inferences based on read mapping, even uniquely mapped reads, are confounded because satellites are underrepresented in the assembly. Our analysis of chromatin fibers suggests that CENP-A spans beyond the islands into the simple satellites, although the precise boundaries remain elusive (Fig. 7A).

The finding that CENP-A can bind to several different sequences that are not uniquely associated with centromere regions is consistent with the epigenetic model of centromere specification, which proposes that specific sequences alone do not govern centromere activity [3]. Yet it is possible that the diverse sequence arrangements observed at each centromere somehow contribute to centromere activity or specification [18, 39]. Possible mechanisms include the promotion of unusual types of transcription, as reported for fission yeast [64], or the formation of non-B DNA structures (*e.g*. stem loops, hairpins, and triplexes) that may promote CENP-A deposition [7, 12, 65]. Knowing the identity of *D. melanogaster* centromeric DNA will enable the functional interrogation of these elements in this powerhouse model organism.

### Author contributions

AML and BGM conceived the project, designed experiments, and wrote the manuscript. AC, JP, BS, and BGM performed wet lab experiments and analyzed data; C-HC, XW, and AML performed computational experiments and analyzed data; NMCM and BJB designed Oligopaints; JE and C-TW provided resources; all authors contributed to editing the manuscript and writing the supplemental results and methods.

## Supporting information

Supplemental results

## Acknowledgments

We are grateful to Gary Karpen for the anti-CENP-A antibody and for comments on the manuscript. We also thank Guy Nir (Oligopaint design), Kevin Wei (kseek), Karen Miga (discussion), Tom Eickbush (discussion), Rachel O’Neill (discussion and comments on the manuscript), Sarah Trusiak (early stages data analysis), Bo Reese and the University of Connecticut Center for Genome Innovation (sequencing resources), and the University of Rochester Center for Integrated Research Computing (computing cluster resources). We thank members of Larracuente’s and Mellone’s labs for comments on the manuscript. Portions of computational work for Oligopaints design were conducted on the Orchestra and O2 High Performance Compute Clusters, supported by the Research Computing Group at Harvard Medical School.

## Materials and methods

### Chromatin immunoprecipitations sequencing (ChIP-seq)

CENP-A ChIPs were performed using an affinity purified rabbit anti-CENP-A antibody (gift of Gary Karpen) that we previously verified works well for ChIP using S2 cells that contain LacI/lacO inducible ectopic centromeres and showing that CENP-A ChIP pulled-down lacO sequences [79] (Fig. S1A).

### ChIP in S2 cells

Chromatin from 10^6^ fixed *Drosophila* Schneider (S2) cells (approx. 90μg) were used for each IP and chromatin was sheared to 100–300bp using a Covaris sonicator. ChIPs were performed using the MAGnify kit (Thermo Fisher). 1μl of the anti-CENP-A antibody was coupled to 10μl of beads for 2 hours followed by incubation with chromatin overnight at 4°C. DNA was eluted in 50μl of elution buffer. Libraries were generated using the TruSeq kit (Illumina) and paired-end sequenced using the Reagent kit v.3. (Illumina) on the NextSeq platform.

### ChIP in Embryos

Embryo (OregonR) collection, fixation, and chromatin isolation were performed as described in [80]. We carried out four ChIP replicates as follows. From one embryo collection, we generated chromatin used in ChIP1, from a second independent embryo collection, we generated chromatin used for replicates, ChIP2-4. We used formaldehyde-crosslinked overnight collections of Oregon R embryos (~1.5g per collection). Chromatin was sheared to 200–500bp using a Bioruptor sonicator (Diagenode), aliquoted, and flash frozen. The first biological replicate (ChIP1) was performed following the protocol in [80] using 165μg of chromatin in 500μl volume and 30μl of protein A agarose beads, and 2μl of anti CENP-A antibody. For ChIP2, 3, and 4 we used the MAGnify kit, with 15μl of dynabeads, approximately 60μg of chromatin in 200μl volume, and 3μl of anti-CENP-A antibody. Libraries were made from eluted DNA using the TruSeq ChIP kit (Illumina) for ChIP1 and 4, while the Accel-NGS 2S Plus DNA Library (Swift Biosciences) was used for ChIP2, 3. Note that ChIP2-3 where performed in parallel and were sequenced the same way and are thus technical replicates. The libraries were sequenced by paired-end on the NextSeq platform using Reagent v.2. Chromatin extracted from the second embryo collection was also used for ChIP-qPCR experiments.

For both chromatin preparations, the quality of the chromatin was confirmed by control ChIPs with 15μg of chromatin in 200μl volume and 2μl of rabbit anti-H3K27Ac (Thermo Fisher). The eluted DNA was analyzed by qPCR confirming enrichment of the *Rpl32* promoter (F-TTGTTGTGTCCTTCCAGCTTCA and R-TTGTTGTGTCCTTCCAGCTTCA), and lack of enrichment of *Rpl32* 5’ region (F-GGCACGGCGCCAAAATTAATCA and R-CCGATGCCACTGCCTCTTTGGT [81, 82].

### ChIP quality control analyses

We estimated read quality of each replicate ChIP experiment using two metrics estimated in phantompeakqualtools [83]: the Normalized Strand Coefficient (NSC) and the relative strand correlation (RSC) (Table S5). These statistics report the cross-correlation between Watson and Crick strands, as ChIP reads from a true positive are expected to be highly clustered and accumulate on either side of the binding site on both strands, with a shift between the peaks on the Watson and Crick strands that is determined by read length and fragment length distribution [84]. This shift should not occur in the input. NSC is the fragment-length cross-correlation peak divided by the background cross-correlation and RSC is the fragment-length cross-correlation peak divided by the read-length peak [83].

### Analysis of repeat enrichment in ChIPs

To determine CeNp-A enrichment in simple tandem repeats, we summarized repeat composition in the trimmed reads and identified overrepresented kmers using kseek (https://github.com/weikevinhc/k-seek; [24]). The CENP-A/Input ratio is normalized by the number of mapped reads to the genome assembly to remove possible read contamination. We consider a class of repeats to be enriched for CENP-A if the minimum number of kmers in the input is ≥10 in each replicate and the median normalized CENP-A/Input ratio is > 1 across all four replicate ChIP experiments (Fig. S1). Simple tandem repeats maybe over or underrepresented due to Illumina library preparation and the effects of PCR amplification on sequence library complexity. To determine CENP-A enrichment on complex repeats, we used a mapping approach. We annotated repeats in our assembly [19] using a custom Drosophila-specific consensus repeat library [34] modified from RepBase to include complex satellite DNAs (Repbase version 20150807; [85]; Supplemental File 1). Using these RepeatMasker annotations, we generated a comprehensive library of all individual repetitive elements in the genome to capture sequence variation among repeats. We mapped ChIP and input reads to this comprehensive repeat library using bowtie2 (default parameters) and summarized read counts for each type of complex repeat (*e.g*. transposable elements, complex satellite DNAs with repeat units >100 bp) using custom python scripts. The CENP-A/Input ratio is normalized by the number of mapped reads to the genome assembly. We consider a class of repeats to be enriched for CENP-A if it is in top 20^th^ percentile of normalized CENP-A/Input in all four replicate ChIP experiments.

To address if any motif in *G2/Jockey-3* is particularly enriched for CENP-A, we constructed a consensus sequence of *G2/Jockey-3* in *D. melanogaster* and *D. simulans*. We mapped ChIP and input reads to this comprehensive repeat library with only one version of *G2/Jockey-3* (either *D. melanogaster* or *D. simulans*) using bwa (default parameters). We then called the depth of reads with samtools depth (v1.7) using “-Q 10 (mapping quality ≥ 10)” and calculated ChIP/Input ratio across each site after normalization by the number of mapped reads to the genome assembly.

### *De novo* ChIP assembly

We used k-mer-based *de novo* assembly methods to detect CENP-A-enriched regions [20]. We trimmed reads using TrimGalore v0.4.4 (https://www.bioinformatics.babraham.ac.uk/projects/trim_galore/) and the parameters “-gzip-length 35 -paired”. For the second replicate, we further subsampled reads to 100x coverage using Bbnorm (v37.54, https://sourceforge.net/projects/bbmap/) with the parameters “threads=24 prefilter=t target=100” for the *de novo* assembly. We created *de novo* ChIPseq contigs (ChIPtigs) with Spades v3.11.0 (-t 24 -careful -sc;[86]) for each replicate (Supplemental Files 2-6). To calculate CENP-A enrichment, we mapped input and ChIP reads to the ChIPtigs. We masked duplicates using Picard MarkDuplicates (v2.12.0; http://broadinstitute.github.io/picard) and filtered low-quality reads with samtools (v1.7) using “-f 3 -F 4 -F 8 -F 256 -F 2048 -q 30” for the paired-end reads and “-q 30” for the single-end reads to keep high-quality reads (mapping quality ≥30 and properly paired). We calculated the enrichment P-value using MACS2 (version 2.1.1.20160309; -q 0.01 --call-summits; [87]). ChIPtigs were mapped back to our assembly using megablast BLAST 2.6.0+ [88] with default setting, and the best hits were chosen. We removed potentially misassembled ChIPtigs and adjusted the peak regions in the reference sequence using custom scripts. We identified 1919, 16310, 14667, 4916 significantly CENP-A enriched ChIPtig regions from 127426, 268663, 625927 and 184133 total *de novo* ChIPtigs for each replicate, respectively (Table S3).

### Identification of candidate centromeric contigs

We identified candidate centromeric contigs in the new iso-1 assembly[19] based on organization: we looked for contigs containing complex DNA flanked by satellites with known centromeric and pericentric locations. We first generated the assembly with long-read sequence data, including PacBio [89] and nanopore reads (Table S4; [90]). We filtered nanopore reads using Porechop and Filtlong (--min_length 500) to remove adaptors and short reads (https://github.com/rrwick/Porechop and https://github.com/rrwick/Filtlong). We assembled the nanopore and PacBio reads into a hybrid assembly using Canu v1.7 with default settings [91]. Our new hybrid PacBio-Nanopore assembly is less contiguous than our previous PacBio-only assembly despite using more reads (Supplemental File 7; Assembly size = 162,798,260 bp in 798 contigs; N50=5,104,646 bp). We thus decided to use our PacBio-only assembly [19], which has a greater representation of heterochromatin compared to previously published assemblies (see details in [19]). To ensure that we were not missing putative centromeric contigs, we looked for sequences with CENP-A-enriched repeats (based on our repeat analysis; Fig. 1B; Fig. S1) in the error-corrected PacBio and nanopore reads, and the hybrid assembly, that were missing from our PacBio-only assembly. We were particularly interested in contigs containing repeat sequences that we identified as enriched in our ChIP data. To annotate contigs and unassembled corrected-reads, we used RepeatMasker 4.06 [92] with Repbase 20150807 and parameters “-species drosophila -s” to annotate interspersed repeats (described above) and Tandem Repeat Finder (TRF v4.09; [93]) to annotate tandem repeats. We extracted 19 non-redundant sequences from our new hybrid assembly and error-corrected reads with candidate centromeric repeats, including *dodeca, Prodsat*, AATAG, IGS^3cen^ (as determined by phylogenetic analysis below), and *G2/Jockey-3* sequences. We added these 19 candidates to our new PacBio-only assembly [19] (Supplemental File 8) to create the final version of our assembly. We polished the final assembly 10 times using Pilon (v1.22 [94]; with Illumina [95, 96] and long synthetic reads[97] (Table S6; with parameters “--mindepth 3 --minmq 10 --fix bases”). We annotated the finished assembly using our customized repeat library (-lib library.fasta -s) and RepeatMasker 4.06 [92] (Supplemental File 9). Additionally, we transferred gene annotations from Flybase r6.20 to our genome using BLAT [98] and CrossMap v0.2.5 [99] (Supplemental File 10).

### Peak calling

We mapped our input and ChIP reads and publicly available data [16] to our genome assembly with new candidate sequences [19] using bwa v0.7.15 [100]. We masked duplicates using Picard MarkDuplicates (v2.12.0; http://broadinstitute.github.io/picard) and filtered low-quality reads with samtools (v1.7) using “-f 3 -F 4 -F 8 -F 256 -F 2048 -q 30” for the paired-end reads and “-q 30” for the single-end reads to keep high-quality reads (mapping quality ≥30 and properly paired). We then called peaks using MACS2 (version 2.1.1.20160309; -q 0.01 —call-summits; [87]) with the alignments. We report top 100 peaks with strongest signal from each replicate (7^th^ column of narrowpeaks files). We used Irreproducible rate (IDR [101]) to overlap the datasets and identify high confidence peaks (https://github.com/nboley/idr) between every replicate (IDR<0.05 corresponding to an IDR score ≥ 540). Since there are many peaks with weak CENP-A enrichment in the comparison between R2 and R3 (16,870), we only chose 37 peaks—the average peak number of other comparisons (27-44)—with strongest signals for our figures (Table S5).

### ChIP qPCR

Quantitative real-time PCR (qPCR) was performed using SYBR-green (Bio-Rad) on a CFX96 Real-Time System (Bio-Rad). 1 μl of Input or ChIP eluted DNA was used in each qPCR reaction. Melting curves were analyzed to ensure primer specificity. Only primers with reaction efficiencies within a linear dynamic range were used. The fold-enrichment of centromeric DNA after immunoprecipitation of CENP-A chromatin compared to its level in the bulk input chromatin was calculated with the equation: 100*E^(Ct*input*-C*tip*)^, where E is the efficiency of the primer set. Enrichment values were normalized by the enrichment value of *RpL32* as a noncentromeric control. qPCR primer sets are listed in Table S7.

### Transcription of centromeric *G2/Jockey-3* elements

Total RNA was extracted from three independent overnight collection of embryos (iso-1). Briefly, embryos were scooped from the apple juice plates and rinsed with water in a mesh basket, dechorionated in 50% bleach for 3.5 min with gentle shaking, rinsed thoroughly with water, moved to a 1.5ml microfuge tube, and resuspended in 300μl of Trizol reagent (Sigma Aldrich). Embryos were homogenized using a motorized pestle until the solution became clear (30-40 seconds). The homogenized solution was centrifuged at 13000rpm for 10 min at 4°C and the clear supernatant was transferred to a new RNAse free tube. RNA was isolated using the Direct-Zol RNA miniprep plus kit (Zymo Research) according to the manufacturers’ protocol.

The RNA was eluted in RNAse free water and quantified with a Nanodrop. A total of three consecutive Turbo DNAase (Invitrogen) followed by RNAeasy Clean up (Qiagen) treatments were performed to remove DNA contamination.

Reverse-transcription was performed using the iScript Select cDNA Synthesis kit (Bio-Rad) according to the manufacturer’s instructions. Briefly, 75ng of total embryo RNA were used to make cDNA libraries using random priming in a 30μl reaction. For the no-RT control, the Reverse Transcriptase was omitted from the reaction.

qPCR was performed as described for ChIP qPCR using 1μl of cDNA in each reaction and primers sets targeting *G2/Jockey-3* copies from each centromere (X-G2, 4-G2, Y-G2, 3-G2, 2-G2). Primers for *Actin5C* were used as a positive control for a highly expressed gene, while a primers for the testis-specific gene, *Mst84Da*, was used as a control for a non-expressed gene. The no-RT samples produced Ct values comparable to the negative non-expressed control showing successful removal DNA.

Gene expression was analyzed as in Schmittgen et al. [102] by determining the mean 2^−ΔCt^, where ΔCt is (Ct_G2/jockey-3_-Ct_Mts84Da_), from three biological replicates. Primer sets are listed in Table S7.

**Table S7.**
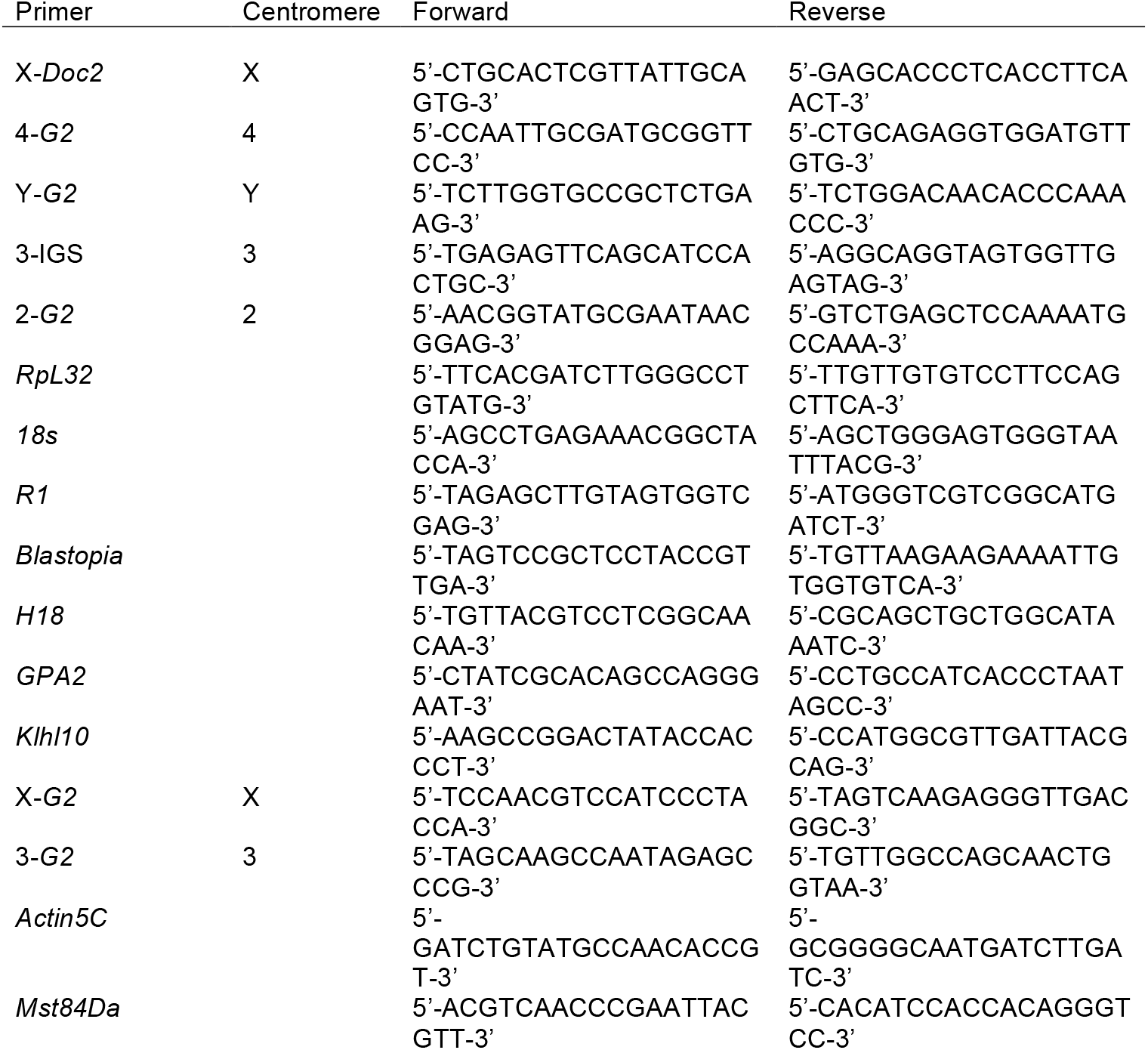
List of qPCR Primers. List of primers used for qPCR in this study. The centromere contig that each target is associated with (X, 4, Y, 3, 2) is designated in the “Centromere” column. Note that *in silico* PCR for the 3_G2 primers predicted 3 specific products from centromere 3, as well as two products on contig tig00022795 and additional non-specific products from the X chromosome when 3 or more mismatches are allowed all of the same 145bp size.

### Immunofluorescence (IF) and Fluorescence *in situ* hybridization (FISH)

#### IF-FISH on mitotic chromosomes

##### S2 mitotic chromosome preparation

Preparation of mitotic chromosomes from *Drosophila* S2 cells was performed as described in [79]. 2×10^5^ cells were treated with 0.5μg/mL demecolcine solution (Sigma-Aldrich) and incubated at 25°C for 1 hour to induce a mitotic arrest. Cells were pelleted (600g/5 min) and resuspended in 250μL 0.5% (w/v) sodium citrate for 8 min. Cells were loaded into cytofunnels and spun onto Superfrost^®^ Plus slides (VWR) at 1200rpm for 5 min using a Shandon Cytospin 4 (Thermo Fisher Scientific). Cells were fixed for 10 min with 3.7% formaldehyde in PBS, 0.1% Triton-X-100 (PBS-T). Slides were washed three times in PBS-T for 5 min and stored at 4°C until ready for use.

##### D. melanogaster and D. simulans mitotic chromosomes preparation

Preparation of mitotic spreads was carried out from iso-1 *D. melanogaster* flies (Bloomington Drosophila Stock Center stock n. 2057: *y^1^; Gr22b^iso-1^ Gr22d^iso-1^ cn^1^ CG33964^iso-1^ bw^1^ sp^1^; MstProX^iso-1^ GstD5^iso-1^ Rh6^1^*) and *D. simulans* (w501, gift of Andy Clark) in larvae following the method in [103] with minor modifications. Male 3^rd^ instar larval brains were dissected in PBS and immersed in 0.5% (w/v) sodium citrate for 8 min. Individual brains were fixed for 6 min in 6μL of 45% acetic acid, 2% formalin on siliconized coverslips. Whole brains were applied to clean poly-L-lysine slides (Thermo Fisher Scientific) and were manually squashed between coverslip and slide by pressing with the thumb. Slides were immersed in liquid nitrogen. Once bubbling stopped, the slides were removed from liquid nitrogen and the coverslip was immediately removed using a razor blade. Slides were immediately immersed in PBS and were either washed for 5 min before proceeding to IF or stored at 4°C in PBS until ready for use.

##### Immunofluorescence staining

For IF, slides were washed in PBS-T for 5 min. S2 cell slides were blocked in 5% milk in PBS-T for 30 min. Larval squashes were blocked in 1% BSA, PBS, 0.02% sodium azide for 30 min. Primary antibodies anti-CENP-A (larval brain slides: rabbit, 1:500, Active motif; S2 cell slides: chicken, 1:1000, [70] and anti-CENP-C (larval brain slides: guinea pig, 1:500 [27]) were diluted in blocking solution and incubated on slides overnight at 4°C. Slides were washed three times for 5 min in PBS-T and incubated with secondary antibodies (Life Technologies Alexa-488, 546, or 647 conjugated, 1:500) diluted in blocking solution and incubated at room temperature for 1 hour or overnight at 4°C. Slides were washed three times for 5 min in PBS-T.

##### Satellite FISH

Satellite FISH was performed following the protocol described in [104] with a few modifications. Slides were post-fixed in 3.7% formaldehyde, PBS for 10 min, followed by a rinse in PBS and two 5 min washes in 2×SSC, 0.1% Tween-20 (2×SSC-T). Slides were washed once for 5 min in 50% formamide, 2×SSC-T at room temperature, once for 20 min in 50% formamide, 2×SSC-T at 60°C, and then cooled to room temperature. For FISH, 25μL of hybridization mix containing 40pmol of each probe, 2×SSC-T, 10% dextran sulfate, 50% formamide, and 1μL of Rnase Cocktail (ThermoFisher) was applied to a 22 × 22mm hybrislip (Electron Microscopy Sciences), mounted on the slide and sealed with paper cement. Slides were denatured at 92°C for 2.5 min and then incubated overnight at 37°C. Slides were washed in 2×SSC-T at 60°C for 20 min, followed by two 5 min washes in 2×SSC-T at room temperature, and one 5 min wash in PBS. Slides were mounted in Slowfade^®^ Gold Reagent (Invitrogen) containing 1μg/mL DAPI and sealed with nail polish.

##### Oligopaint FISH

Oligopaint FISH was performed as described above with the following modifications. 25μL of hybridization mix containing 10pmol of Oligopaint, 2×SSC-T, 10% dextran sulfate, 60-68% formamide, and 1μL RNase cocktail was applied to a 22 × 22mm hybrislip (Electron Microscopy Sciences), mounted on the slide and sealed with paper cement. Slides were denatured at 92°C for 2.5 min in a thermocycler (Eppendorf) and incubated overnight at either 37°C or 42°C (see Table S10 for % formamide and hybridization temperatures used). For fluorescence detection, 10pmol of Alexa-488 labeled secondary oligo were applied either during the overnight hybridization or following post-hybridization washes, in which 25μL of 2×SSC, 30% formamide, 10pmol of probe was applied to each slide and incubated at room temperature for 30 min.

Slides were washed twice in 2×SSC, 40% formamide for 20 min, once in 2×SSC-T for 15 min, and once in PBS for 5 min. Slides were mounted as described above and successful hybridization was checked under fluorescent microscope. Satellite probes were added after imaging by removing the coverslip with a razor blade, washing slides three times in 2×SSC-T for 5min, applying 25μL of 2×SSC, 30% formamide, 40pmol of satellite probe to each slide and incubating at 37°C for 1 hour. Slides were washed once in 2×SSC-T at 60°C for 20 min, twice in 2×SSC-T for 15 min, once in PBS for 5 min, and mounted as described above.

##### G2/Jockey-3 FISH

FISH for *G2/Jockey-3* was performed as described in Dimitri *et al*. [105]. Slides were dehydrated in an ethanol row (successive 3 min washes in 70%, 90%, and 100% ethanol) and allowed to air dry completely. 20μL of probe mix containing 2×SSC, 50% formamide, 10% dextran sulfate, 1μL RNase cocktail, and 100ng of DIG-labeled G2 probe was boiled at 80°C for 8 min, incubated on ice for 5 min, and then applied to slides, covered with a glass coverslip and sealed with paper cement. Sealed slides were denatured on a slide thermocycler for 5 min at 95°C and incubated at 37°C overnight. Slides were then washed three times for 5 min in 2×SSC, 50% formamide at 42°C, three times for 5 min in 0.1xSSC at 60°C, and then blocked in block buffer 1% BSA, 4xSSC, 0.1% Tween-20 at 37°C for 45 min. Slides were incubated with 50μL of block buffer containing a fluorescein labeled anti-DIG antibody (sheep, 1:100, Roche) for 60 min at 37°C. Slides were then washed three times for 5 min in 4xSSC, 0.1% Tween-20 at 42°C, and mounted as described above.

**Table S10.** Oligopaint hybridization conditions. Hybridization conditions used for FISH with specific Oligopaints.

| Oligopaint | Formamide (%) | Temperature (°C) |
| --- | --- | --- |
| Maupiti | 60 | 37 |
| Lipari | 68 | 42 |
| Giglio | 68 | 42 |
| Lampedusa | 60 | 37 |
| 80C4 | 50 | 37 |
| 61C7 | 50 | 37 |

**Table S11.**
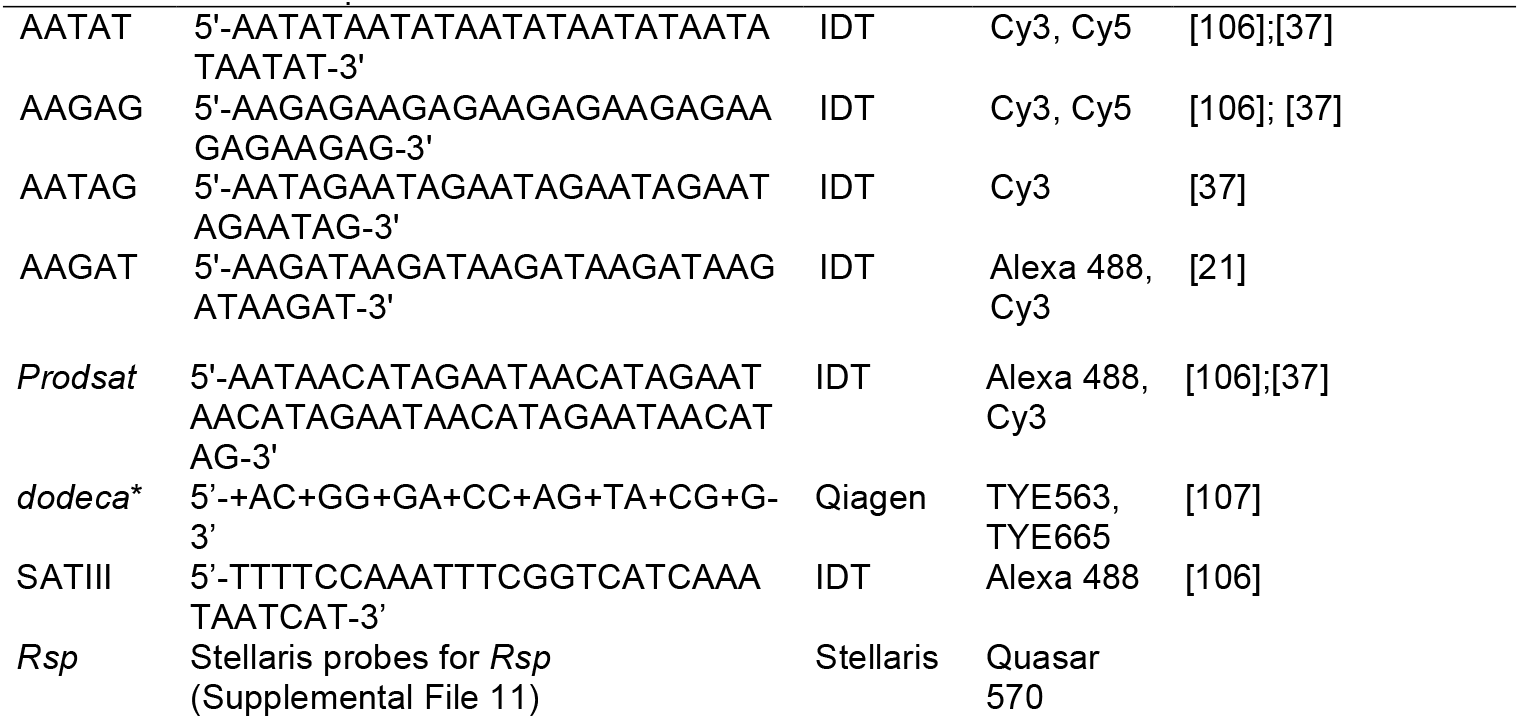
Labeled Satellite Probes. Information on the fluors used and sequences of satellite FISH probes used in this report. * = “+N” designates the incorporation of a locked nucleic acid (LNA).

**Table S12.**
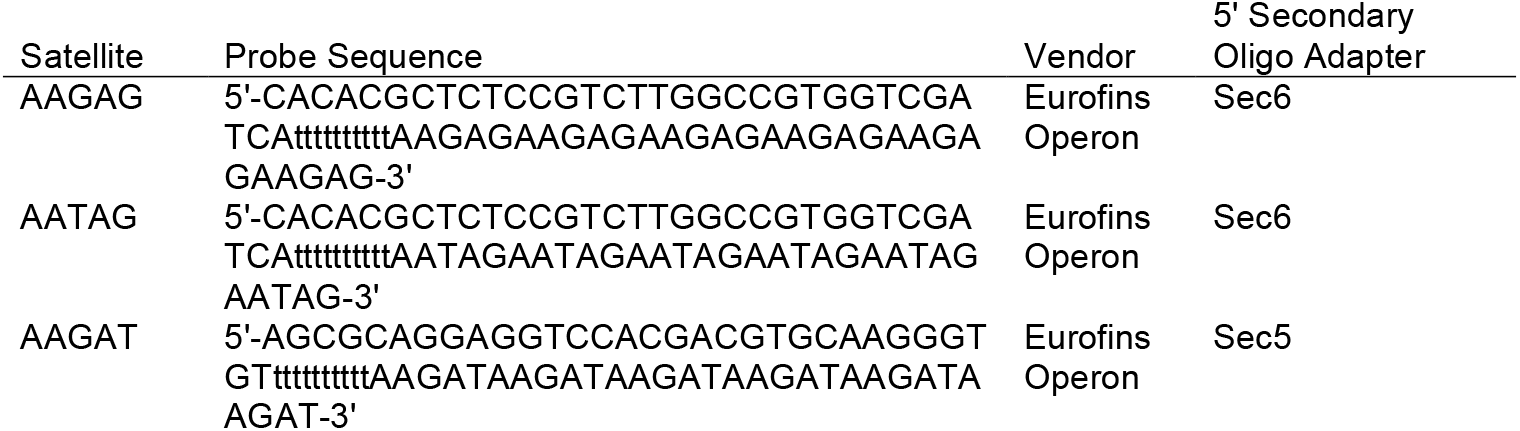
Unlabeled satellite probes. Information on the 5’ secondary oligo adapter site and sequence of satellite probes used in this report.

**Table S13.**
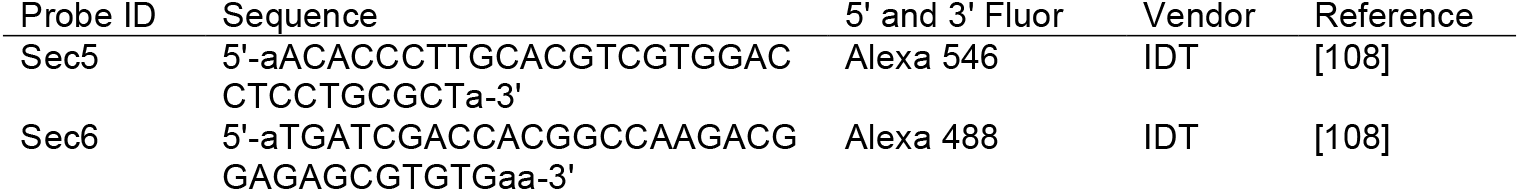
Secondary Oligo Probes. Sequence and fluors of secondary oligo probes used for fluorescence detection of Oligopaints and unlabeled satellite probes.

### Preparation of extended chromatin fibers and IF-FISH

Extended chromatin fibers were prepared as described in Sullivan [33], with a few modifications. 3–4 brain imaginal discs from third-instar iso-1 wandering larvae (females were selected to avoid cross-centromere hybridization of our X^*Maupiti*^ and 4^*Lampedusa*^ Oligopaints with the Y centromere while males were used for centromere Y) were dissected in 0.7% NaCl and dissociated in 250μl 0.5% (w/v) sodium citrate containing 40μg collagenase/dispase (Sigma-Aldrich) by incubating at 37°C for 10 min. This mixture was briefly vortexed and spun, and loaded into a single chamber Shandon cytofunnel for centrifugation in a Shandon Cytospin 4 at 1200rpm for 5 min onto a clean polysine slide (Thermo Fisher Scientific). After centrifugation, the slides were immediately immersed in a glass coplin jar containing lysis buffer (500mM NaCl, 250mM Urea, 25mM Tris-HCl pH7.4, 1% Triton X-100) for 13–15 min following which the slides were gently removed at a steady state of ~25–30 seconds per slide. Fibers were fixed in 4% formaldehyde solution and washed in PBS for 5 min. After washing, the slides were processed for IF-FISH.

Fibers were extracted in PBS-T for 10 min then incubated in a 1.5% BSA, PBS blocking solution for 30 min. Slides were incubated with an anti-CENP-A antibody (rabbit, 1:100, Active Motif) diluted in blocking solution overnight at 4°C in a humidified chamber. Slides were washed three times in PBS for 5 min and then incubated for 45 min with secondary antibodies (Cy5-conjugated donkey anti-rabbit, 1:500, Life Technologies) diluted in blocking buffer at room temperature, followed by three 5 min washes in PBS. Slides were post-fixed in 3.7% formaldehyde, PBS for 10 min followed by one quick rinse and two 5 min washes in PBS. FISH was performed as described for 3D-FISH [104] with a few modifications. Slides were washed twice in 2×SSC-T at room temperature for 5 min, followed by denaturation in 50% formamide, 2×SSC-T at room temperature for 5 min, transferred at 60°C for 20 min, then cooled to room temperature. 10pmol of primary Oligopaint probes (except X^*Maupiti*^ which was 25pmol) and 40pmol of satellite DNA probes were each added to the slides in 25μL of hybridization solution: 2×SSCT, 60% or 68% (v/v) formamide (see TableS10), 1μL RNase cocktail (Invitrogen), 10% dextran sulfate (Millipore) and sealed with a 22×22 hybrislip (Electron Microscopy Sciences) using rubber cement. Slides were then denatured at 92°C for 3 min on a slide thermo-cycler and allowed to hybridize overnight at 37°C or 42°C (See Table S10) in a humidified chamber. Slides were washed once in 2×SSC-T for 15 min at 60°C, once in 2×SSC-T for 10 min at room temperature, and once in 0.2×SSC for 10 min at room temperature. Following washes, 25μL of hybridization mix containing 2×SSC, 30% formamide, 40pmol fluor-labeled secondary oligo probes (see Table S13) was added on to each slide and incubated for 45 min at room temperature except for the X^*Maupiti*^ slides where the satellite probe was also added with the secondary Oligopaint probe. The slides were then washed once in 2×SSC-T at 60°C for 15 min, followed by one wash in 2×SSC-T and 0.2×SSC for 10 min at room temperature and mounted as described above.

For FISH with only satellite probes, post-hybridization washes consisted of one wash in 2×SSC-T for 20 min at 60°C, followed by one wash with 2×SSC-T at room temperature for 10 min and two 5 min washes in 0.2×SSC at room temperature. Slides were mounted in as described above.

For fiber measurement calibration, FISH using the 61C7 and 80C4 probes was performed using the conditions for Oligopaint FISH (see Table S10 for % formamide and hybridization temperatures used), while FISH using the *Rsp* probe was performed using the satellite FISH protocol.

### Microscopy and image analysis

Image acquisition was done at 25°C using an Inverted Deltavision RT restoration Imaging System (Applied Precision) equipped with a Cool Snap HQ^2^ camera (Photometrics) and 100x/1.40 NA oil immersion lens (Olympus). Image acquisition and processing was performed using softWoRx software (Applied Precision). For mitotic chromosomes, 20 z-stacks were taken per image at 0.2μm per slice. For fibers, 12–15 z-stacks were taken per image at 0.15μm per slice. Images were deconvolved using the conservative method for 5 cycles. Maximum intensity projections were made using 3–5 z-stacks. Images were saved as Photoshop files and were scaled using Adobe Photoshop. Figure assembly was done using Adobe Illustrator.

Maximum intensity projections of individual fibers were analyzed to measure the signal length of various signals on fibers using the “measure distances” tool in Softworks (GE). Three calibration probes of known length (100kb; see Tables S10 and 15 for 80C4 and 61C7 Oligopaints; see Supplemental File 11 for *Rsp* probe) were used to determine the degree of stretching in our experiments. At least 20 fibers for each probe were measured in all cases. Length measurements were visualized by scatter plot using Prism. These lengths were then used to determine the average stretching in kb/μm and student t-test was used for statistical analyses.

We noticed that the variation in the measurements for the island Oligopaints was greater than what we observed for the probes used for calibration. We attribute this higher variation to the lower density of island Oligopaint probes (some of the island sequences were not targeted by probes to increase specificity), which causes the signal to be weaker and less consistent than in standard Oligopaint FISH. It is also important to note that we analyzed fibers from a mixed population at different stages of the cell cycle, which could display differences in CENP-A density. It is also possible that the stretching of the chromatin at the centromere is more variable than at non-centromeric regions.

### Oligopaints

#### Design

Oligopaint libraries were designed using the OligoMiner pipeline [30, 109] with some variations. The genomic regions that showed significant enrichments of CENP-A via MACS and enriched ChIPtigs were targeted for Oligopaint design. The blockparse.py script (v1.3) using overlap mode was used to identify as many candidate probes as possible, with genome-targeting regions 35–41bp long, and a desired Tm of 42–47°C. Unlike standard Oligopaints design, the candidate probes were not aligned to the genome using Bowtie2 [110] or filtered with OutputClean.py, so that probes which align multiple times would not be discarded. Candidate probes with partial alignments of 18bp-long kmers were filtered out using kmerfilter.py (v1.3) and Jellyfish [111], excluding any that matched 6 or more times to the genome. Probes were filtered further for least secondary structures using StructureCheck.py (v1.3) and NUPACK [112]. Finally, coverage and density of probes across the regions of interest and presence of densely clustered off-target alignments were manually checked by Bowtie2 alignment, filtering for different levels of mismatch to assess the effects of hybridization stringency.

For the design of control regions for length standards in chromatin fiber stretching measurements, in loci 80C4 (3L: 23, 047,118..23,147,118) and 61C7 (3L: 626,646..726,646), we used conventional Oligopaint design for non-repetitive genomic regions. The blockparse.py script (v1.3) was used to identify candidate probes, with genome-targeting regions 35–41bp long, and a desired Tm of 42–47°C. Candidate probes were then aligned to the dm6 reference genome (with NNN masking of repetitive regions) using Bowtie2, and its output filtered using outputClean.py (v1.5.4), to keep only those probes that are predicted to thermodynamically only hybridize on-target under the specific conditions used. Finally, candidate probes were then further analyzed through kmerFilter.py (v1.3) to reject any probes containing regions of microhomology to off-target sites, and through StructureCheck.py (v1.3) to exclude any probes forming restrictive secondary structures.

Each oligo included universal primers at the 5’ and 3’ ends for PCR amplification, and a library-specific barcode for both PCR amplification and FISH detection of each individual centromere set. One library per centromere was synthesized as a single ChIP by Custom Array.

#### Library amplification

Raw libraries were amplified in 100μL reactions containing 10μL KAPA Buffer A and 1μL KAPA Taq from the KAPA Taq PCR Kit (Fisher Scientific), 1μL of library, 0.4mM dNTPs, 2μM of each universal primer, and amplified using the following cycles: 95°C for 5 min; 25 cycles of 95°C for 30 seconds, 58°C for 30 seconds, 72°C for 15 seconds, and a final extension at 72°C for 5 min. Reactions were purified using DNA Clean & Concentrator^™^-5 (Zymo Research) using the manufacturer’s protocol. Universal primers are listed in Table S14.

**Table S14.**
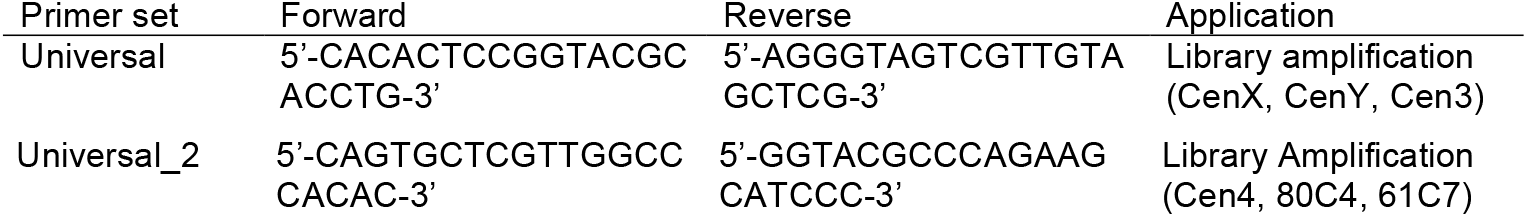
Universal primers. List of primer sets used for library amplification and G2 probe synthesis.

Sub-libraries were amplified in two 100μL reactions containing 10μL KAPA Buffer A and 1μL KAPA Taq from the KAPA Taq PCR Kit (Fisher Scientific), 0.5ng of amplified library, 0.4mM dNTPs, 0.4μM of each sub-library-specific primer and amplified using the following cycles: 95°C for 5 min; 35 cycles of 95°C for 30 seconds, 60°C for 30 seconds, 72°C for 15 seconds, and a final extension at 72°C for 5min. Reactions for individual sub-libraries were pooled and purified as describe above. Sub-library-specific primers are listed in Supplemental Table 15.

**Table S15.**
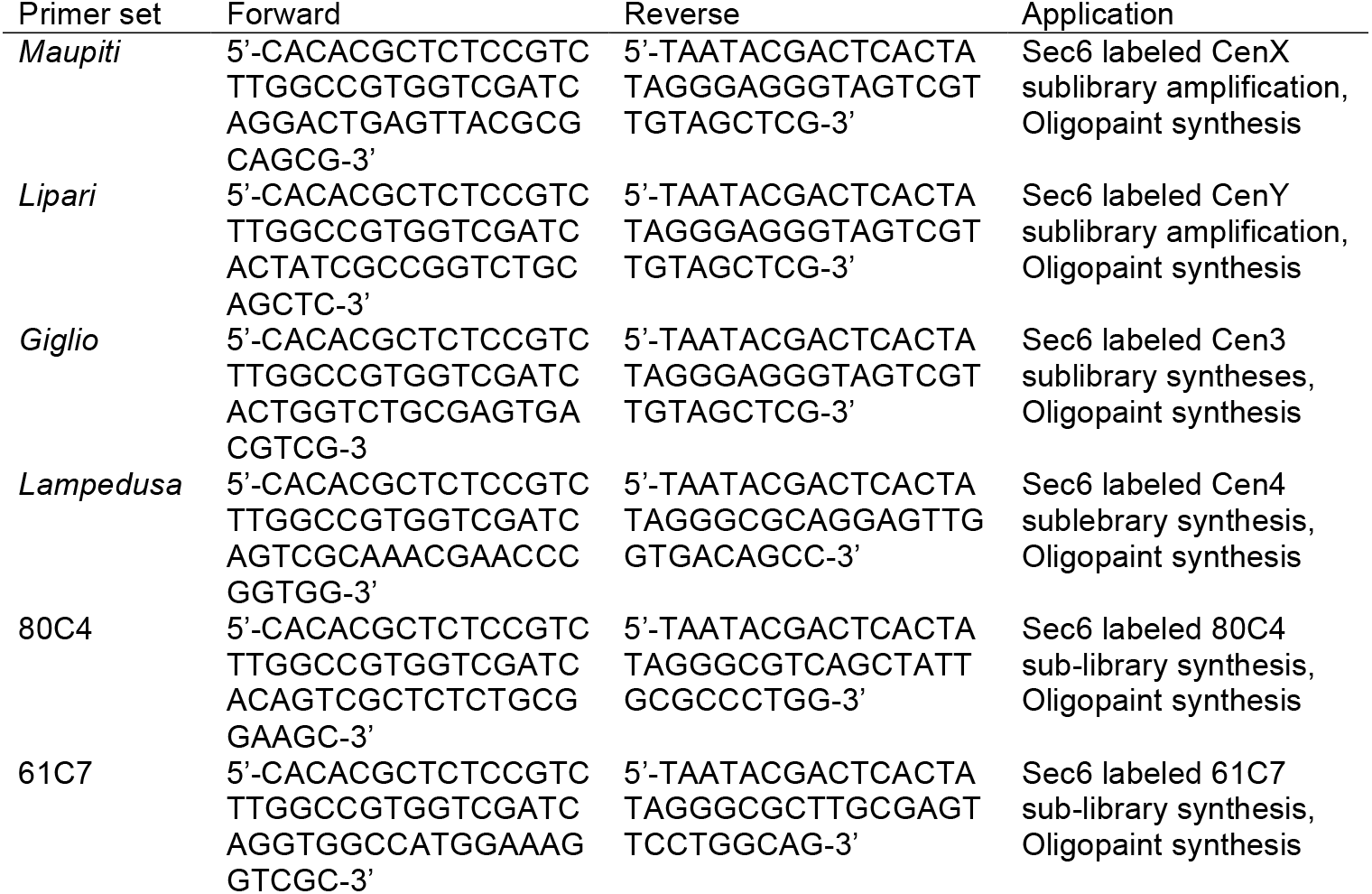
Sub-library-specific primers. List of primer sets used for sub-library amplification and Oligopaint synthesis.

#### Oligopaint Synthesis and Purification

T7 RNA synthesis was performed in 40μL reactions containing 4μL 10x T7 Buffer, 4μL each NTP, and 4μL T7 Pol Mix from the MEGAscript™ T7 Transcription Kit (Thermo Fisher Scientific), 2μg of amplified sub-library, and 2μL RNaseOUT™ Ribonuclease Inhibitor (Thermo Fisher Scientific). Reactions were incubated at 37°C for 20 hours. cDNA synthesis was performed in 300μL reactions containing the entire T7 RNA synthesis reaction, 10μM sublibrary-specific forward primer, 1.6mM dNTPs, 60μL 5x RT Buffer and 4μL Maxima H Minus Reverse Transcriptase from the Maxima H Minus Reverse Transcriptase kit (Thermo Fisher Scientific), and 3μL of RNaseOUT. Reactions were incubated at 50°C for 2 hours, followed by heat inactivation at 85°C for 5 min. RNA hydrolysis was performed by adding 300μL of 0.25M EDTA, 0.5M NaOH to cDNA synthesis reactions and incubating at 95°C for 5 min. Reactions were then put on ice. Oligopaints were purified using DNA Clean & Concentrator^™^-100 (Zymo Research) following the manufacturer’s protocol, substituting 4.8mL of 100% ethanol and 1.2mL of Oligo Binding Buffer (Zymo Research) instead of the DNA Binding Buffer. Oligopaints were eluted using 150μL mqH_2_O. The concentration of each Oligopaint was determined using a NanoDrop™ 2000c Spectrophotometer using the ssDNA setting. The molarity of each Oligopaint was calculated using the following formula: eluate concentration [ng/μL]*(1pmol/330pg)*(1/Oligopaint length in nt) = Oligopaint molarity [μM]

#### *G2/Jockey-3* Probe

##### Probe design

We designed a 1643 bp *G2/Jockey-3* oligo against the consensus of the 3’ region of *G2/Jockey-3* elements found within most centromere contigs. A 5’ addition containing 5’-CAGT-3’ followed by universal forward primer binding sites separated by an XhoI cut site (5’-cacactccggtacgcacctgctcgagcagtgctcgttggcccacac-3’). A 3’ addition containing 5-ACTG-3’ followed by universal reverse primer binding sites separated by a SpeI cut site (5’-agggtagtcgttgtagctcgactagtggtacgcccagaagcatccc-3’). The *G2/Jockey-3* sequence was ordered as a “custom gene” (IDTDNA.com) and synthesized in the pUCIDT (AMP) vector (pUCIDT-G2). The sequence of the insert is as follows (primer binding sites = *italicized;* restriction sites = bold; *G2/Jockey-3* sequence = CAPITALIZED).

5’-cagtcacactccggtacgcacctg**ctcgag**cagtgctcgttggcccacacCGGACGGCTCTTGGTGCCGCTCT GAAGCCGAAAGAGCTGAAGCGTTTGCAGATCACCTCCAGAATGCATTCACACCATTTGACAGATGCACTGGCGAAGAGCGTGCTGCAACCACCAGGTTCCTAGAGAGTCCATGTCCTCCTAGCCTGCCCATAGAGCCCGTCACCCCAGAAGAGGTTGCGCAAGAGTCGCCTCACTAAAGGCTAGCAAATCCCCAGGACTGGATCGCATCGACGCCACATCCCTTAAAATGCTGCCACCTCCCTGTTCCCAGTTGCTGGCCAACATATACAACAGATGCTTCTCACTAGGGTACTTCCCGAGATCATGGAAACGTGCAGAAGTCATTCTCATCCTCAAACCTGGAAAACCTGAAGCCAATCTTGCCTCATATAGACCGATTAGTCTGCTGGCAATCCTCTCCAAAATACTCGAAAGAGTATTTCTGCGCAGAGTGTTGCCAGTACTGGACGAGGCTGGACTGATCCCTGATCACCAGTTTGGCTTCAGGCGATCCCACGGAACACCCGAGCAATGCCACCGGCTCGTAGCACGCATCCTAGATGCATTCGAGAACAAACGATACTGTTCGGCCGTATTCCTGGATGTCAAGCAGGCGTTCGACAGAGTGTGGCATCCTGGACTCCTCTACAAACTCAAGTCCCACCTTCCCAGTTCCCACTATGCCCTACTCAAATCGTATACTGAAGGAAGAGAGTTCCAAGTGCGATGCGGTTCCTCAACCAGCACGACAAGGCCTATACGAGCCGGAGTACCTCAAGGCAGCGTCCTTGGTCCCATCCTCTACACCCTGTTTACAGCAGACCTCCCTATCATACCCTCCCGTTACCTCACAGCAGCCACCTATGCAGATGACACGGCGTTCCTTGCCACCGCAACAAACCCTCAACTAGCATCAGCCATCATCCAGAGGCAACTGGATGCATTGGATCCATGGCTGAAACGCTGGAACATCGTGATCAACGCTGATAAATCCTCCCACACCACCTTCTCTCTGCGCAGAGGAGAATGCCCCCCGGTCTCACTCGACGGCGACACAATCCCTACCTCCAGCACCCCCAAATATTTAGGGCTGACCCTGGACAGAAGGCTGACTTGGGGCCCCCACATCAACAGAAAGCGTATCCAGGCCAACATACGCCTAAAGCAACTCCACTGGCTCATCGGTAAAAAGTCCAAGCTGCGAGAGAAACTAAAGATTCTCGTCTACAAGACTATTCTCAAGCCAATCTGGACGTACGGAATTCAGCTGTGGGGCACTGCAAGCACATCACATAGAAGGAAGATCCAGCGATTTCAAAACAGATGTTTGAGAATAGTCTCCAACGCCCATCCCTACCACGAAAATTCCGCCATCCACGAGGAGCTCGGGATTCCATGGGTAGACGACGAAATCTACAGACACAGTGTGAGATATGCTAGCAGACTGGAGAACCACCACAACCACCTGGCCGTCAACCTTCTAGACCATAGCCAATCCCTAAGACGCCTGCAGAGAACGCACCCGCTTGACCTTACTCAACATACTTAATCATACTTAACCCCTACCCAAGTACACTCGATGTACTCCCCTTAAGTT AATGTTTCCCTCCAAAAAATTTAATTATTGTCCACTAGGACAG*gggatgcttctgggcgtacc**actagt**cg agctacaacgactaccctcagt*-3’

##### G2/Jockey-3 DIG probe synthesis

500ng of pUCIDT-G2 was digested using SpeI and XhoI restriction enzymes in 1x Cutsmart Buffer for 1 hour at 37°C. The digest was run on a 1.0% SeaPlaque™ GTG™ agarose gel (Lonza) and a 1689bp band containing the *G2* sequence was gel extracted and purified using the PureLink™ Quick Gel Extraction Kit (Invitrogen). DIG labeled G2 probes were generated via PCR in 50μL reactions consisting 0.09ng of gel extracted *G2* DNA, 0.5μM of forward and reverse primers from the Universal_2 primer set (see Methods and Table S14: Universal Primers), 1x HF Buffer, 1 Unit of Phusion Polymerase (NEB), 0.2μM dGTP, 0.2μM dATP, 0.2μM dCTP, 0.15μM dTTP, 5nM DIG-dUTP (Roche). Probe was synthesized using the following cycles: 98°C for 30 seconds; 30 cycles of 98°C for 10 seconds, 72°C for 1 min, and a final extension at 72°C for 5 min. Unpurified PCR produce was used as a probe for FISH.

### Hi-C analysis

We used a publicly available Hi-C dataset from embryos (Gene Expression Omnibus accession number GSE103625) to provide additional support our candidate centromeric contigs [113]. We mapped Hi-C sequence reads to our assembly and processed the output with the HiC-Pro pipeline [114] to obtain informative valid interaction pairs (default parameters). We used a customized python script to count interactions between regions of interest and then normalized to the size of the regions (per 100 kb). To count interactions between different sized windows, we used BEDTools [115] to create windows of specified sizes across the assembly. We established the euchromatin-heterochromatin boundaries in our assembly based on previous studies. For chromosome 2, 3, X and Y, we transferred the euchromatin-heterochromatin boundary coordinates previously reported for *D. melanogaster* [116] to our assembly. For chromosome 4, we assigned the ~70 kb closest to the centromere in the assembled chromosome 4 as heterochromatin based one previously reported [117], and the rest of it as euchromatin. We then binned the genome into different regions based on their sequence content: centromere, centromere proximal heterochromatin, centromere distal heterochromatin, and euchromatin (Table S16). We then classified interactions between centromeric contigs and the different categories based on their genomic region (*e.g*. centromere to proximal heterochromatin, centromere to distal heterochromatin *etc.*). We reported the median count for each category and conducted data visualization and statistics in R.

We calculated the significance between different categories using a Kruskal-Wallis test by ranks with Dunn’s test for post-hoc analysis, and the pairwise Wilcoxon rank sum test with false discovery rate (FDR) correction [118] of type I error rates for multiple comparisons. We deem a result to be significant only if both tests agree.

### Phylogenetic analyses of IGS and *G2/Jockey-3* elements

We extracted all IGS elements from the genome using BLAST v2.7.1 [88] with parameters “task blastn -num_threads 24 -qcov_hsp_perc 90” and custom scripts. We extracted the *G2/Jockey-3* sequences based on Repeatmasker annotations and custom scripts. We aligned and manually inspected *G2/Jockey-3* and IGS alignments using Geneious v8.1.6 [119] (see Supplemental Files 12 and 13). We constructed maximum likelihood phylogenetic trees for *G2/Jockey-3* and IGS using RAxML v.8.2.11 with parameters “-m GTRGAMMA -T24 -d -p 12345 -# autoMRE -k -x 12345 -f a” [120]. We used the APE phylogenetics package in R [121] to plot the trees.

### *G2/Jockey-3* activity

We asked if *G2/Jockey-3* non-LTR retroelements have evidence for recent activity based on insertion polymorphism and expression. We examined RNAseq reads from testes for evidence of *G2/Jockey-3* because of the enrichment of these elements on the Y chromosome. We mapped poly-A [122] and total RNA [123] (Table S6) transcriptome data to our Repeat library using HISAT 2.1.0 [124] and estimated read depth of uniquely mapped read using samtools (depth -Q10; v1.7 [125]).

### Data availability

All supporting data associated with the manuscript is in the supplemental results. Supplemental files containing raw data and analyses are submitted to the dryad digital repository (doi:10.5061/dryad.rb1bt3j), Github (https://github.com/LarracuenteLab/Dmel.centromeres), and NCBI’s SRA under the bioproject PRJNA482653 and accessions SRR7588743 - SRR7588752.

