## Supplemental results for "Islands of retroelements are the major components of *Drosophila* centromeres"

**SUPPLEMENTAL REFERENCES** ..... **48**

#### Effects of library preparation

Centromeres are enriched in repetitive sequences, therefore CENP-A ChIPseq may be particularly sensitive to biases due to different library preparations. We carried out four replicate ChIPseq experiments: two with TruSeq (R1 and R4) and two with Accel-NGS 2S Plus (R2 and R3). The Accel-NGS 2S plus library kit requires less starting material, requires fewer PCR cycles, and is reported to lead to higher library complexity. Our mapping rate to the genome is >98% except for R1 (57%), which has bacterial read contamination from Brevundimonas. R2 and R3 have a higher reproduction rate in MACS ChIP peak calls than the R1 and R4 TruSeq libraries (Fig. S3; Table S4). To determine if the non-centromeric peaks were truly enriched in CENP-A, we used ChIP-qPCR. We did not detect an enrichment of CENP-A over a subset of the strongest non-centromeric peaks using qPCR (Fig. S5). We therefore think that most of these non-centromeric peaks represent noise from library bias rather than true signal. The signal-to-noise ratio in the TruSeq libraries was higher than the Accel libraries (Fig. S3). The CENP-A ChIP data from Talbert et al. 2018 [1] were obtained from independent sources (S2 cells and CID-GFP embryos), with a different ChIP method (*i.e.* MNase digestion) and library preparation. The results largely agree between these independent datasets (Table S4 and Fig. S3B).

Among centromeres and across replicates, Contig119 (cen4 candidate) and Y_Contig26 (cenY candidate) consistently appear in the top three contigs with the strongest signals in the IDR tests (Table S4). This is consistent with published cytological data showing the strongest CENP-A signal on the $4^{th}$ and Y chromosomes [2]. S2 cells lack a Y chromosome, therefore it is reassuring that we cannot detect cenY in the top 100 strongest signals in any S2 dataset. Outside of missing cenY, the CENP-A ChIP in S2 cells differs from the embryo ChIP, suggesting possible centromere rearrangements in S2 cells. We only detect weak signals from Contig79 (CenX candidate; *Maupiti*) in S2 cells and the MNase treated CENP-A ChIP data from

Talbert et al. [1] (Table S4 and Fig.S3B). We are unsure of the reasons for the weaker CenX signal in some datasets, however it is possible that our results are sensitive to ChIP and library preparation methods. Additionally, S2 cells have large-scale genomic rearrangements [3] that likely affect their centromeres.

#### Non-centromeric CENP-A associations

While most of the strongest CENP-A peaks in embryos are in our centromere islands, we do find significant peaks at other sites throughout the genome. The two replicate Accel libraries show far more peaks outside of the centromere islands than the two replicate TruSeq libraries (Table S4; Fig. S3A). However, many of the non-centromeric peaks were not reproducible between replicates using different library methods (Table S9). We therefore only focused on peaks overlapping in two of four replicate CENP-A ChIPseq experiments as potentially interesting sites (Table S4). We do not consistently find CENP-A enrichment on non-centromeric *G2/Jockey-3* elements (Table S9). Notably, some transposable elements not found on the centromere islands were consistently enriched in the CENP-A ChIP replicates (Fig 1C; Table S2)—among enriched non-centromeric sites are telomeric non-LTR retroelements and R1 elements. While many of these associations could represent non-specific peaks [4], there may be precedent for an association with the rDNA. The CENP-A assembly factor CAL1 [5] localizes at the nucleolus as well as at the centromere [6], therefore these reads could represent the fraction of CENP-A associated with nucleolar CAL1.

While functional kinetochores are unlikely to form on the regions of non-centromeric CENP-A enrichment, these sites might be poised for neo-centromere function. We hypothesized that a subset of the non-centromeric CENP-A-enriched regions would also be identified in CENP-A overexpression data. We therefore compared published CENP-A ChIP in CENP-A overexpression data from S2 cells [7] with our S2 data. Overall, there are only four contigs that overlap with our S2 data using standard peak-calling methods: cen2, cen3, one contig with *dodeca* and *Prodsat,* and one contig with mostly 1.688 satellite and a *Max* transposon (IDR ≤ 0.05; Table S17). The original study used custom peak calling analysis that binned regions to call large, lowly enriched peaks, which could explain these differences. Most of the regions of CENP-A enrichment in the S2 overexpression data were in contigs almost exclusively filled with *dodeca* (*e.g.* Contig86) and variants of the *1.688* family of repeats (*e.g.* Contig102 and 96). These contigs are proximal to canonical centromeres. Because *dodeca* is also part of the canonical 3$^{rd}$ chromosome centromere and the pericentromeric *1.688* repeats associate with the kinetochore-specific protein, BUB1, in prometaphase [8], these repeats may have some properties of centromeric DNA. In addition, both *dodeca* and *1.688* repeats can assume non-B form DNA structures[9]. We therefore hypothesize when CENP-A is overexpressed, the centromeres expand to include proximal pericentric heterochromatin and non-B form DNA [9, 10].

#### Centromere-associated repeat evolutionary genomics

##### Centromeric IGS

Centromeres are enriched for repetitive elements whose locations are not exclusive to centromeres (e.g. IGS and *G2/Jockey-3*). To determine if there are centromere-specific variants or if CENP-A recognizes and binds to a particular part of those repeats, we explored the evolutionary relationships between each repeat copy across the genome. The IGS$^{3cen}$ sequences represent a duplication from the IGS sequences that are spacers between the rDNA genes on the sex chromosomes. The IGS$^{3cen}$ repeats at cen3 and island with moderate CENP-A enrichment form their own distinct clade (Fig. S10; Supplemental File 14). The duplication event was happened near the divergence between *D. melanogaster* and the *Drosophila simulans* clade. However, $IGS^{3cen}$ is not found in the *Drosophila simulans* clade. We then infer that $IGS^{3cen}$ is a derived centromere and the expansion of $IGS^{3cen}$ is associated with the acquisition of its CENP-A binding ability.

***G2/Jockey-3***
We looked for orthologs of *G2* in other Drosophila species. We find orthologs of *G2* in the simulans clade, *D. erecta,* and *D. yakuba,* but not more distantly related species. Reciprocal best BLAST hits suggest that *G2* is a *Jockey*-type element orthologous to *Jockey-3* in *D. simulans*. *Jockey-3* is not annotated in *D. melanogaster* (according to Repbase), but RepeatMasker does annotate *Jockey-3* in our genome using the simulans *Jockey-3* consensus. We inferred the phylogeny of the 36 *Jockey-3* and 57 *G2* annotations >1 kb in *D. melanogaster* and the consensus of *Jockey-3* from *D. simulans, D. sechellia,* and *D. yakuba* using maximum likelihood methods (see Methods). Our phylogenetic analysis suggests that in *D. melanogaster G2* and *Jockey-3* are the same element (Fig. S3B and Supplemental file 15). Most of the *G2* elements are truncated at the 5' end. CENP-A does not pileup over a particular part of *G2* (based on the ratio of ChIP/Input across the full length of a *G2* element; Fig. 6). *D. melanogaster*'s version of *Jockey-3* contains an additional 1,102 bp with a predicted ORF upstream of the 5' end of *G2* annotations. We do not know what is required for *G2/Jockey-3* activity, we therefore define 'complete' elements as those containing intact ORFs encoding an endonuclease and a reverse transcriptase regardless of length and combine the 235 *G2* annotations and 63 *Jockey-3* annotations in our analyses giving us a total of 298 *G2/Jockey-3* partial and full-length elements. We believe that the first 487 bp of *D. simulans Jockey-3* consensus sequence is misannotated and instead belongs to a different repeat (see section on *D. simulans Jockey-3* below).

*G2/Jockey-3* elements are not randomly distributed in the genome, they are concentrated at centromeres [enrichment in centromeric contigs (63%) relative to other heterochromatin (31%; FET with FDR correction $P < 10^{-15}$; Table S9; Fig. 4; Fig. S7). We find evidence for *G2/Jockey-3* transcription among poly-A and total RNA-seq reads (Fig. S9; see Methods). *G2/Jockey-3* insertions segregate in populations of *D. melanogaster* [11]. Moreover, most *G2/Jockey-3* elements are truncated at the 5' end but are otherwise intact, which is a hallmark of non-LTR insertion (Supplemental File 13). Taken together, these results suggest that *G2/Jockey-3* are recently or currently active elements, consistent with a recent study of active elements in oogenesis [12]. The enrichment of *G2/Jockey-3* at centromeres is not a typical genomic distribution for recently active TEs. We plotted the distribution of other TEs enriched in CENP-A according to our ChIPseq analysis (*Jockey*-1; *G*; *Doc2*; *DM1731; R1; TART*), and inactive TE (*ProtoP*) (Fig. S11). Besides *G2/Jockey-3*, *G* (FET with FDR correction P = 0.013), *Doc2* (FET with FDR correction $P < 10^{-7}$) and *Jockey-1* (FET with FDR correction *P* = 0.013) are also enriched at centromeres compared to other heterochromatin regions (Table S9).

| **Location** | *G2/Jockey-3* | *G* | *Doc2* | *Jockey-1* | *DM1731* | *TART* | *ProtoP* |
|---|---|---|---|---|---|---|---|
| **Centromere** | 188 | 4 | 11 | 2 | 0 | 0 | 0 |

| | | | | | | | |
|---|---|---|---|---|---|---|---|
| **Pericentromeric heterochromatin** | 91 | 46 | 84 | 5 | 53 | 451 | 432 |
| **Other** | 19 | 8 | 9 | 0 | 6 | 49 | 68 |
| **P-value** (cen-het) | $<10^{-15}$ | 0.013 | $<10^{-7}$ | 0.013 | ns | 0.037* | 0.037* |
| **P-value** (cen-genome) | $<10^{-15}$ | 0.00013 | $<10^{-14}$ | 0.00079 | ns | ns | ns |

**Table S9. Statistical analysis of TE distributions**. We show the copy numbers of TEs in different genomic regions. The sum of base pairs in the assembly size in centromeres (432,440 bp), pericentromeric heterochromatin (37,089,066 bp) and other regions (118,457,213 bp) were used to compute the distribution statistics of TEs. We created a 2-by-2 contingency table for each TE comparing observed to expected (based on the sum of bp) for each comparison: centromere to heterochromatin (cen-het) regions or centromeres to whole genome (cen-genome). We computed a Fisher's exact test with FDR correction to get adjusted P values. *G2/Jockey-3*, *G*, *Doc2* and *Jockey-1* are significantly enriched in centromeres relative to other heterochromatic regions and to the whole genome. Asterisk signs show that *TART* and *ProtoP* are significantly underrepresented in centromeres relative to other heterochromatic regions.

#### Analysis of inter-centromere interactions by Hi-C

We divided the genome into the following regions: centromere, centromere proximal heterochromatin, centromere distal heterochromatin, and euchromatin. We compared intrachromosome interactions between centromeric contigs and its heterochromatin (proximal and distal) and euchromatin; and interchromosome interactions between that centromere and other chromosome's heterochromatin (proximal and distal) and euchromatin (Table S16). We tested for differences between interactions between genome compartments with Kruskall Wallis tests with a post-hoc Dunn's test and report P values adjusted based on false discovery rate (FDR)[13]. We find significant differences in the number of interactions between centromeres and interchromosome genomic regions (heterochromatin and euchromatin) and intrachromosome genomic regions (proximal heterochromatin, distal heterochromatin) (Kruskal Wallis $P<10^{-16}$; Fig S14). Among these genomic categories, we find more centromere to centromere interactions than centromere to distal-heterochromatin interactions (Dunn's $P_{adjusted}<10^{-11}$ for embryonic cycle 1–8, Dunn's $P_{adjusted}<10^{-3}$ for embryonic stage 16), centromere to inter-proximal heterochromatin interactions (Dunn's $P_{adjusted}=0.0088$ for embryonic cycle 1–8; Dunn's $P_{adjusted}=0.017$ for embryonic stage 16), centromere to inter-distal heterochromatin interactions (Dunn's $P_{adjusted}<10^{-8}$ for embryonic cycle 1–8; Dunn's $P_{adjusted}<10^{-4}$ for embryonic stage 16), and centromere to euchromatin interactions (Dunn's $P_{adjusted}<10^{-19}$ for embryonic cycle 1–8; Dunn's $P_{adjusted}<10^{-19}$ for embryonic stage 16). However, centromere to centromere interactions are not more abundant than centromere to proximal heterochromatin interactions (Dunn's $P_{adjusted}=0.59$ for embryonic cycle 1-8; Dunn's $P_{adjusted}=0.43$ for embryonic stage 16).

#### Localization of satellites on iso-1 mitotic chromosomes

We determined the position and organization of centromere-associated satellite DNA within the iso-1 line's genome using satellite specific FISH probes (see Methods and Tables S11–12) and an anti-CENP-C antibody to mark the centromeres (Fig. S7A–G). On the X chromosome, CENP-C overlaps with AATAT (100%, N = 27 spreads) and partially with AAGAG (56.3%, N = 16 spreads) (Fig. S7E–F). *SATIII* (100%, N = 34 spreads) (Fig. S12E) and a small block of AAGAT (26.7%, N = 30 spreads) (see adjusted signal inset on Fig. S12A) are adjacent to CENP-C on the X, thus they are categorized as pericentric. On chromosome 2, AAGAG (100%, N = 16 spreads) and *Prodsat* (100%, N = 12 spreads) overlaps with CENP-C (Fig. S7B). Chromosome 2 also contains two blocks of AATAG, with one small block overlapping with CENP-C (100%, N = 38 spreads) (Fig. S7C, G; white arrow) and one large block in heterochromatin (100%, N = 38 spreads) (Fig. S7C, G; yellow arrow). On chromosome 3, CENP-C overlaps with *dodeca* (100%, N = 12 spreads) (Fig. S7D) and is adjacent to *Prodsat* (100%, N = 12 spreads) (Fig. S7B, D, G). A pericentric block of AATAG can also be found on chromosome 3 (100%, N = 38 spreads) when FISH is performed without the *Prodsat* FISH probe (Fig. S7C). On chromosome 4, CENP-C overlaps with AAGAT (100%, N = 30 spreads) (Fig. S7A) and is flanked by AAGAG (100%, N = 16 spreads) (Fig. S7A–C, F) and AATAT (100%, N = 27 spreads) (Fig. S12E–F). On the Y, AATAT is the closest satellite to CENP-C (Fig. S7E–F), but it does not overlap with CENP-C. Refer to Table 1 in the main manuscript for the position of satellites within heterochromatin.

#### Localization of Oligopaint FISH probes on iso-1 mitotic chromosomes

We validated the positions of the $X^{Maupiti}$, $4^{Lampedusa}$, $Y^{Lipari}$, and $3^{Giglio}$ using Oligopaint FISH on mitotic spreads from the iso-1 line. $X^{Maupiti}$ probe signal overlapped with CENP-C and AAGAG on the X chromosome (100%, N = 31 spreads), recapitulating the organization seen in Contig 79 (Fig. S12A; white box). Interestingly, $X^{Maupiti}$ Oligopaint also overlaps with CENP-C, but not AAGAG on the Y chromosome (93.5%, N = 31 spreads) (Fig. S12A; yellow box). The $4^{Lampedusa}$ Oligopaint overlaps with CENP-C and is next to AAGAT (100%, N = 22 spreads) (Fig. S12B; white box). $4^{Lampedusa}$ also hybridized to the Y chromosome's centromere (100%, N = 22 spreads), which is devoid of AAGAT repeats confirming the identity of Contig119 as being centromere 4, which contains AAGAT (Fig. S12B; yellow box). The $Y^{Lipari}$ Oligopaint overlaps with CENP-C specifically on the Y chromosome (100%, N = 14 spreads) (Fig. S12C; white box). The closest satellite to the Y centromere is AATAT, however, this satellite does not overlap with CENP-C or $Y^{Lipari}$ (Fig. S12C; blue; white box). The $3^{Giglio}$ Oligopaint overlaps with CENP-C and *dodeca* on chromosome 3 (100%, N = 34 spreads) (Fig. S12D; white box) consistent with the composition of contig 3R_5. $3^{Giglio}$ also hybridizes to the rDNA on the X chromosome (38.2%, N = 34 spreads), away from CENP-C (Fig. S12D).

#### Localization of Oligopaint FISH probes on S2 cells mitotic chromosomes

All centromere islands (*e.g.* Fig. 2) are enriched in S2 cells (Table S4 and Fig. S3B) except for $Y^{Lipari}$, consistent with the Y chromosome being absent from S2 cells. However, we find some large differences between CENP-A domains in S2 cells and embryos. CENP-A is enriched over additional transposable elements (*e.g. Max-I*; Table S2) and strongly enriched in some simple tandem repeats (*e.g. Prodsat* and *AATAG*, Table S1) in S2 cells compared to the embryo ChIP-seq data using the same library method (R4).

We determined whether or not the positions of $X^{Maupiti}$, $Y^{Lipari}$, $3^{Giglio}$, and $4^{Lampedusa}$ in the iso-1 genome had been maintained in S2 cells using our Oligopaints for FISH and IF with an antibody for CENP-A to mark the centromere (Fig. S13A-D; green). DAPI staining combined with FISH revealed the S2 genome consisted of the canonical chromosomes X, 2, 3, and 4. In addition, S2 cells also contained a Robertsonian translocation between X and 4 (chromosome X;4), two centric fragment chromosomes generated most likely by a break within the centromere of chromosome 2 (designated cf(2R) and cf(2L)), and a small chromosome 4 (chromosome $4^s$) (See Fig. S8 and Table S20). The $X^{Maupiti}$ Oligopaint overlaps with CENP-A on the X;4 chromosome and is adjacent to CENP-A on the X chromosome, indicating $X^{Maupiti}$'s position on X;4 is conserved with iso-1, but CENP-A may have shifted on the normal X chromosome. Interestingly, the $X^{Maupiti}$ Oligopaint also hybridizes to the centromere of chromosome 3 (see "Signal Adjusted" panel) and other heterochromatic loci on the X, chromosomes 2, cf(2R), cf(2L), and 4 (Fig. S13A), suggesting that transposable elements within $X^{Maupiti}$ may have remobilized in S2 cells. The $4^{Lampedusa}$ Oligopaint hybridizes to loci adjacent to CENP-A on cf(2L), the X and X;4 chromosomes, and is heterochromatic on chromosomes X, X;4, 2, cf(2R), 3, and 4 (Fig. S13B). Surprisingly, the $4^{Lampedusa}$ Oligopaint overlaps with CENP-A on chromosomes 2, cf(2R), and 3, but not on chromosomes 4 and $4^s$ (Fig. S13B). These data suggest that like $X^{Maupiti}$, $4^{Lampedusa}$ elements have also remobilized and integrated into these chromosomes. On normal chromosome 4 the signals for $4^{Lampedusa}$ and that of CENP-C do not overlap, so it is possible that the centromere has repositioned. In the case of chromosomes $4^s$, the lack of $4^{Lampedusa}$ centromeric signal suggests that the centromere was lost, perhaps via a translocation with chromosome X;4. Unlike $X^{Maupiti}$ and $4^{Lampedusa}$, the $Y^{Lipari}$ Oligopaint shows no hybridization to anywhere in the S2 genome (Fig. S13C), consistent with the ChIP-seq results. Lastly, the $3^{Giglio}$ Oligopaint hybridizes to the centromere and pericentromere of chromosome 3, as well as the heterochromatic locus associated with rDNA of the X and X;4 chromosomes (Fig. S13D), indicating that $3^{Giglio}$'s position has been retained between iso-1 and S2 cells (Fig. S12D). Additionally, the pericentric position of $3^{Giglio}$ suggests that the CENP-A domain on a subset of chromosome 3's has shifted. See Table S18 for quantification of Oligopaint hybridization locations in S2 cells.

| | | Chromosome | | | | | | | |
|---|---|---|---|---|---|---|---|---|---|
| Probe | Location | X | X;4 | 2 | cf(2R) | cf(2L) | 3 | 4 | 4s |
| AATAT | C | 0.0 | 15.0 | 10.0 | 5.9 | 5.6 | 0.0 | 85.0 | 84.2 |
| | P | 100.0 | 100.0 | 90.0 | 47.1 | 61.1 | 45.0 | 100.0 | 100.0 |
| | H | 0.0 | 0.0 | 100.0 | 76.5 | 66.7 | 100.0 | 100.0 | 100.0 |
| | N | 20 | 20 | 20 | 17 | 18 | 20 | 20 | 19 |
| AAGAG | C | 0.0 | 14.3 | 64.3 | 72.7 | 69.2 | 0.0 | 23.1 | 0.0 |
| | P | 100.0 | 92.9 | 100.0 | 100.0 | 100.0 | 0.0 | 84.6 | 91.7 |
| | H | 0.0 | 7.1 | 100.0 | 100.0 | 92.3 | 100.0 | 100.0 | 0.0 |
| | N | 14 | 14 | 14 | 11 | 13 | 14 | 13 | 12 |
| AATAG | C | 0.0 | 0.0 | 100.0 | 69.2 | 71.4 | 0.0 | 0.0 | 0.0 |
| | P | 5.9 | 5.9 | 100.0 | 92.3 | 100.0 | 100.0 | 5.9 | 0.0 |
| | H | 0.0 | 0.0 | 100.0 | 53.8 | 100.0 | 0.0 | 0.0 | 0.0 |
| | N | 17 | 17 | 17 | 13 | 14 | 17 | 17 | 17 |
| AAGAT | C | 7.1 | 7.1 | 0.0 | 0.0 | 0.0 | 0.0 | 0.0 | 0.0 |
| | P | 100.0 | 92.9 | 92.9 | 72.7 | 53.8 | 0.0 | 15.4 | 8.3 |
| | H | 0.0 | 0.0 | 92.9 | 72.7 | 38.5 | 0.0 | 84.6 | 0.0 |
| | N | 14 | 14 | 14 | 11 | 13 | 14 | 13 | 12 |
| *dodeca* | C | 0.0 | 0.0 | 0.0 | 0.0 | 0.0 | 100.0 | 0.0 | 0.0 |
| | P | 0.0 | 0.0 | 0.0 | 0.0 | 0.0 | 100.0 | 0.0 | 0.0 |
| | H | 0.0 | 0.0 | 0.0 | 0.0 | 0.0 | 0.0 | 0.0 | 0.0 |
| | N | 18 | 18 | 18 | 18 | 18 | 18 | 18 | 18 |
| *Prodsat* | C | 0.0 | 0.0 | 100.0 | 33.3 | 27.8 | 100.0 | 0.0 | 0.0 |

| | | | | | | | | | |
|---|---|---|---|---|---|---|---|---|---|
| | P | 0.0 | 0.0 | 100.0 | 46.7 | 100.0 | 100.0 | 0.0 | 0.0 |
| | H | 0.0 | 0.0 | 0.0 | 0.0 | 0.0 | 0.0 | 0.0 | 0.0 |
| | N | 18 | 18 | 18 | 15 | 18 | 18 | 18 | 18 |
| *SATIII* | C | 35.0 | 0.0 | 0.0 | 0.0 | 0.0 | 0.0 | 0.0 | 0.0 |
| | P | 100.0 | 100.0 | 0.0 | 0.0 | 0.0 | 5.0 | 0.0 | 0.0 |
| | H | 100.0 | 100.0 | 0.0 | 0.0 | 0.0 | 100.0 | 0.0 | 0.0 |
| | N | 20 | 20 | 20 | 20 | 20 | 20 | 20 | 20 |
| $X^{Maupiti}$ | C | 23.3 | 83.3 | 0.0 | 0.0 | 0.0 | 93.3 | 3.3 | 3.3 |
| | P | 100.0 | 100.0 | 0.0 | 43.3 | 10.0 | 16.7 | 3.3 | 0.0 |
| | H | 100.0 | 100.0 | 100.0 | 26.7 | 16.7 | 10.0 | 80.0 | 13.3 |
| | N | 30 | 30 | 30 | 30 | 30 | 30 | 30 | 30 |
| $Y^{Lipari}$ | C | 0.0 | 0.0 | 0.0 | 0.0 | 0.0 | 0.0 | 0.0 | 0.0 |
| | P | 0.0 | 0.0 | 0.0 | 0.0 | 0.0 | 0.0 | 0.0 | 0.0 |
| | H | 0.0 | 0.0 | 0.0 | 0.0 | 0.0 | 0.0 | 0.0 | 0.0 |
| | N | 35 | 35 | 35 | 35 | 35 | 35 | 35 | 35 |
| $3^{Giglio}$ | C | 0.0 | 0.0 | 0.0 | 0.0 | 0.0 | 100.0 | 0.0 | 0.0 |
| | P | 0.0 | 0.0 | 0.0 | 0.0 | 0.0 | 0.0 | 0.0 | 0.0 |
| | H | 100.0 | 100.0 | 0.0 | 0.0 | 0.0 | 0.0 | 0.0 | 0.0 |
| | N | 25 | 25 | 25 | 25 | 25 | 25 | 25 | 25 |
| $4^{Lampedusa}$ | C | 20.0 | 6.7 | 73.3 | 46.7 | 13.3 | 93.3 | 0.0 | 0.0 |
| | P | 86.7 | 93.3 | 93.3 | 60.0 | 73.3 | 66.7 | 0.0 | 0.0 |
| | H | 100.0 | 100.0 | 100.0 | 73.3 | 93.3 | 66.7 | 80.0 | 0.0 |
| | N | 15 | 15 | 15 | 15 | 15 | 15 | 15 | 15 |

**Table S18: S2 cell FISH quantification.** Percentage of probe signals that overlap with different cytological locations (C: Centromere; P: Pericentromere; H: Heterochromatin and N: number of spreads analyzed) in S2 cells.

#### Localization of satellites on S2 cells mitotic chromosomes

To determine whether or not the positions of centromere associated satellite sequences have been maintained from iso-1 flies to S2 cells, we performed FISH for centromeric satellites on S2 mitotic chromosomes using an anti-CENP-A antibody to mark the centromere (Fig. S8; green). Consistent with the X chromosome of iso-1 (Fig. S7E–F), the centromeres of the X and X;4 chromosomes are flanked by *SATIII* on the long arm and AATAT (Fig. S8A), and AAGAG (Fig. S8D–F), and AAGAT (Fig. S8F) on the short arm. However, AAGAT, which is only found on chromosomes 2 and 4 in iso-1 flies (Fig. S7A), is also found on the short arm of chromosomes X and X;4 (Fig. S8F). On chromosome 2 CENP-A overlaps with AAGAG (Fig. S8D–F), AATAG (Fig. S8C, E), and *Prodsat* (Fig. S8B–D), consistent with iso-1's chromosome 2 (Fig. S12B–C, G), but has also gained AATAT repeats in all chromosome 2's (Fig. S8A) and has lost AAGAT on all but one chromosome 2 (Fig. S8F). The centromere of cf(2R) overlaps with AAGAG (Fig. S8B) and AATAG, with a small block of *Prodsat* adjacent to CENP-A (adjusted signal inset of Fig. S8C). The centromere of cf(2L) overlaps with AAGAG and *Prodsat* (Fig. S8B–D) and is flanked by two blocks of AATAG (Fig. S8C). Blocks of AATAT (see cf(2R) and cf(2L) in Fig. S8A) and AAGAT (see cf(2R) in Fig. S8F) are in the pericentric heterochromatin of cf(2R) and cf(2L). As is seen on iso-1 chromosome 3's, the centromere of the S2 chromosome 3 overlaps with *dodeca* (Fig. S8B) and *Prodsat* (Fig. S8B–D) and contains a pericentric block of AATAG only visible when not combined with *Prodsat* FISH probes (see adjusted signal inset on Fig. S8E). Chromosome 4 and $4^{s}$'s centromeres overlap with AATAT (Fig. S8A), contain AAGAG within the pericentric heterochromatin (Fig. S8E, F), and AAGAT is present on the heterochromatin of chromosome 4 but absent on chromosome $4^s$ (Fig. S8F). One explanation for the loss of AAGAT on S2 chromosome $4^s$ is that the AAGAT on X;4 was inherited from the S2 chromosome $4^s$ via a Robertsonian translocation. As for chromosome 4, the movement of AAGAT away from the centromere could have occurred via a pericentric inversion with breakpoints in AATAT and AAGAT, which either left behind the original centromere surrounded by an amount of AAGAT undetectable by FISH or inactivated the endogenous centromere within AAGAT and activated a new centromere within AATAT. See Supplementary Tables 18–19 for detailed quantification of all satellite locations.

| Satellite | Cen | Peri | Het |
|---|---|---|---|
| AATAT | 4, $4^s$ | X, X;4, 2, cf(2R), cf(2L) | 3 |
| AAGAG | 2 | X, X;4, cf(2R), cf(2L), 4, $4^s$ | 3 |
| AATAG | 2, cf(2R) | 2, cf(2R), cf(2L), 3 | |
| AAGAT | | X, X;4, 2, cf(2R) | 4 |
| *dodeca* | 3 | | |
| *Prodsat* | 2, 3 | cf(2R), cf(2L) | |
| *SATIII* | | X, X;4 | 3 |

**Table S19: S2 cell satellite locations.** Locations of satellite repeats determined by IF/FISH on S2 cell chromosomes X, X;4 (Robertsonian translocation between chromosomes X and 4), 2, cf(2R) (centric fragment of chromosome 2R), cf(2L) (centric fragment of chromosome 2L), 3, 4, and $4^s$ (small chromosome 4), using an anti-CENP-A antibody to mark the centromere. Locations were designated as centromeric (Cen), pericentric (Peri), or heterochromatic (Het).

#### *D. simulans* centromeric *G2/Jockey-3*

A repeat identified by Talbert et al. [1] as enriched in CENP-A has homology to the first 487 bp of the *D. simulans Jockey-3* annotation in Repbase (500U). However, no other Drosophila species *Jockey-3* element contains this 500-bp fragment, including *G2/Jockey-3* in *D. melanogaster*. Because we find that *Jockey-3* is significantly enriched in CENP-A, including sequences 3' to the first 487-bp with homology to the centromeric satellite (Fig. 6), we suspect that the Repbase consensus is misannotated. The chimeric annotation may come from *Jockey-3* elements having an insertion preference for the 500-bp satellite, or selection for centromeric insertions of the TE. We cannot distinguish between these possibilities, but a detailed assembly-based method is forthcoming. We therefore refer to this element, less the first 487-bp found in the Repbase annotation [14] (https://www.girinst.org/protected/repbase_extract.php?access=Jockey-3_DSim; last accessed 1/15/2019), as *G2/Jockey-3* and use the *D. simulans* consensus of this element for read mapping in Fig 6B.

### List of Supplemental Figures

Figure S1. Enrichment of simple tandem repeats enriched in CENP-A ChIP-seq across four replicates.
Figure S2. G2 and Jockey-3 belong to the same repeat family.
Figure S3. Enrichment of CENP-A ChIP-seq data in embryos and S2 cells.
Figure S4. CENP-A occupies DNA sequences within putative centromere contigs
Figure S5. ChIP-qPCR validation of CENP-A-enriched regions
Figure S6. Relative depth of Pacbio reads across centromeric contigs.
Figure S7. Satellite FISH on iso-1 larval brain mitotic spreads
Figure S8. Satellite FISH on S2 cell mitotic spreads
Figure S9. Transcription of *G2/Jockey-3* elements.
Figure S10. Relationship of IGS in *D. melanogaster* and closely related species of the simulans clade (*D. simulans* and *D. sechellia*) and *D. yakuba*.
Figure S11. Genomic TE distribution across chromosomes.
Figure S12. Oligopaint FISH on larval brain mitotic spreads from iso-1 flies.
Figure S13. Oligopaint FISH on S2 cell mitotic spreads. .
Figure S14. Quantification of interactions between centromeres and different genomic regions by Hi-C.
Figure S15. Calibration of extended chromatin fiber stretching.
Figure S16. Organization of the X centromere.
Figure S17. Organization of centromere 4.
Figure S18. Organization of the Y centromere.
Figure S19. Organization of centromere 3.
Figure S20. Tracking of longer centromere 3 fibers reveals a second region containing CENP-A on *dodeca*.
Figure S21. Organization of the Centromere 2.

### List of Supplemental Tables and their location

Table S1. Enrichment of simple tandem repeats in kseek analyses.
Table S2. Raw and normalized counts of reads mapped to the complex repeats.
Table S3. ChIPtigs with MACS peaks.
Table S4. MACS peaks called by mapping to the genome assembly.
Table S5. IDR tests between different replicates from OreR ChIP-seq.
Table S6. Summary of all sequencing datasets used in this study.
Table S7. List of qPCR Primers (in methods).
Table S8. Non-centromeric overlapping MACS peaks in the OreR embryo ChIP replicates.
Table S9. Statistical analysis of TE distributions (in supplemental results).
Table S10. Oligopaint hybridization conditions (in methods).
Table S11. Labeled Satellite Probes (in methods).
Table S12. Unlabeled satellite probes.
Table S13. Secondary Oligo Probes.
Table S14. Universal primers.
Table S15. Sub-library-specific primers.
Table S16. Chromatin status assignments for contigs.
Table S17. Overlap between normal and CENP-A overexpression S2 cells.
Table S18. S2 cell FISH quantification
Table S19. S2 cell satellite locations.

#### Supplementary figures

**Figure S1. Enrichment of simple tandem repeats in CENP-A ChIP-seq across four replicates.** Plot of normalized CENP-A/Input for simple tandem repeats for each ChIP-seq replicate, sorted by median (red lines). Shown are only the simple tandem repeats with median CENP-A/Input > 1 in all four CENP-A ChIP replicates (see details in Table S1). The simple tandem repeats with less than 10 counts of input reads in any one replicate are not shown.

**Figure S2. *G2* and *Jockey-3* correspond to the same Non-LTR retroelement.** A maximum likelihood phylogenetic tree showing the relationship between *G2* and *Jockey-3* sequences in *D. melanogaster* genome and closely related species in the simulans clade (*D. simulans* and *D. sechellia*) and *D. yakuba*. In *D. melanogaster*, *G2* and *Jockey-3* are interleaved across the phylogeny, and thus likely correspond to the same repeat type. We therefore refer to these elements collectively as *G2/Jockey-3* throughout the manuscript.

**Figure S3. Reproducibility of CENP-A ChIP enrichment among replicates in embryos and S2 cells.** Locations of the top 100 strongest peaks for each ChIP experiment. **A)** Plot of the location of top 100 strongest peaks for each ChIP experiment on the diagonal (see details in Table S4). For the four replicate ChIP experiment in our OreR embryos, we examined the reproducibility of our experiments by first applying the IDR (Irreproducible Discovery Rate) test and only keeping peaks with IDR≤0.05. The number of these peaks are plotted below the diagonal. Between Replicates 2 and 3, we found a total of 16,870 overlapping peaks, but 16,833 were weakly enriched relative to the overlapping peaks between other datasets because they are technical repeats with a shared library bias (Accel, see Supplemental Methods). We therefore only report the 37 strongest peaks (the average peak number of other comparisons between replicates). The IDR dataset comparisons are in Table S5. We show the correlation between the CENP-A ChIP replicates above the diagonal. Plotted are the signal strength after IDR tests (normalized ChIP over input ratio from 1-1,000 on a log10 scale) with Spearman's rho. The five contigs with the most consistent peaks within and among replicates correspond to the five centromeric candidates. **B)** Plot of ChIP-seq data from S2 cells (this paper, Talbert et al. 2018, and Chen et al. 2015) and an independent embryo CID-GFP ChIP-seq dataset (see details in Table S4; Talbert et al. 2018; 5m and 15m represent different MNase treatments). The centromeric contigs are also CENP-A enriched in these independent datasets, with the exception of the X chromosome centromere contig. S2 cells lack a Y and are therefore not expected to have peaks on the Y candidate centromere contig.

**Figure S4. CENP-A occupies DNA sequences within putative centromere contigs.** Organization of each CENP-A enriched island corresponding to centromere candidates: **A)** X centromere, **B)** centromere 4; **C)** Y centromere; **D)** centromere 3; **E)** centromere 2. Different repeat families are color coded (see legend; note that *Jockey* elements are shown in one color even though they are distinct elements). The normalized CENP-A enrichment over input (plotted on a log scale) is shown for three replicates (Replicate 2 is in Fig. 2) colored in gray for simple repeats and black for complex island sequences. While the mapping quality scores are high in simple repeat regions, we do not use these data to make inferences about CENP-A distribution (see text for details). The coordinates of the significantly CENP-A-enriched ChIPtigs mapped to these contigs (black) and the predicted ChIP peaks (orange) are shown below each plot.

**Figure S5. ChIP-qPCR validation of CENP-A-enriched regions. A)** Diagram showing putative centromere contigs showing the locations of CENP-A ChIPtigs in black and CENP-A MACS peaks in orange as in Figure 2. Locations of contig-specific qPCR primer binding sites are shown by magenta arrows. **B)** Graph showing our ChIP-qPCR results using these primers. The enrichment is calculated relative to the input and is normalized by the *RpL32* promoter region as a noncentromeric control. **C)** Graph showing our ChIP-qPCR results using primers targeting other regions that showed CENP-A enrichment, but that were not in our contigs. Again, the enrichment is calculated relative to the input and is normalized by *RpL32* promoter as a noncentromeric control. We did not observe a robust CENP-A enrichment at these sites.

**Figure S6. Relative depth of Pacbio reads across centromeric contigs.** Pacbio reads were mapped to the genome using Minimap (v 2.11) and the parameter "-ax map-pb.". Shown are **A)** X centromere, **B)** centromere 4; **C)** Y centromere; **D)** centromere 3; **E)** centromere 2. The depth of only the high-quality mapped reads (mapped Q ≥ 30) was estimated for each position and normalized by the median depth of other genomic regions (98.32x for autosomes and 49.16x for sex chromosomes) to get relative depth. The relative depth of the TE-rich islands are close to 1, whereas the depth of the flanking simple satellites are uneven with some regions >1 and some <1. We therefore exclude simple repeats from any assembly-based analyses and color these regions gray in Fig. 2 and Fig. S4 to indicate that caution should be used in interpreting these regions of the assembly.

**Figure S7. Satellite FISH on iso-1 larval brain mitotic spreads.** IF/FISH using an anti-CENP-C antibody (green) and satellite FISH probes in the following combinations: **A)** AAGAT (magenta), and AAGAG (blue), with a high contrast inset of AAGAT on the X chromosome; **B)** *Prodsat* (magenta) and AAGAG (blue); **C)** AATAG (magenta) and AAGAG (blue), with AATAG blocks identified by white (small block) and yellow (large block) arrows; **D)** *Prodsat* (magenta) and *dodeca* (blue); **E)** AATAT (magenta) and *SATIII* (blue); **F)** AATAT (magenta) and AAGAG (blue); **G)** AATAG (magenta) and *Prodsat* (blue), with AATAG blocks identified by white (small block) and yellow (large block) arrows. DAPI is shown in gray. Bar 5µm

**Figure S8. Satellite FISH on S2 cell mitotic spreads.** IF-FISH using an anti-CENP-A antibody (green) and satellite FISH probes in the following combinations: **A)** AATAT (magenta) and *SATIII* (blue); **B)** *dodeca* (magenta) and *Prodsat* (blue); **C)** AATAG (magenta) and *Prodsat* (blue), with a high contrast inset of AATAG and *Prodsat* on cf(2R); **D)** AAGAG (magenta) and *Prodsat* (blue); **E)** AAGAG (magenta) and AATAG (blue), with a high contrast inset of AATAG on chromosome 3; **F)** AAGAT (magenta) and AAGAG (blue). DAPI is shown in gray. Bar 5 µm.

**Figure S9. Transcription of *G2/Jockey-3* elements.** A) Shown is the plot of the normalized reads depth from uniquely mapped reads (mapping quality ≥ 10) across the *G2/Jockey-3* consensus element obtained from mapping total and poly-A RNA-seq data from testes [15, 16] to our repeat library. B) Quantitative RT-PCR analysis of total RNA extracted from three independent overnight embryo collections. Expression levels were compared to the negative control gene Mst84Da (testis-specific). Cen X, 4 and 3, but not Y and 2, show low levels of transcription compared to the housekeeping gene Actin. Although the primers (Table S7) are specific for each centromere, the primer sets could amplify *G2/Jockey-3* copies not included in our assembly. Error bars=SD.

**Figure S10. Relationship of IGS in *D. melanogaster* and closely related species of the simulans clade (*D. simulans* and *D. sechellia*) and *D. yakuba*.** Maximum likelihood phylogenetic tree of all individual IGS sequences found in the *D. melanogaster* genome with related outgroups. Node support is only shown for key nodes in the tree (complete tree is in supplemental file 14). All centromeric IGS sequences appear to have a single origin: they duplicated from sex-linked IGS interspersed at the rDNA loci at some time near the divergence of the simulans clade and *D. melanogaster*. IGS repeats in blue (extra) are similar to the IGS at the cen 3 island, but are on small contigs of unknown origin, one of which is moderately enriched in CENP-A. We refer to all black and blue IGS repeats as $IGS^{3cen}$.

**Figure S11. Genomic TE distribution across chromosomes.** Distribution of TEs (represented by different colors) along the following chromosomes: A) chromosome 2; B) chromosome 3; C) chromosome 4; D) chromosome X; and E) chromosome Y. Contigs from each chromosome were concatenated in order with an arbitrary insertion of 100 kb of 'N'. Distances along the x-axis are approximate. The order and orientation of the Y chromosome contigs is based on gene order (see Chang and Larracuente 2018). Each triangle corresponds to one TE, where filled shapes indicate full length TEs and open shapes indicate truncated TEs. The vertical grey bars represent the arbitrary 100kb window inserted between contigs, indicating where we have gaps in our assembly. The centromere positions are set to 0 for each chromosome. Chromosomes are not drawn to scale (chromosome 4 and Y are enlarged). We show the genomic distribution of a sample of TEs enriched in CENP-A according to our ChIPseq analysis (all except *PROTOP*). *PROTOP* are DNA transposons that have not been recently active and their distribution is primarily in heterochromatin. *TART* elements are non-LTR retroelements highly enriched at telomeres, and are also moderately CENP-A enriched. *DM1731* is a retroelement moderately enriched for CENP-A, but not enriched in the centromere islands. *DOC2*, *G*, *Jockey-1* and *G2/Jockey-3* are CENP-A enriched non-LTR retroelements abundant in the centromere islands (see Table S2 and Table S9).

**Figure S12. Oligopaint FISH on larval brain mitotic spreads from iso-1 flies.** IF-FISH using an antibody for CENP-C (green), centromere Oligopaint FISH probes (magenta), and FISH probes for centromeric satellites (blue) in the following combinations: **A)** *Maupiti* (X; magenta) and AAGAG (blue); **B)** *Lampedusa* (4; magenta) and AAGAT (blue); **C)** *Lipari* (Y; magenta) and AATAT (blue); **D)** *Giglio* (3; magenta) and *dodeca* (blue). White boxes show the separate signals at the targeted centromeres. Yellow boxes show centromeric hybridizations at other centromeres. DAPI is shown in gray. Bar 5µm.

**Figure S13. Oligopaint FISH on S2 cell mitotic spreads.** IF/FISH using an antibody for CENP-A (green) and centromere oligopaint FISH probes designed to target centromere contigs (magenta). **A)** *Maupiti* (X); **B)** *Lampedusa* (4); **C)** *Lipari* (Y); **D)** *Giglio* (3). The "Signal Adjusted" panels for **A** and **B** show high contrast Oligopaint hybridization for visualization of weak foci. Bar 5µm.

**Figure S14. Quantification of interactions between centromeres and different genomic regions by Hi-C.** Plots showing intra- and inter-chromosomal interactions between regions in Hi-C data from: A) stage 16 embryos (end of embryogenesis) and B) embryonic cycles 1–8 (before zygotic genome activation; data from Ogiyama et al. 2018). The different colors indicate interactions with individual centromeres of all chromosomes. Centromere-centromere interactions are significantly more frequent than interactions between centromeres and distal heterochromatin, inter distal heterochromatin and euchromatin, and marginally more significant than centromere-inter proximal heterochromatin interactions. **** adjusted $P<0.0001$; * adjusted $P<0.02$, Pairwise Wilcoxon rank sum test with false discovery rate (FDR) correction; Kruskal-Wallis test by ranks with Dunn's test for post-hoc analysis.

**Figure S15. Calibration of extended chromatin fiber stretching.** Stretched chromatin fibers from female 3rd instar larval brain cells using the following probes: **A)** *Rsp* locus (heterochromatic; ~100kb; green); **B)** 100kb Oligopaint for a heterochromatic region on chromosome 3L (80C4; magenta); **C)** 100kb Oligopaint for a euchromatic region ~600kb from the telomere of chromosome 3L (61C7; cyan). Arrows show the region of the fiber that was measured. Bar 5µm. **D)** Scatter plot showing the quantification of fiber lengths. Mean lengths were used to estimate the size in kb (~10kb/1µm). Error bars show the standard deviation. P>0.09 (n.s.) for each pair of measurements compared (two-tailed t-test).

**Figure S16. Organization of the X centromere. (A-G)** Examples of fibers visualized with IF with anti-CENPA antibody (green), FISH with Oligopaints for *Maupiti* (magenta), and a AAGAG probe (cyan) on female 3rd instar larval brain cells. DAPI is shown in gray. CENPA occupies *Maupiti* and the AAGAG satellite. We observed some variation in FISH signals and *Maupiti* and CENP-A domain lengths, likely due to the efficiency of oligopaint binding and variable stretching in this region. Arrows show the region of the fiber that was measured. **H)** Scatter plot showing the quantification of the length of *Maupiti* FISH and CENP-A IF signals. Error bars show the standard deviation. N=24 fibers. Bar 5µm.

**Figure S17. Organization of centromere 4**. **(A-G)** Examples of fibers visualized by IF with anti-CENPA antibody (green), FISH Oligopaint FISH for *Lampedusa* (magenta), and a AAGAT probe (cyan). DAPI is shown in gray. CENP-A occupies predominantly the island *Lampedusa*. Arrows show the region of the fiber that was measured. **H)** Scatter plot showing the quantification of the length of *Lampedusa* FISH and CENP-A IF signals. Error bars show the standard deviation. N=25 fibers. Bar 5µm.

**Figure S18. Organization of the Y centromere. (A-E)** Examples of fibers visualized by IF with anti-CENPA antibody (green), FISH with Oligopaints for *Lipari* (magenta). DAPI is shown in gray. We did not include satellite FISH because no centromeric satellites are known for the Y. Note that the oligopaints only target to part of *Lipari* (see Figure 4). CENP-A is observed occupying sequences beyond the oligopaint region, likely over the remaining part of the island. Arrows show the region of the fiber that was measured. **F)** Scatter plot showing the quantification of the length of *Lipari* FISH and CENP-A IF signals. Error bars show the standard deviation. N=19 fibers. Bar 5µm.

**Figure S19. Organization of centromere 3. A-E)** Examples of fibers visualized by IF with anti-CENP-A antibody (green), FISH with Oligopaints for *Giglio* (magenta), and a probe for the centromere 3 specific *dodeca* satellite (cyan). DAPI is shown in gray. CENP-A occupies primarily *Giglio*, and a small stretch of *dodeca* satellite. Note that the binding of the *dodeca* (an LNA probe) is quite variable between fibers and results in several gaps that could be a result of the higher stringency conditions needed for *Giglio* Oligopaint FISH. Arrows show the region of the fiber that was measured. **F)** Scatter plot showing the quantification of the length of *Giglio* FISH and CENP-A IF signals. Error bars show the standard deviation. N=30 fibers. Bar 5µm.

**Figure S20. Tracking of longer centromere 3 fibers reveals a second region containing CENP-A on *dodeca*. (A-D)** Examples of longer fibers tracked along *dodeca* from the experiment in Figure 21, visualized by IF with with anti-CENPA antibody (green), Oligopaint FISH for *Giglio* (magenta), and FISH with *dodeca* probe (cyan). DAPI is shown in gray. Note the presence of *Giglio* signal on the *dodeca* CENP-A region. Multiple, overlapping panels were often acquired to follow an individual fiber. Panels were then cropped and juxtaposed in the figure, with white lines showing the separate images. White boxes show the CENP-A domain on *Giglio,* yellow boxes show the smaller domain on *dodeca*. N=5 (these are rare fibers to find in our preparations due to their length). Bar 5µm.

**Figure S21. Organization of the Centromere 2. (A-D)** Examples of fibers visualized with IF with anti-CENP-A antibody (green) and FISH with satellites. DAPI is shown in gray. **(A-B)**

Examples of fibers showing colocalization of CENP-A (green) with *Prodsat* (magenta) and AATAG (cyan). **(C-D)** Examples of fibers with AAGAG (cyan) and *Prodsat* (magenta). **E)** Example of fiber with AAGAG (magenta) and AATAG (cyan). We propose that *Capri* is located between flanking blocks of AAGAG and AATAG satellites that reside very close to where the *Prodsat* begins. Arrows show the region that was measured for each fiber. **F)** Scatter plot of CENP-A IF signals lengths. **G)** Model for the organization of centromere 2 showing a possible location of *Capr*i. Error bars show the standard deviation. N= 18 fibers. Bar 5µm.

### Legends for large Tables (Supporting Information)

**Table S1. Enrichment of simple tandem repeats in kseek analyses**. We used kseek [17] to estimate read counts for each kmer, and normalized these read counts using the total mapped reads for each dataset (ChIP and input). We identified CENP-A-enriched kmers using the ratio of normalized counts for each ChIP experiment and its corresponding input. The enriched kmers reflect simple tandem repeats enriched in CENP-A discussed in the main text and Figure S1. Figure 1 summarizes kmers with satellite repeats associated with centromeres.

**Table S2. Raw and normalized counts of reads mapped to the complex repeats.** Rows correspond to complex repeat families (TEs and complex satellites), with the counts per family in the ChIP and input reads from every dataset. We calculated enrichment for each repeat type by normalizing by total mapped reads for each dataset and taking the ratio of normalized values for each ChIP and its corresponding input.

**Table S3. ChIPtigs with MACS peaks.** We mapped all ChIP-seq data to the *de novo* assembled ChIPtigs and called peaks using MACS with high-quality reads (mapping quality ≥ 30 and masked PCR duplicates). We also mapped ChIPtigs to the genome to determine its genomic location and assigned repeat IDs based on BLAST results.

**Table S4. MACS peaks called by mapping to the genome assembly.** We mapped the ChIP and input reads to our genome assembly and used the high-quality reads (mapping quality ≥ 30 and masked PCR duplicates) to call ChIP peaks with MACS. We show the peak locations for each dataset.

**Table S5. IDR tests between different replicates from OreR ChIP-seq.** We used IDR to compare MACS peaks from different ChIP-seq replicates. We show the statistics for shared peaks from each comparison.

**Table S6. Summary of all sequencing datasets used in this study.** We list reads and mapping summaries of all Illumina and long read datasets generated in this paper or downloaded from NCBI's SRA.

**Table S8. Non-centromeric overlapping MACS peaks in the OreR embryo ChIP replicates.** We listed peaks outside canonical centromeres with any agreement between replicate ChIP experiments (IDR ≤ 0.05). We also report any genes or repeat annotations that overlap the MACS peaks. Note that there is no general enrichment in *G2/Jockey-3* outside of the centromeric islands.

**Table S16. Chromatin status assignments for contigs.** We assigned contigs from the assembly to a chromosome and a chromatin status (heterochromatin/euchromatin, etc. based on Riddle et al. 2011,2012; see supplemental methods). Blank cells indicate that a region could not be assigned.

**Table S17. Overlap between normal and CENP-A overexpression S2 cells.** We compared the MACS peaks shared between ‘normal’ S2 (this study) and S2 with CENP-A overexpression using the IDR test. Some non-centromeric regions should have more CENP-A enrichment after CENP-A overexpression, however only four peaks have IDR ≤ 0.05. None of these peaks have *G2/Jockey-3*.

11. Kofler R, Betancourt AJ, Schlotterer C. Sequencing of pooled DNA samples (Pool-Seq) uncovers complex dynamics of transposable element insertions in Drosophila melanogaster.

PLoS Genet. 2012;8(1):e1002487. doi: 10.1371/journal.pgen.1002487. PubMed PMID: 22291611; PubMed Central PMCID: PMCPMC3266889.
