## Supplemental figures for "Islands of retroelements are the major components of *Drosophila* centromeres"

### Figure S1

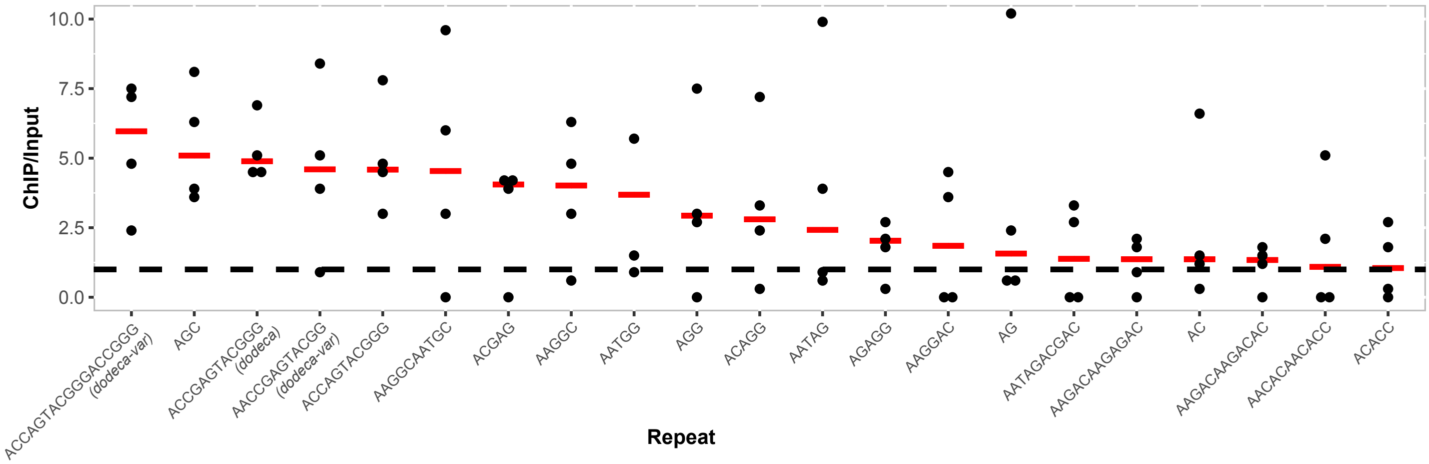


**Figure S1. Enrichment of simple tandem repeats in CENP-A ChIP-seq across four replicates.** Plot of normalized CENP-A/Input for simple tandem repeats for each ChIP-seq replicate, sorted by median (red lines). Shown are only the simple tandem repeats with median CENP-A/Input > 1 in all four CENP-A ChIP replicates (see details in Table S1). The simple tandem repeats with less than 10 counts of input reads in any one replicate are not shown.

### Figure S2

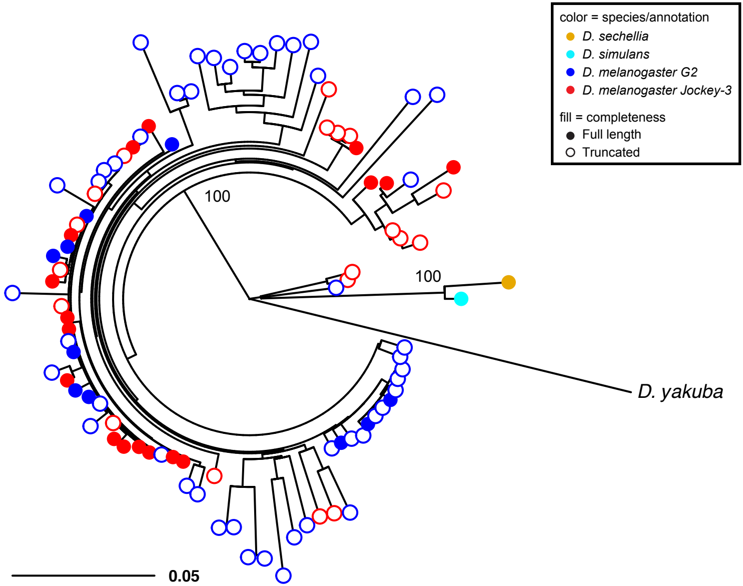


**Figure S2. *G2* and *Jockey-3* correspond to the same Non-LTR retroelement.** A maximum likelihood phylogenetic tree showing the relationship between *G2* and *Jockey-3* sequences in *D. melanogaster* genome and closely related species in the simulans clade (*D. simulans* and *D. sechellia*) and *D. yakuba*. In *D. melanogaster*, *G2* and *Jockey-3* are interleaved across the phylogeny, and thus likely correspond to the same repeat type. We therefore refer to these elements collectively as *G2/Jockey-3* throughout the manuscript.

### Figure S3

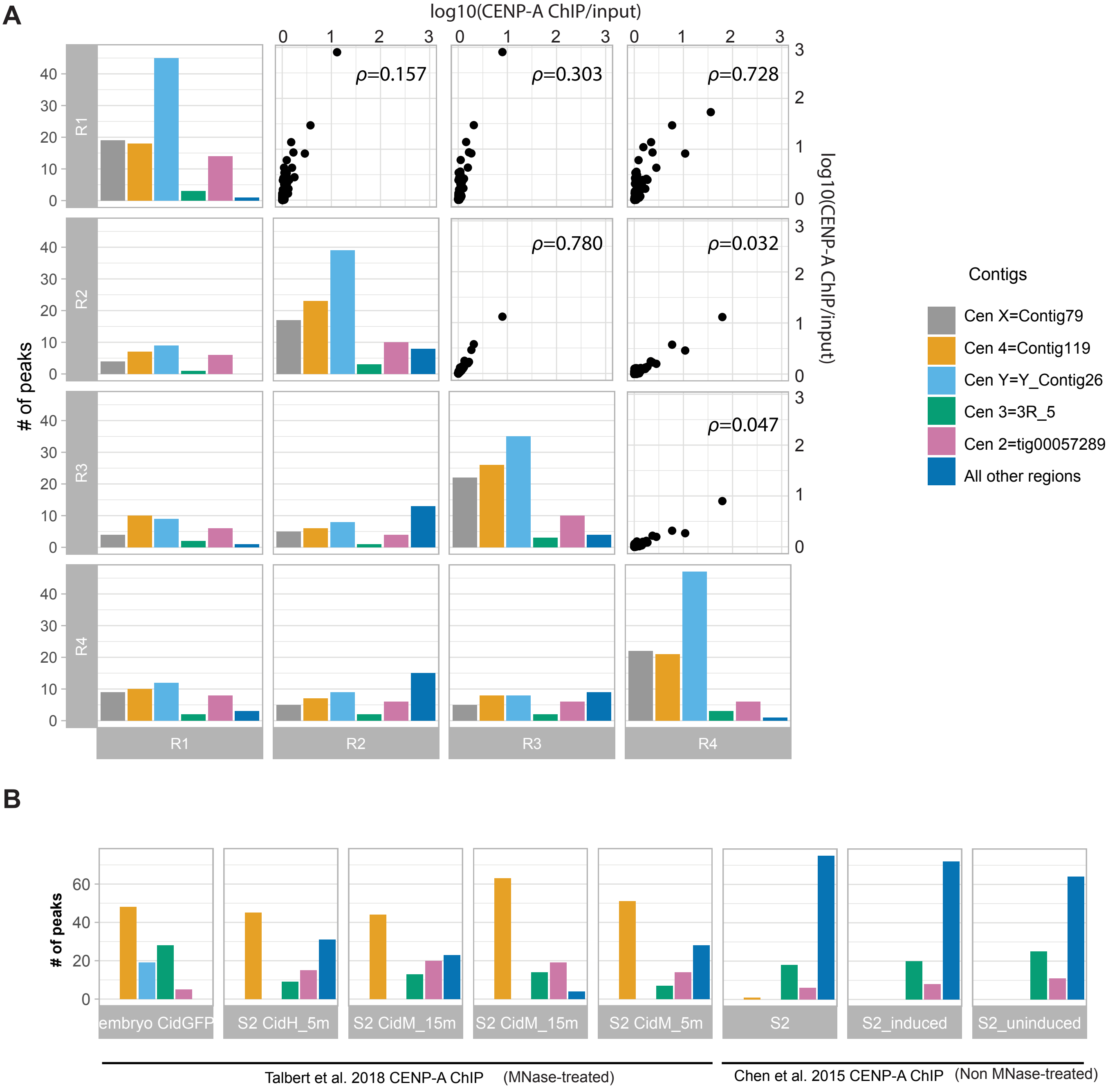


**Figure S3. Reproducibility of CENP-A ChIP enrichment among replicates in embryos and S2 cells.** Locations of the top 100 strongest peaks for each ChIP experiment. **A)** Plot of the location of top 100 strongest peaks for each ChIP experiment on the diagonal (see details in Table S4). For the four replicate ChIP experiment in our OreR embryos, we examined the reproducibility of our experiments by first applying the IDR (Irreproducible Discovery Rate) test and only keeping peaks with IDR≤0.05. The number of these peaks are plotted below the diagonal. Between Replicates 2 and 3, we found a total of 16,870 overlapping peaks, but 16,833 were weakly enriched relative to the overlapping peaks between other datasets because they are technical repeats with a shared library bias (Accel, see Supplemental Methods). We therefore only report the 37 strongest peaks (the average peak number of other comparisons between replicates). The IDR dataset comparisons are in Table S5. We show the correlation between the CENP-A ChIP replicates above the diagonal. Plotted are the signal strength after IDR tests (normalized ChIP over input ratio from 1-1,000 on a log10 scale) with Spearman's rho. The five contigs with the most consistent peaks within and among replicates correspond to the five centromeric candidates. **B)** Plot of ChIP-seq data from S2 cells (this paper, Talbert et al. 2018, and Chen et al. 2015) and an independent embryo CID-GFP ChIP-seq dataset (see details in Table S4; Talbert et al. 2018; 5m and 15m represent different MNase treatments). The centromeric contigs are also CENP-A enriched in these independent datasets, with the exception of the X chromosome centromere contig. S2 cells lack a Y and are therefore not expected to have peaks on the Y candidate centromere contig.

### Figure S4

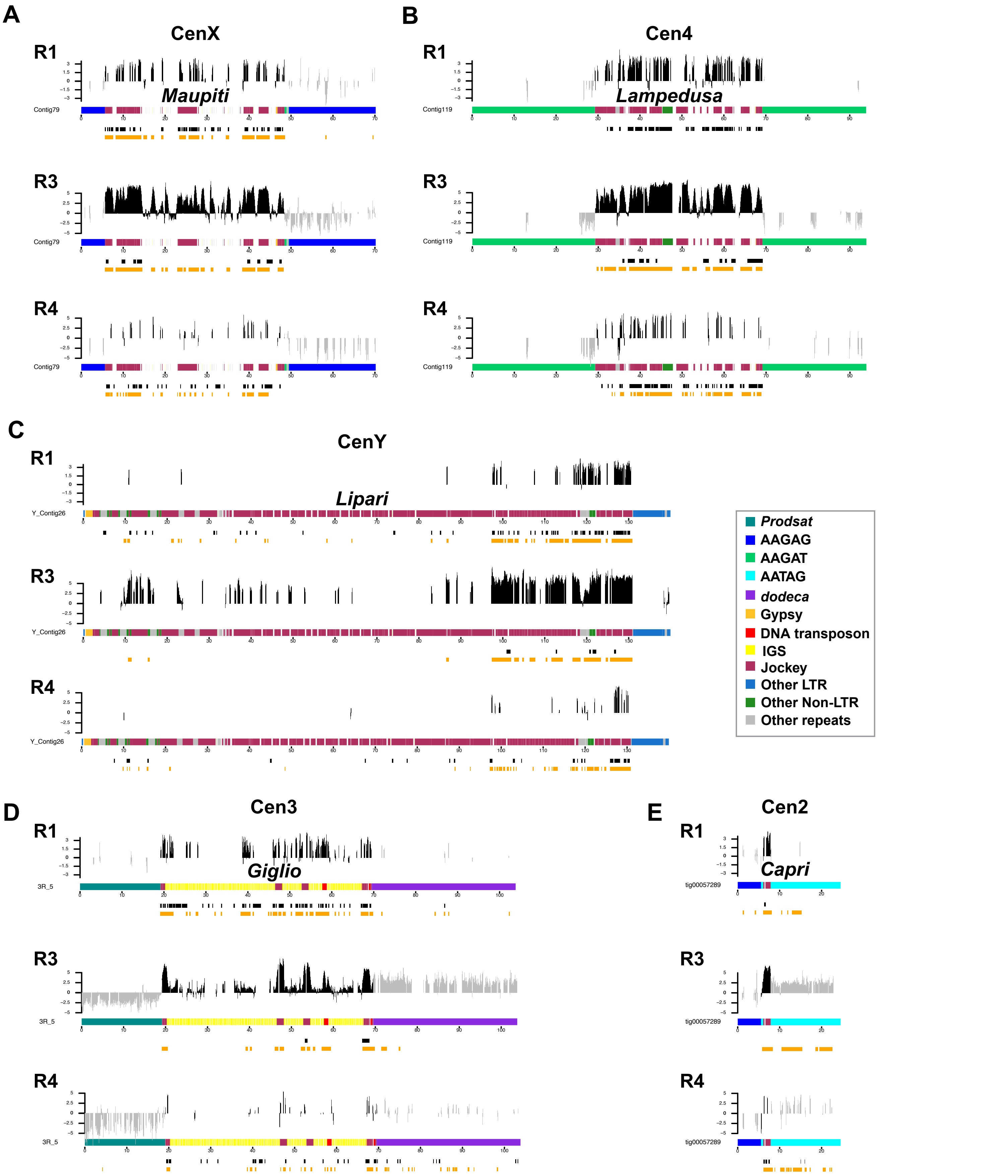


**Figure S4. CENP-A occupies DNA sequences within putative centromere contigs.** Organization of each CENP-A enriched island corresponding to centromere candidates: **A)** X centromere, **B)** centromere 4; **C)** Y centromere; **D)** centromere 3; **E)** centromere 2. Different repeat families are color coded (see legend; note that *Jockey* elements are shown in one color even though they are distinct elements). The normalized CENP-A enrichment over input (plotted on a log scale) is shown for three replicates (Replicate 2 is in Fig. 2) colored in gray for simple repeats and black for complex island sequences. While the mapping quality scores are high in simple repeat regions, we do not use these data to make inferences about CENP-A distribution (see text for details). The coordinates of the significantly CENP-A-enriched ChIPtigs mapped to these contigs (black) and the predicted ChIP peaks (orange) are shown below each plot.

### Figure S5

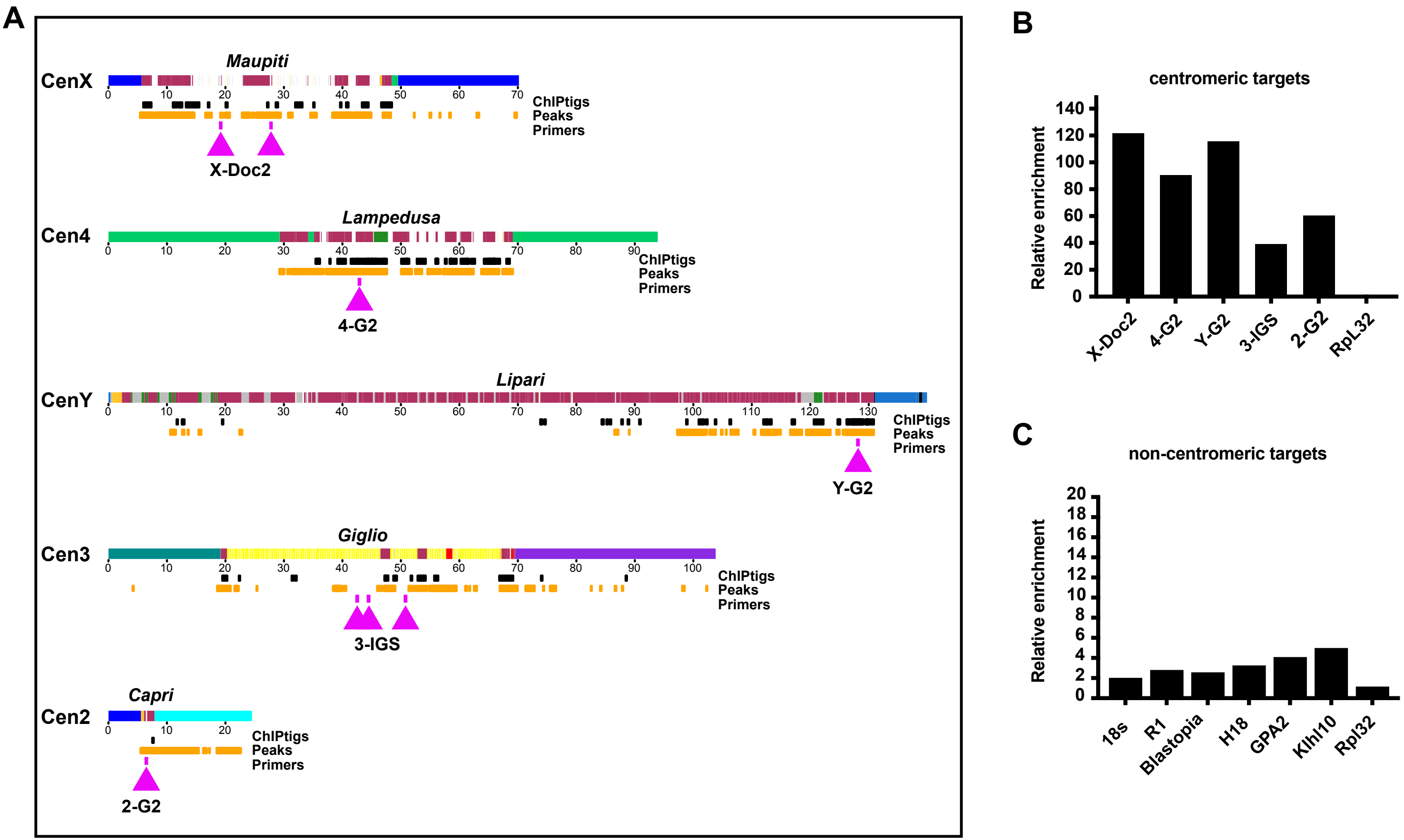


**Figure S5. ChIP-qPCR validation of CENP-A-enriched regions. A)** Diagram showing putative centromere contigs showing the locations of CENP-A ChIPtigs in black and CENP-A MACS peaks in orange as in Figure 2. Locations of contig-specific qPCR primer binding sites are shown by magenta arrows. **B)** Graph showing our ChIP-qPCR results using these primers. The enrichment is calculated relative to the input and is normalized by the *RpL32* promoter region as a noncentromeric control. **C)** Graph showing our ChIP-qPCR results using primers targeting other regions that showed CENP-A enrichment, but that were not in our contigs. Again, the enrichment is calculated relative to the input and is normalized by *RpL32* promoter as a noncentromeric control. We did not observe a robust CENP-A enrichment at these sites.

### Figure S6

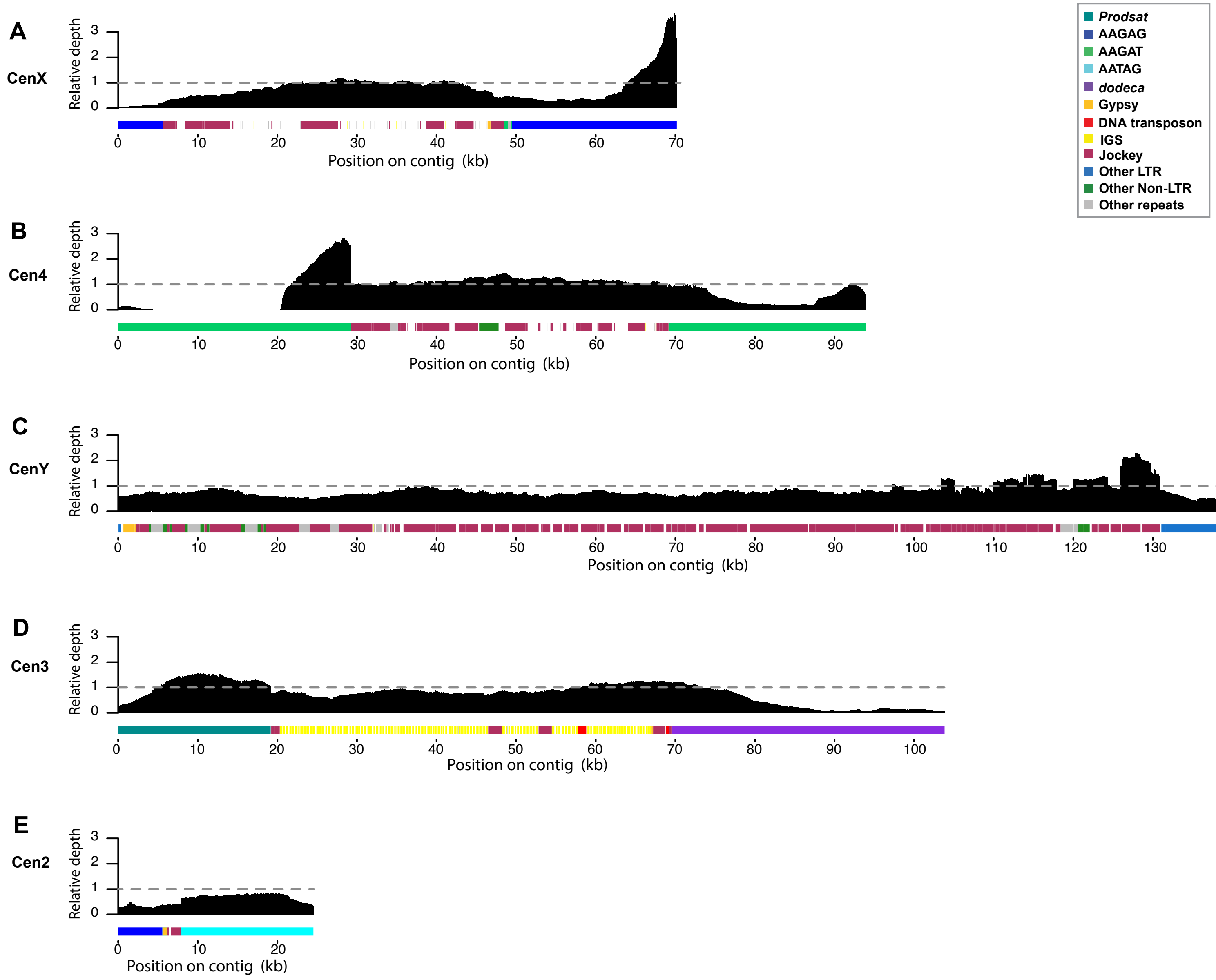


**Figure S6. Relative depth of Pacbio reads across centromeric contigs.** Pacbio reads were mapped to the genome using Minimap (v 2.11) and the parameter "-ax map-pb.". Shown are **A)** X centromere, **B)** centromere 4; **C)** Y centromere; **D)** centromere 3; **E)** centromere 2. The depth of only the high-quality mapped reads (mapped Q ≥ 30) was estimated for each position and normalized by the median depth of other genomic regions (98.32x for autosomes and 49.16x for sex chromosomes) to get relative depth. The relative depth of the TE-rich islands are close to 1, whereas the depth of the flanking simple satellites are uneven with some regions >1 and some <1. We therefore exclude simple repeats from any assembly-based analyses and color these regions gray in Fig. 2 and Fig. S4 to indicate that caution should be used in interpreting these regions of the assembly.

### Figure S7

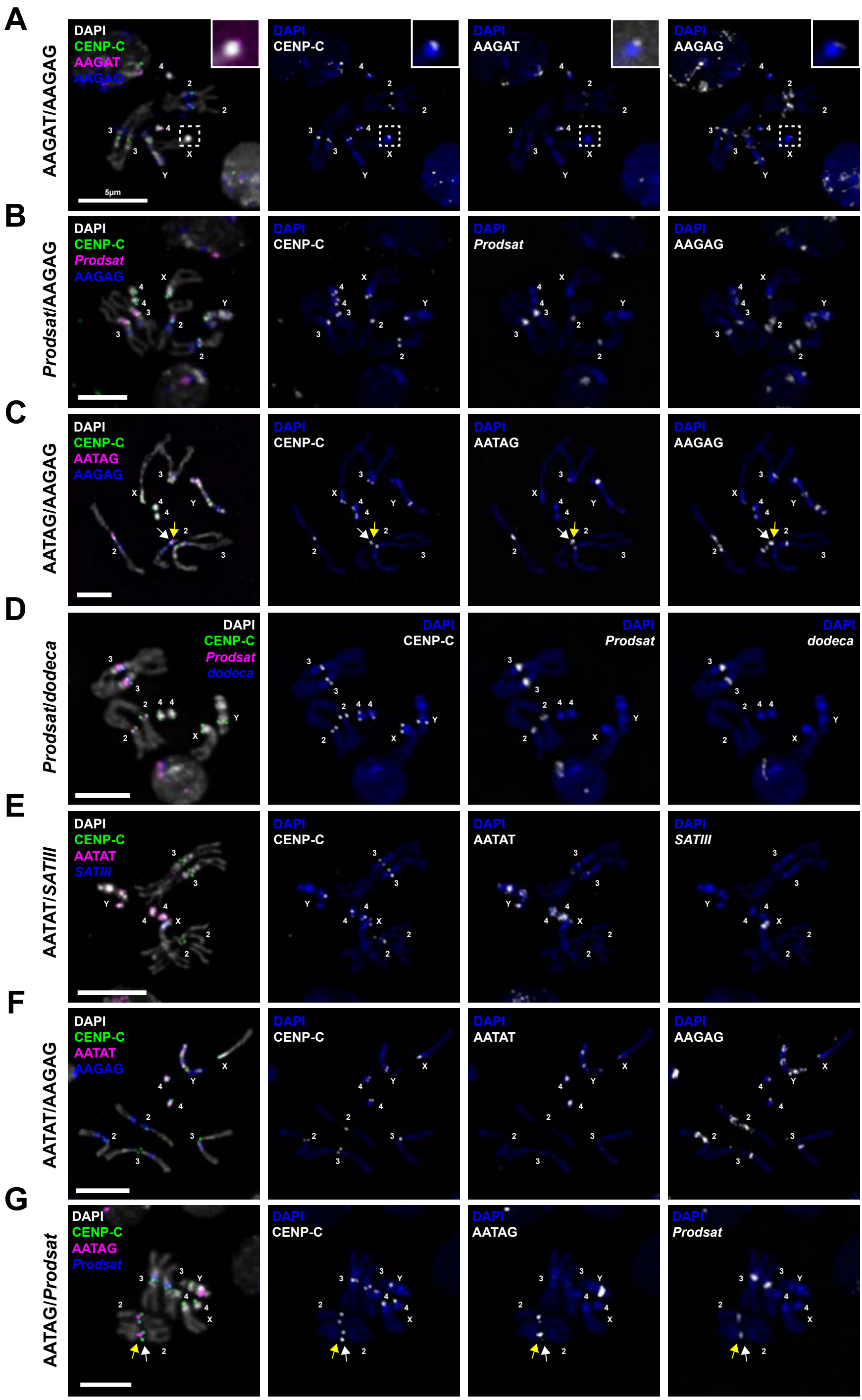


**Figure S7. Satellite FISH on iso-1 larval brain mitotic spreads.** IF/FISH using an anti-CENP-C antibody (green) and satellite FISH probes in the following combinations: **A)** AAGAT (magenta), and AAGAG (blue), with a high contrast inset of AAGAT on the X chromosome; **B)** *Prodsat* (magenta) and AAGAG (blue); **C)** AATAG (magenta) and AAGAG (blue), with AATAG blocks identified by white (small block) and yellow (large block) arrows; **D)** *Prodsat* (magenta) and *dodeca* (blue); **E)** AATAT (magenta) and *SATIII* (blue); **F)** AATAT (magenta) and AAGAG (blue); **G)** AATAG (magenta) and *Prodsat* (blue), with AATAG blocks identified by white (small block) and yellow (large block) arrows. DAPI is shown in gray. Bar 5µm

### Figure S8

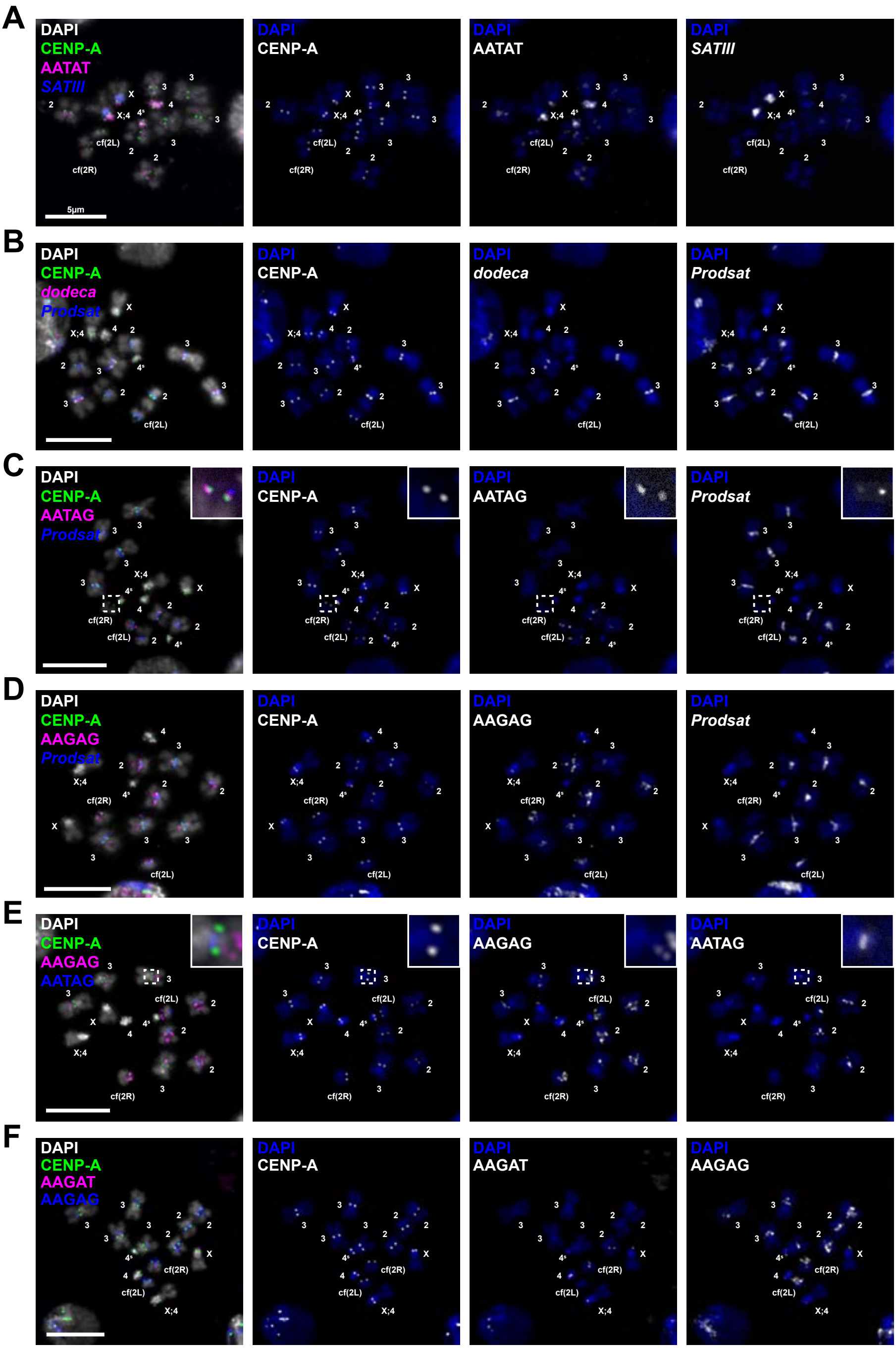


**Figure S8. Satellite FISH on S2 cell mitotic spreads.** IF-FISH using an anti-CENP-A antibody (green) and satellite FISH probes in the following combinations: **A)** AATAT (magenta) and *SATIII* (blue); **B)** *dodeca* (magenta) and *Prodsat* (blue); **C)** AATAG (magenta) and *Prodsat* (blue), with a high contrast inset of AATAG and *Prodsat* on cf(2R); **D)** AAGAG (magenta) and *Prodsat* (blue); **E)** AAGAG (magenta) and AATAG (blue), with a high contrast inset of AATAG on chromosome 3; **F)** AAGAT (magenta) and AAGAG (blue). DAPI is shown in gray. Bar 5 µm.

### Figure S9

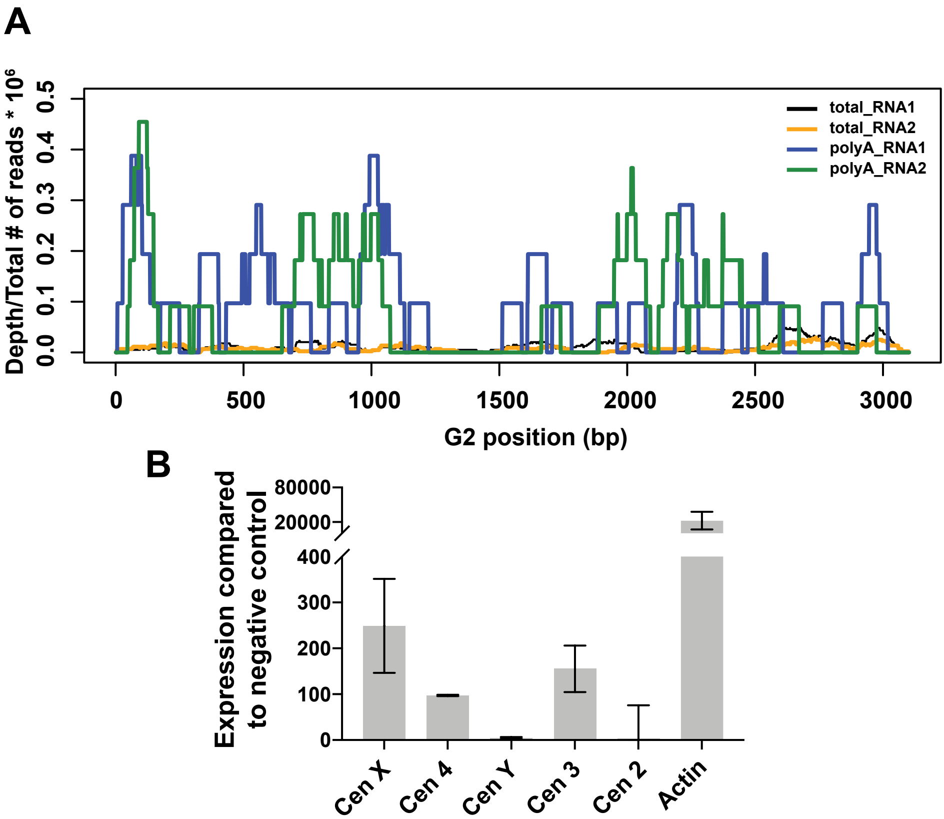


**Figure S9. Transcription of *G2/Jockey-3* elements.** A) Shown is the plot of the normalized reads depth from uniquely mapped reads (mapping quality ≥ 10) across the *G2/Jockey-3* consensus element obtained from mapping total and poly-A RNA-seq data from testes (Laktionov et al. 2018; Gerstein et al. 2014) to our repeat library. B) Quantitative RT-PCR analysis of total RNA extracted from three independent overnight embryo collections. Expression levels were compared to the negative control gene Mst84Da (testis-specific). Cen X, 4 and 3, but not Y and 2, show low levels of transcription compared to the housekeeping gene Actin. Although the primers (Table S7) are specific for each centromere, the primer sets could amplify *G2/Jockey-3* copies not included in our assembly. Error bars=SD.

### Figure S10

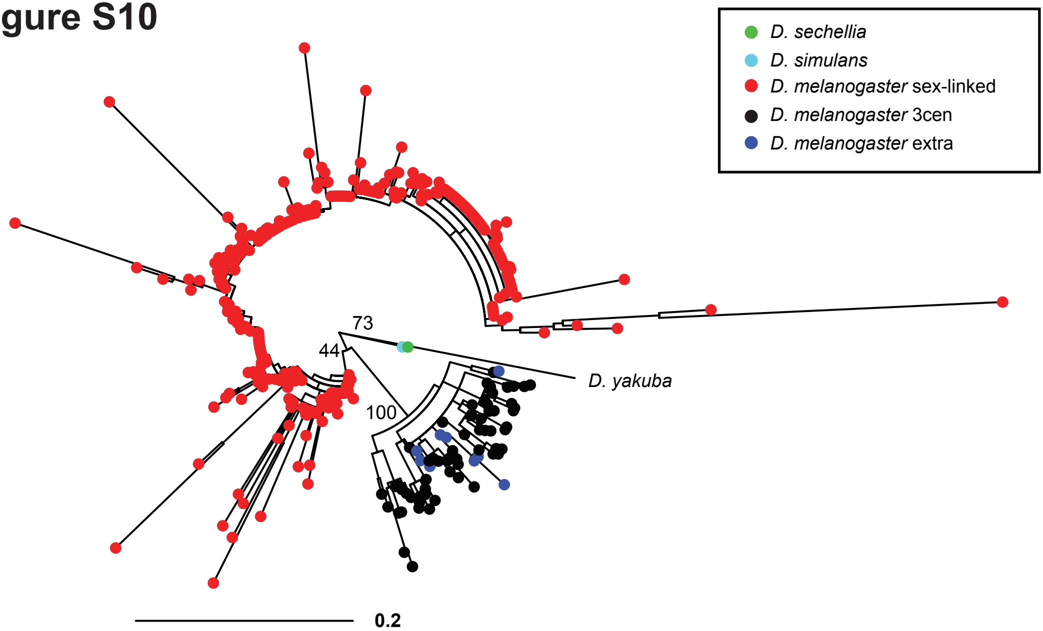


**Figure S10. Relationship of IGS in *D. melanogaster* and closely related species of the simulans clade (*D. simulans* and *D. sechellia*) and *D. yakuba*.** Maximum likelihood phylogenetic tree of all individual IGS sequences found in the *D. melanogaster* genome with related outgroups. Node support is only shown for key nodes in the tree (complete tree is in supplemental file 14). All centromeric IGS sequences appear to have a single origin: they duplicated from sex-linked IGS interspersed at the rDNA loci at some time near the divergence of the simulans clade and *D. melanogaster*. IGS repeats in blue (extra) are similar to the IGS at the cen 3 island, but are on small contigs of unknown origin, one of which is moderately enriched in CENP-A. We refer to all black and blue IGS repeats as $IGS^{3cen}$.

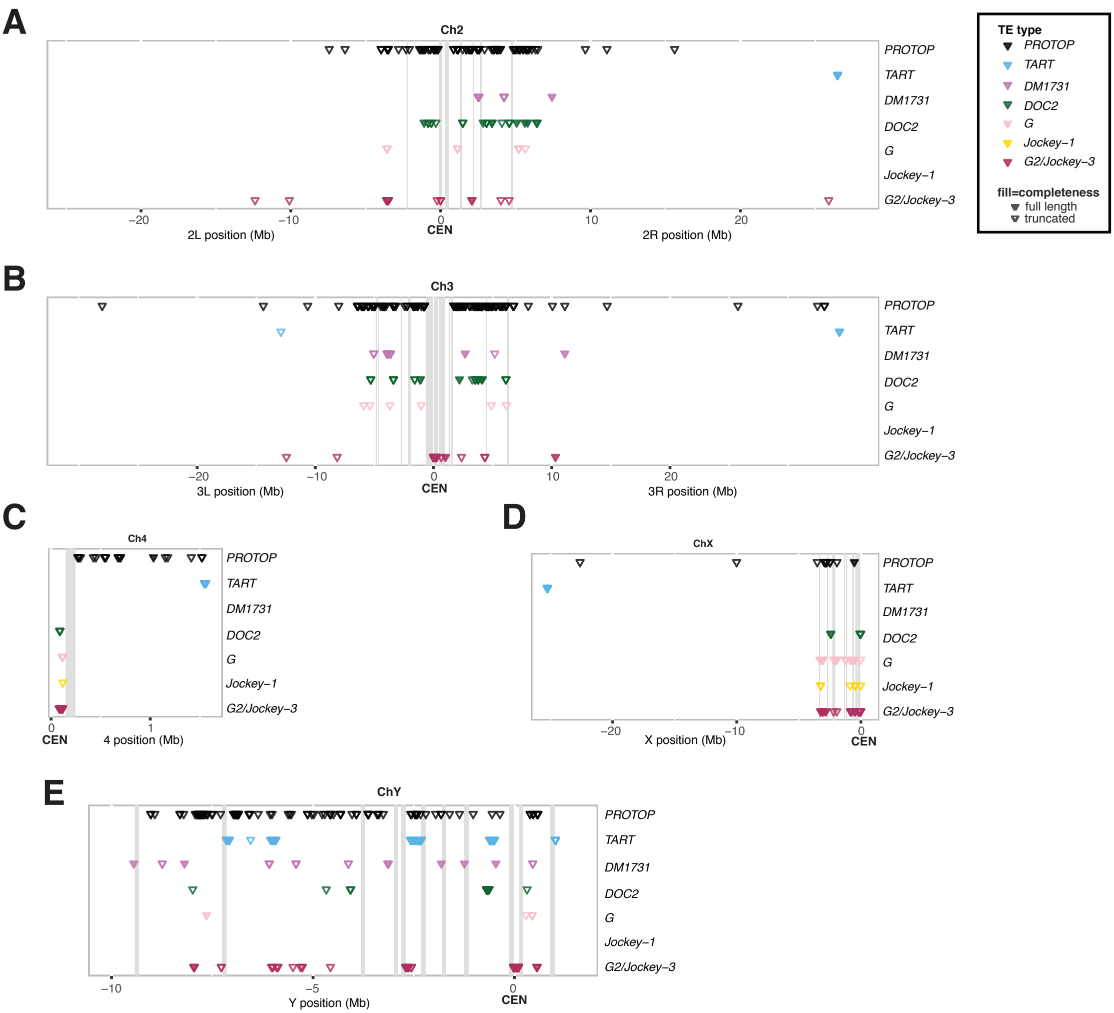


**Figure S11. Genomic TE distribution across chromosomes**. Distribution of TEs (represented by different colors) along the following chromosomes: **A)** chromosome 2; **B)** chromosome 3; **C)** chromosome 4; **D)** chromosome X; and **E)** chromosome Y. Contigs from each chromosome were concatenated in order with an arbitrary insertion of 100 kb of ‘N’. Distances along the x-axis are approximate. The order and orientation of the Y chromosome contigs is based on gene order (see Chang and Larracuente 2018). Each triangle corresponds to one TE, where filled shapes indicate full length TEs and open shapes indicate truncated TEs. The vertical grey bars represent the arbitrary 100kb window inserted between contigs, indicating where we have gaps in our assembly. The centromere positions are set to 0 for each chromosome. Chromosomes are not drawn to scale (chromosome 4 and Y are enlarged). We show the genomic distribution of a sample of TEs enriched in CENP-A according to our ChIPseq analysis (all except *PROTOP*). *PROTOP* are DNA transposons that have not been recently active and their distribution is primarily in heterochromatin. TART elements are non-LTR retroelements highly enriched at telomeres, and are also moderately CENP-A enriched. *DM1731* is a retroelement moderately enriched for CENP-A, but not enriched in the centromere islands. *DOC2*, *G*, *Jockey-1* and *G2/Jockey-3* are CENP-A enriched non-LTR retroelements abundant in the centromere islands (see Table S2 and Table S9).

### Figure S12

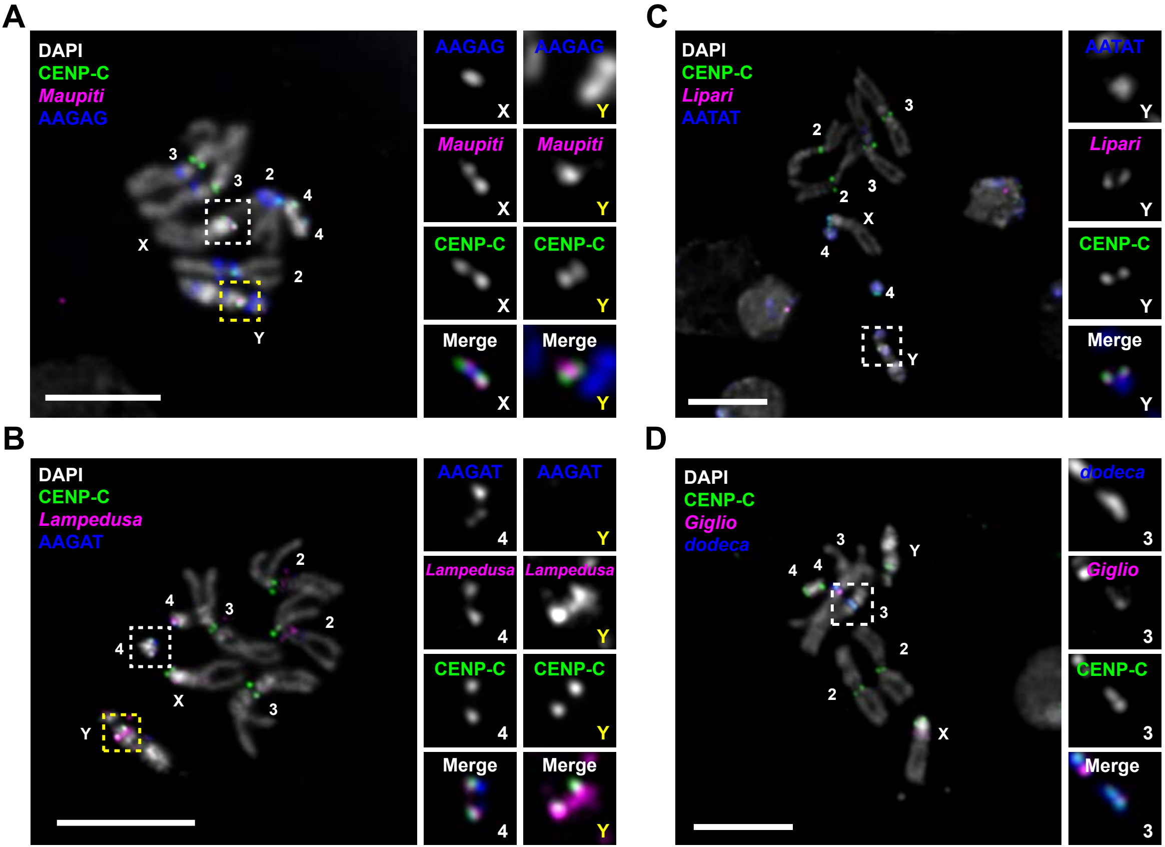


**Figure S12. Oligopaint FISH on larval brain mitotic spreads from iso-1 flies.** IF-FISH using an antibody for CENP-C (green), centromere Oligopaint FISH probes (magenta), and FISH probes for centromeric satellites (blue) in the following combinations: **A)** *Maupiti* (X; magenta) and AAGAG (blue); **B)** *Lampedusa* (4; magenta) and AAGAT (blue); **C)** *Lipari* (Y; magenta) and AATAT (blue); **D)** *Giglio* (3; magenta) and *dodeca* (blue). White boxes show the separate signals at the targeted centromeres. Yellow boxes show centromeric hybridizations at other centromeres. DAPI is shown in gray. Bar 5μm.

### Figure S13

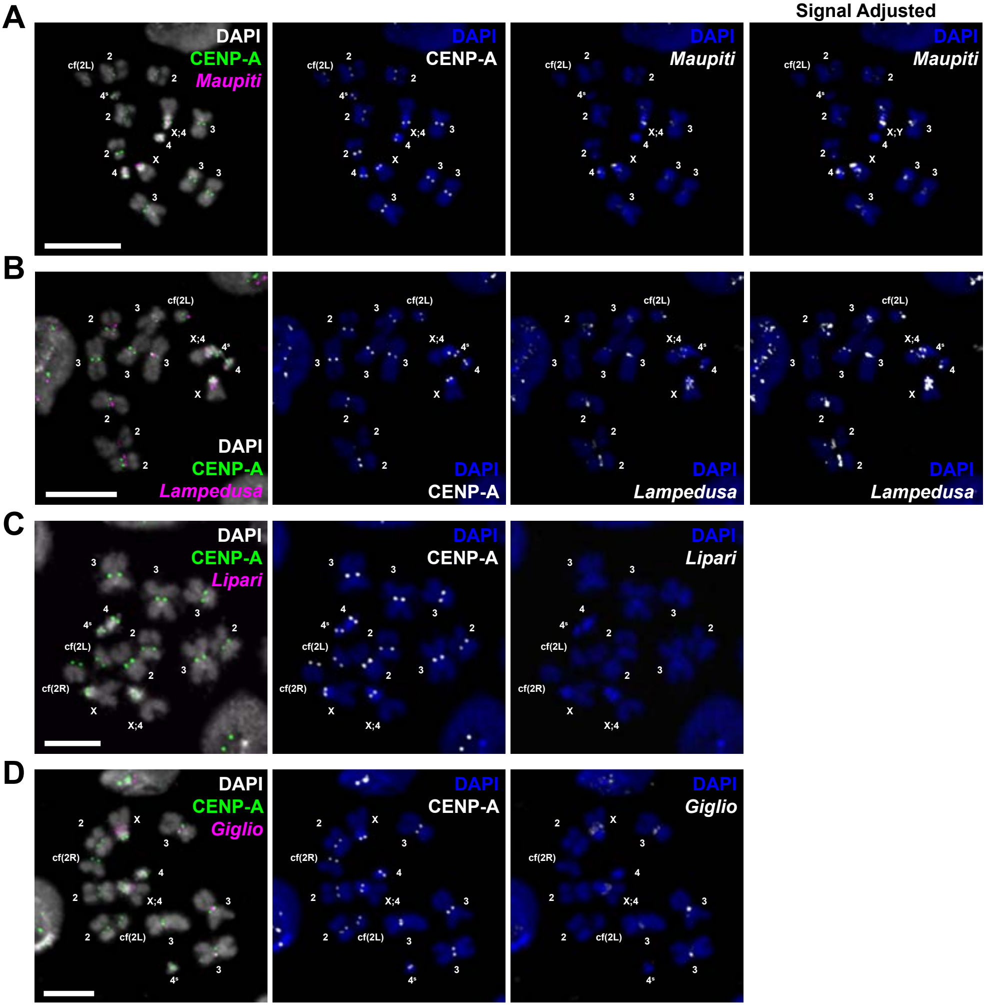


**Figure S13. Oligopaint FISH on S2 cell mitotic spreads.** IF/FISH using an antibody for CENP-A (green) and centromere oligopaint FISH probes designed to target centromere contigs (magenta). **A)** *Maupiti* (X); **B)** *Lampedusa* (4); **C)** *Lipari* (Y); **D)** *Giglio* (3). The "Signal Adjusted" panels for **A** and **B** show high contrast Oligopaint hybridization for visualization of weak foci. Bar 5μm.

#### Figure S14

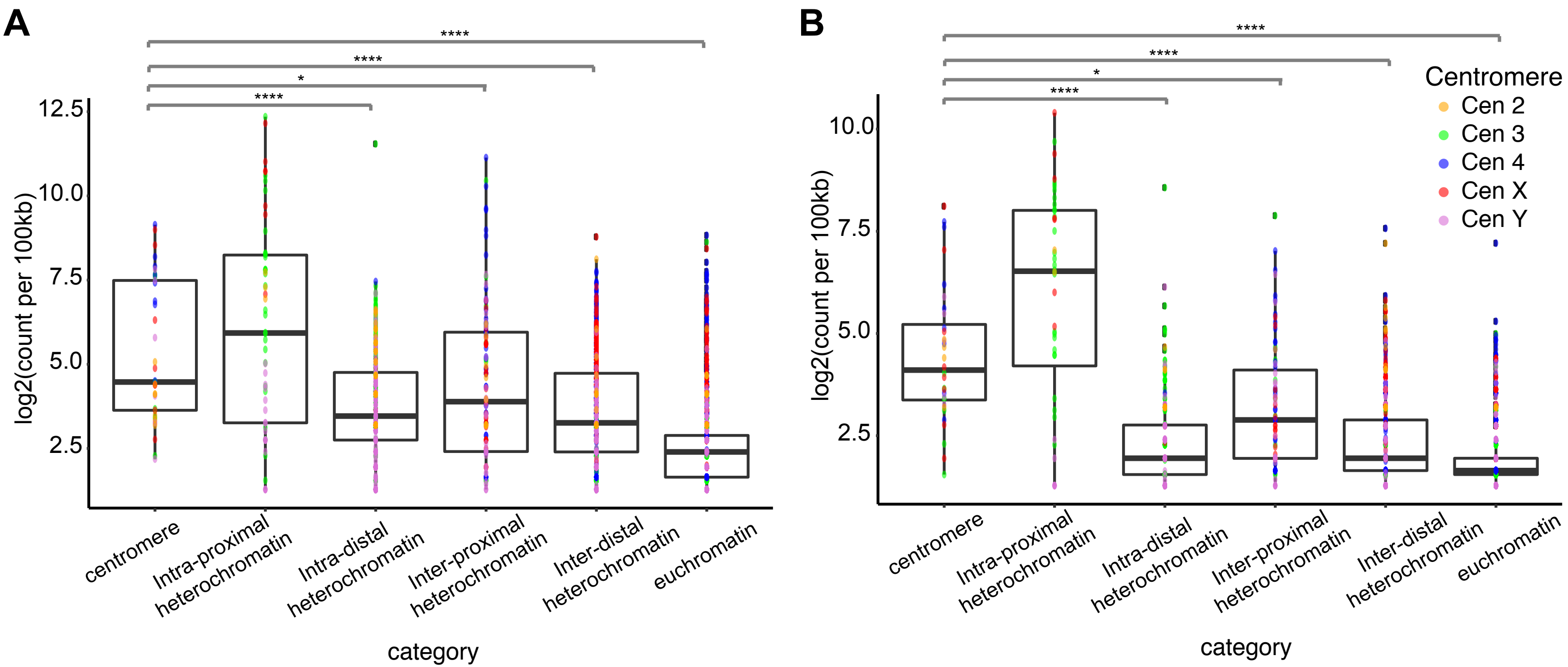


**Figure S14. Quantification of interactions between centromeres and different genomic regions by Hi-C.** Plots showing intra- and inter-chromosomal interactions between regions in Hi-C data from: A) stage 16 embryos (end of embryogenesis) and B) embryonic cycles 1–8 (before zygotic genome activation; data from Ogiyama et al. 2018). The different colors indicate interactions with individual centromeres of all chromosomes. Centromere-centromere interactions are significantly more frequent than interactions between centromeres and distal heterochromatin, inter distal heterochromatin and euchromatin, and marginally more significant than centromere-inter proximal heterochromatin interactions. **** adjusted $P<0.0001$; * adjusted $P<0.02$, Pairwise Wilcoxon rank sum test with false discovery rate (FDR) correction; Kruskal-Wallis test by ranks with Dunn's test for post-hoc analysis.

### Figure S15

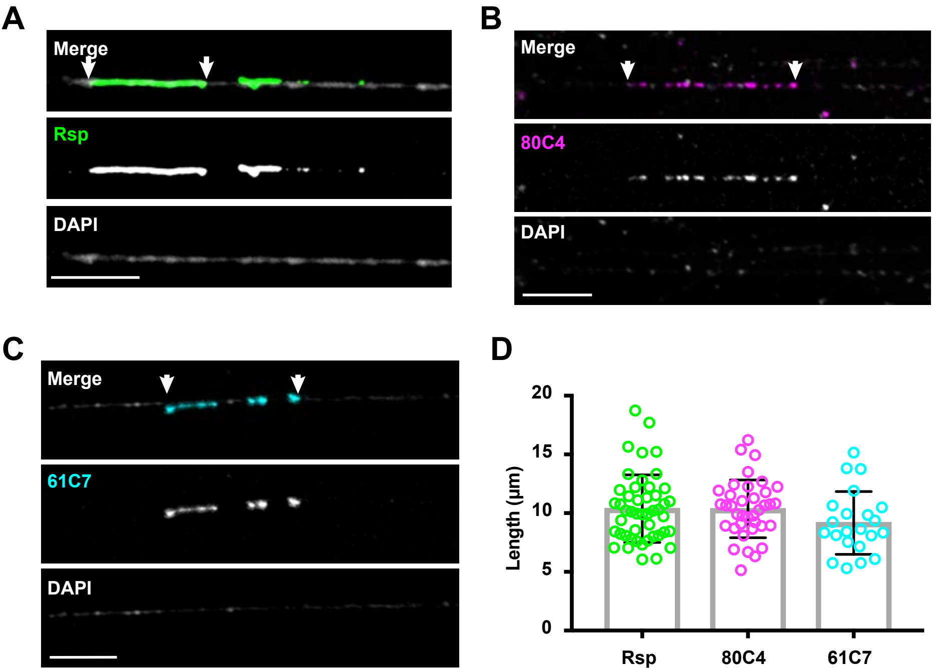


**Figure S15. Calibration of extended chromatin fiber stretching.** Stretched chromatin fibers from female 3rd instar larval brain cells using the following probes: **A)** *Rsp* locus (heterochromatic; ~100kb; green); **B)** 100kb Oligopaint for a heterochromatic region on chromosome 3L (80C4; magenta); **C)** 100kb Oligopaint for a euchromatic region ~600kb from the telomere of chromosome 3L (61C7; cyan). Arrows show the region of the fiber that was measured. Bar 5µm. **D)** Scatter plot showing the quantification of fiber lengths. Mean lengths were used to estimate the size in kb (~10kb/1µm). Error bars show the standard deviation. P>0.09 (n.s.) for each pair of measurements compared (two-tailed t-test).

### Figure S16

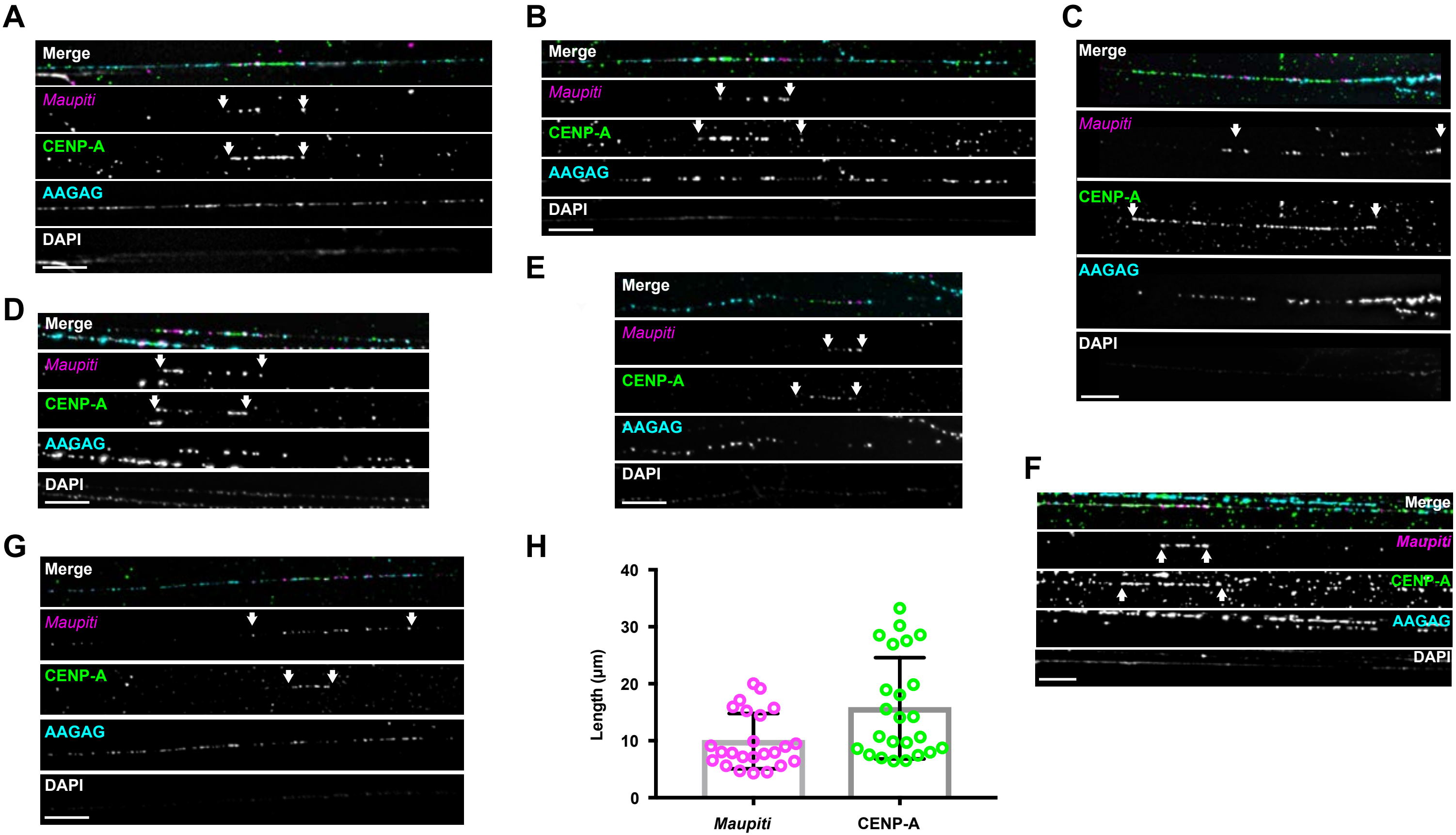


**Figure S16. Organization of the X centromere. (A-G)** Examples of fibers visualized with IF with anti-CENPA antibody (green), FISH with Oligopaints for *Maupiti* (magenta), and a AAGAG probe (cyan) on female 3rd instar larval brain cells. DAPI is shown in gray. CENPA occupies *Maupiti* and the AAGAG satellite. We observed some variation in FISH signals and *Maupiti* and CENP-A domain lengths, likely due to the efficiency of oligopaint binding and variable stretching in this region. Arrows show the region of the fiber that was measured. **H)** Scatter plot showing the quantification of the length of *Maupiti* FISH and CENP-A IF signals. Error bars show the standard deviation. N=24 fibers. Bar 5µm.

### Figure S17

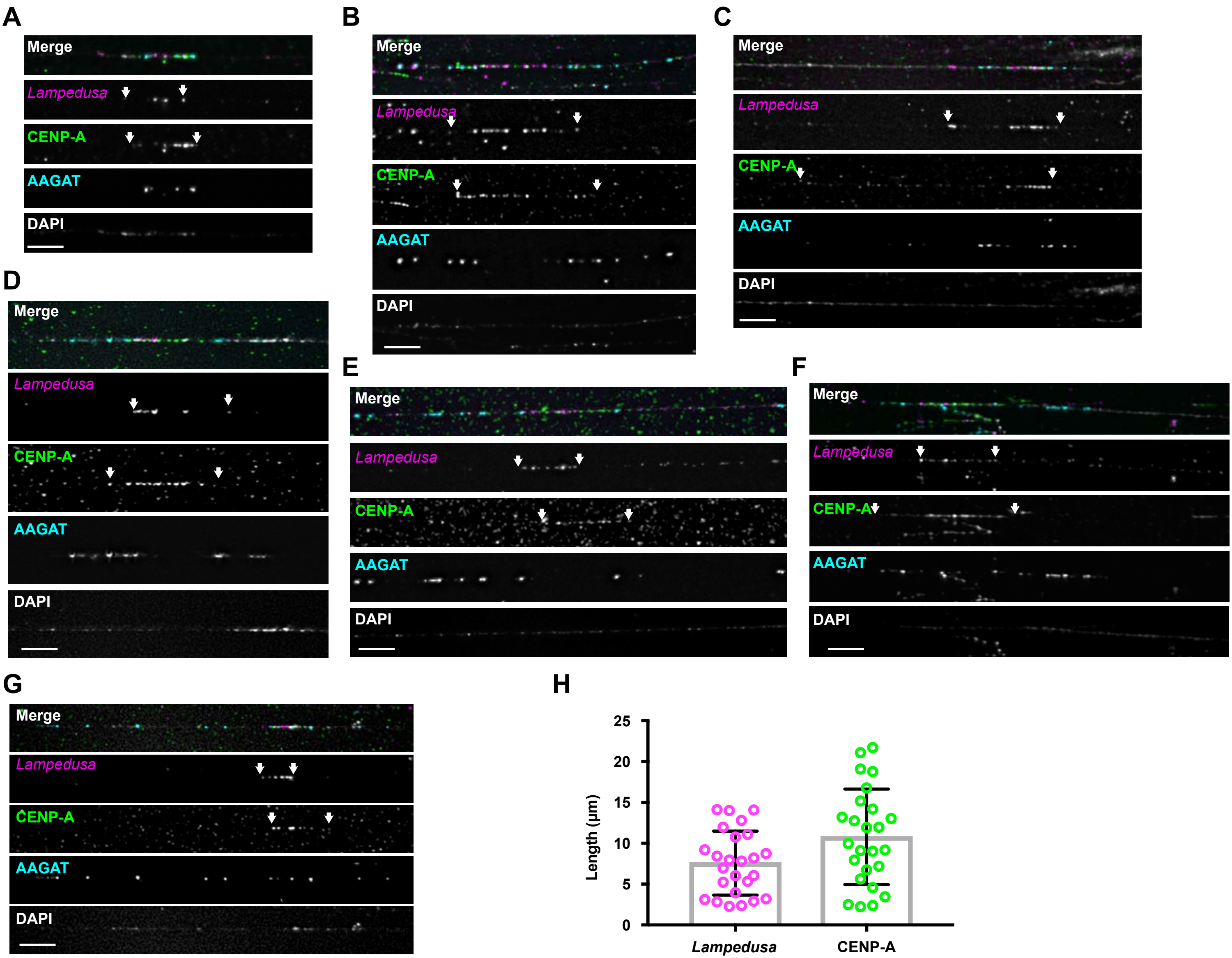


**Figure S17. Organization of centromere 4**. **(A-G)** Examples of fibers visualized by IF with anti-CENPA antibody (green), FISH Oligopaint FISH for *Lampedusa* (magenta), and a AAGAT probe (cyan). DAPI is shown in gray. CENP-A occupies predominantly the island *Lampedusa*. Arrows show the region of the fiber that was measured. **H)** Scatter plot showing the quantification of the length of *Lampedusa* FISH and CENP-A IF signals. Error bars show the standard deviation. N=25 fibers. Bar 5μm.

### Figure S18

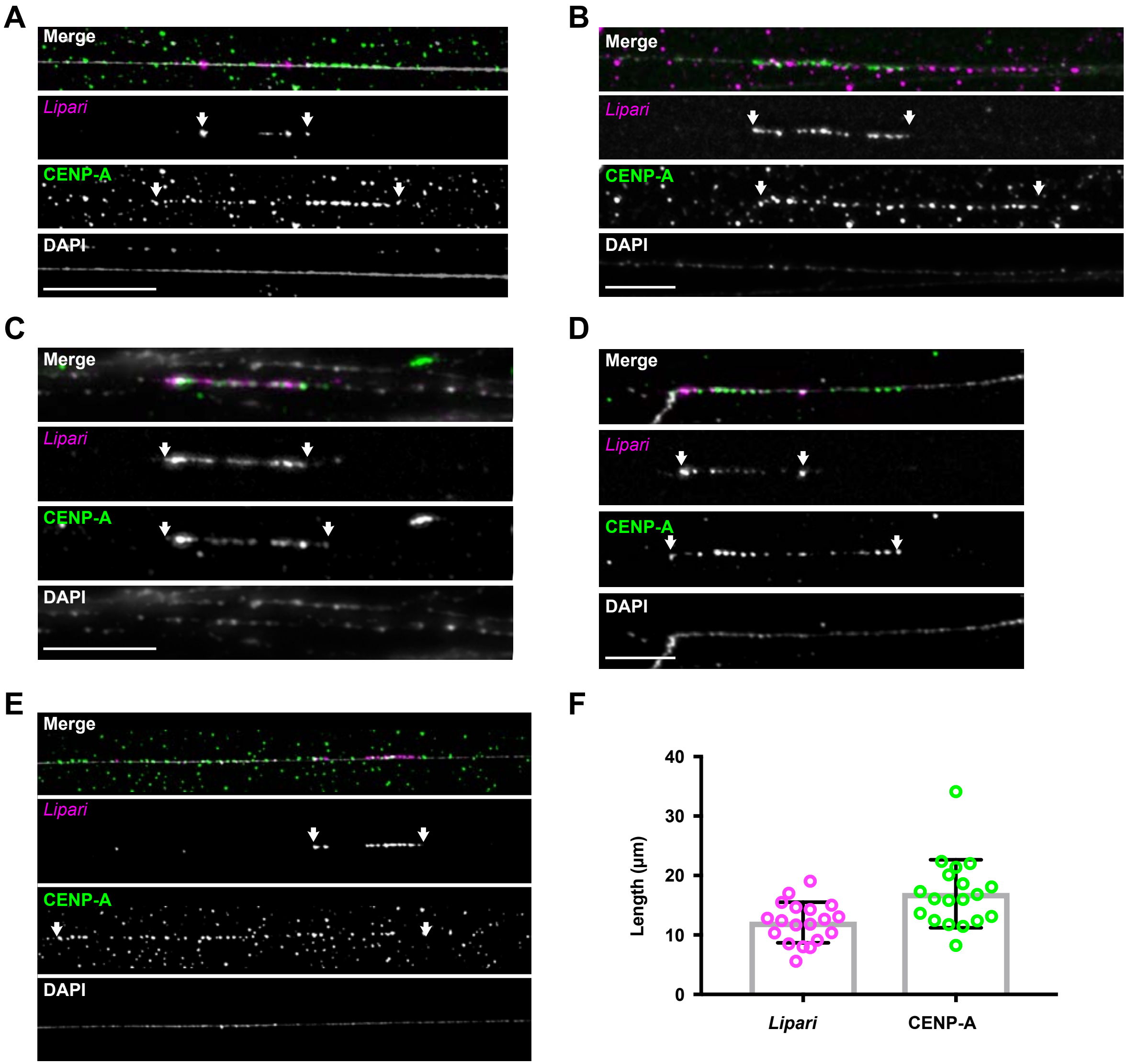


**Figure S18. Organization of the Y centromere. (A-E)** Examples of fibers visualized by IF with anti-CENPA antibody (green), FISH with Oligopaints for *Lipari* (magenta). DAPI is shown in gray. We did not include satellite FISH because no centromeric satellites are known for the Y. Note that the oligopaints only target to part of *Lipari* (see Figure 4). CENP-A is observed occupying sequences beyond the oligopaint region, likely over the remaining part of the island. Arrows show the region of the fiber that was measured. **F)** Scatter plot showing the quantification of the length of *Lipari* FISH and CENP-A IF signals. Error bars show the standard deviation. N=19 fibers. Bar 5μm.

### Figure S19

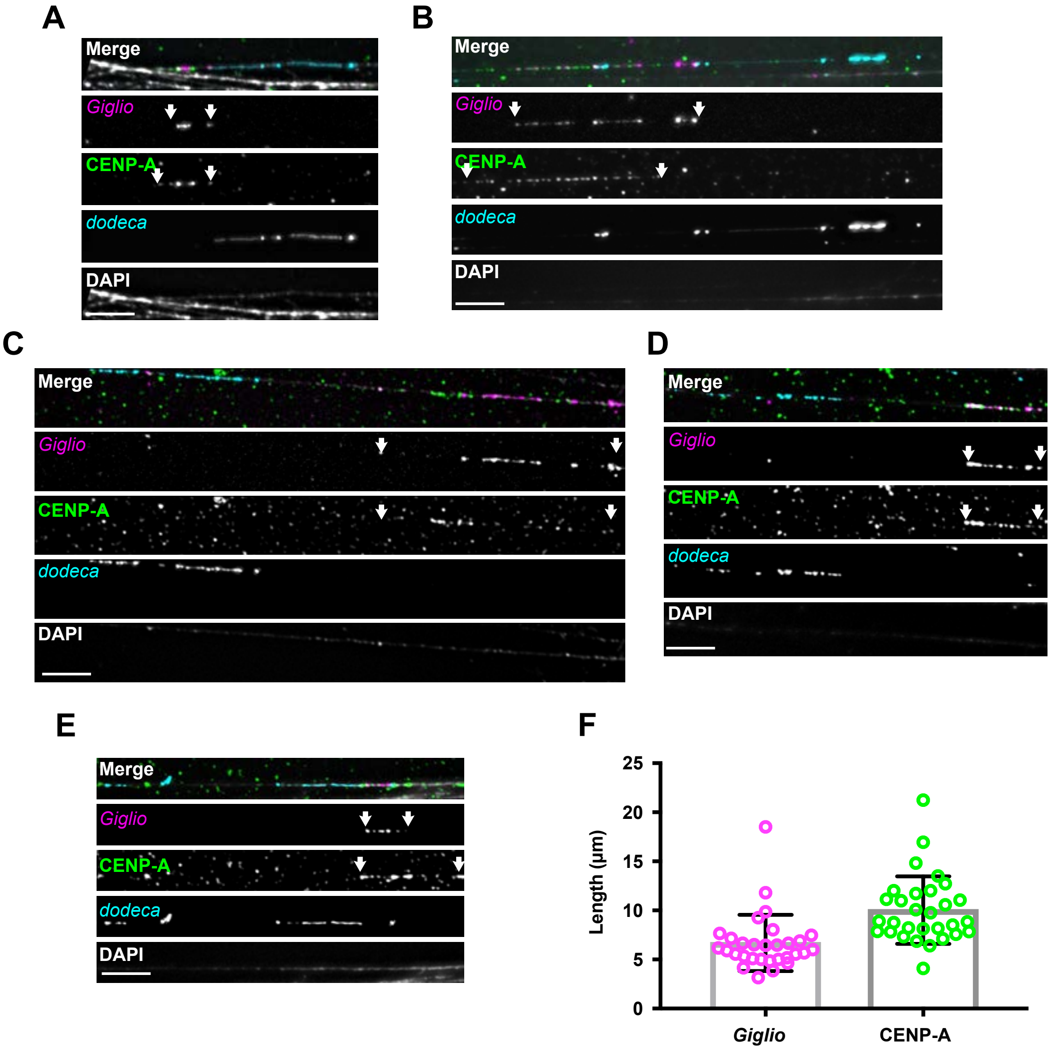


**Figure S19. Organization of centromere 3. A-E)** Examples of fibers visualized by IF with anti-CENP-A antibody (green), FISH with Oligopaints for *Giglio* (magenta), and a probe for the centromere 3 specific *dodeca* satellite (cyan). DAPI is shown in gray. CENP-A occupies primarily *Giglio*, and a small stretch of *dodeca* satellite. Note that the binding of the *dodeca* (an LNA probe) is quite variable between fibers and results in several gaps that could be a result of the higher stringency conditions needed for *Giglio* Oligopaint FISH. Arrows show the region of the fiber that was measured. **F)** Scatter plot showing the quantification of the length of *Giglio* FISH and CENP-A IF signals. Error bars show the standard deviation. N=30 fibers. Bar 5µm.

### Figure S20

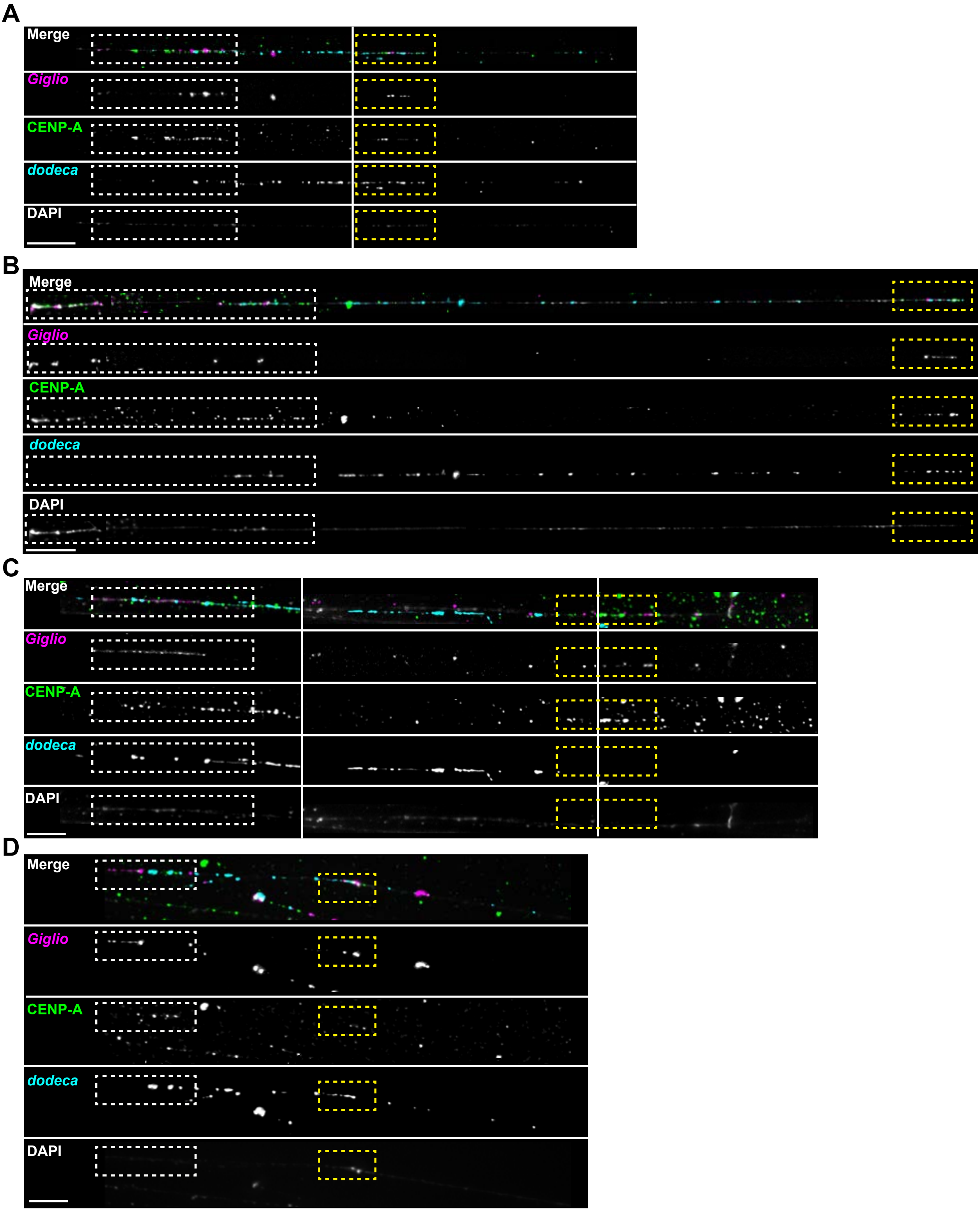


**Figure S20. Tracking of longer centromere 3 fibers reveals a second region containing CENP-A on *dodeca*. (A-D)** Examples of longer fibers tracked along *dodeca* from the experiment in Figure 21, visualized by IF with with anti-CENPA antibody (green), Oligopaint FISH for *Giglio* (magenta), and FISH with *dodeca* probe (cyan). DAPI is shown in gray. Note the presence of *Giglio* signal on the *dodeca* CENP-A region. Multiple, overlapping panels were often acquired to follow an individual fiber. Panels were then cropped and juxtaposed in the figure, with white lines showing the separate images. White boxes show the CENP-A domain on *Giglio,* yellow boxes show the smaller domain on *dodeca*. N=5 (these are rare fibers to find in our preparations due to their length). Bar 5μm.

### Figure S21

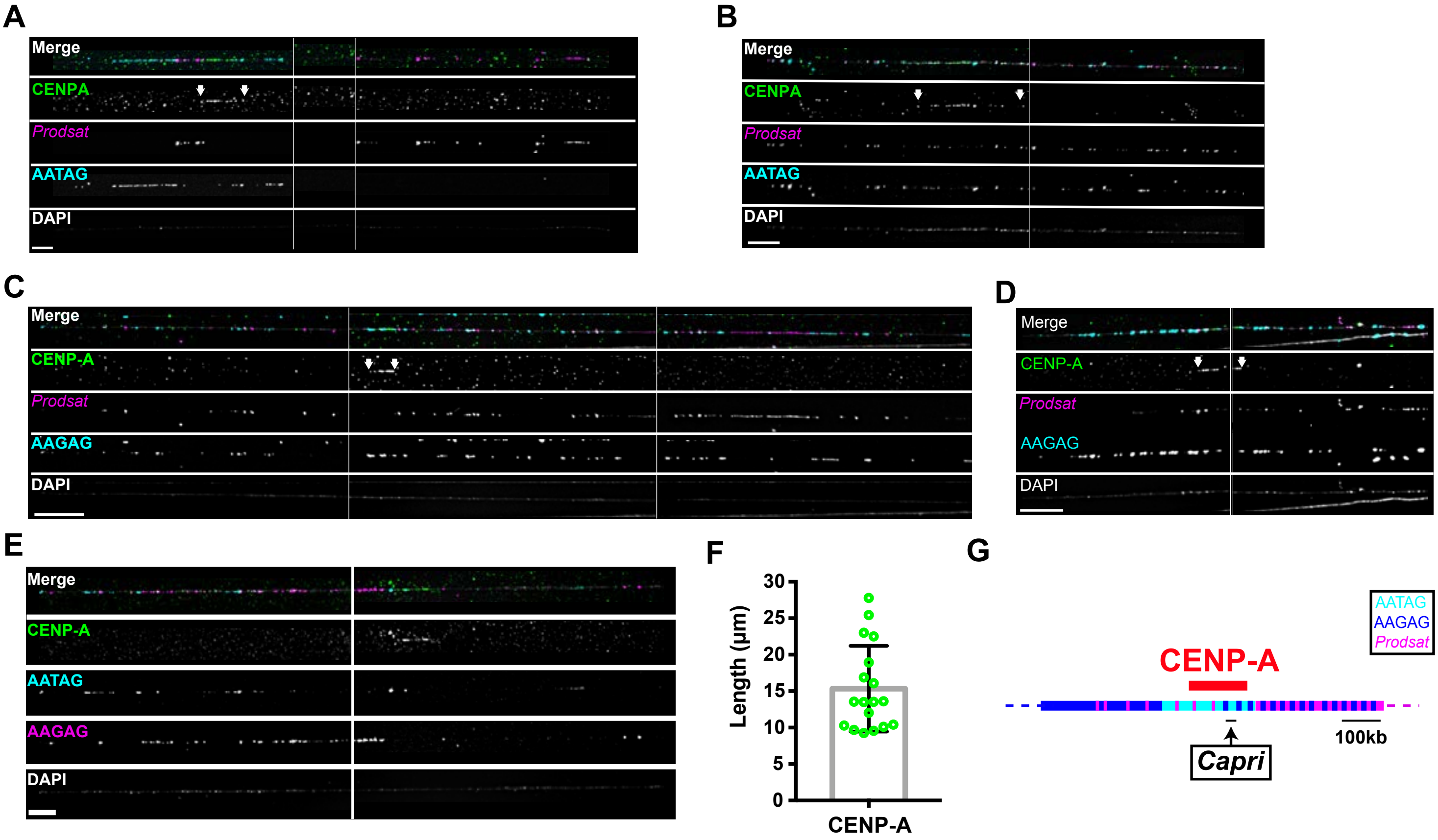


**Figure S21. Organization of the Centromere 2. (A-D)** Examples of fibers visualized with IF with anti-CENP-A antibody (green) and FISH with satellites. DAPI is shown in gray. **(A-B)** Examples of fibers showing colocalization of CENP-A (green) with *Prodsat* (magenta) and AATAG (cyan). **(C-D)** Examples of fibers with AAGAG (cyan) and *Prodsat* (magenta). **E)** Example of fiber with AAGAG (magenta) and AATAG (cyan). We propose that *Capri* is located between flanking blocks of AAGAG and AATAG satellites that reside very close to where the *Prodsat* begins. Arrows show the region that was measured for each fiber. **F)** Scatter plot of CENP-A IF signals lengths. **G)** Model for the organization of centromere 2 showing a possible location of *Capri*. Error bars show the standard deviation. N= 18 fibers. Bar 5µm.
